# graphsim: An R package for simulating gene expression data from graph structures of biological pathways

**DOI:** 10.1101/2020.03.02.972471

**Authors:** S. Thomas Kelly, Michael A. Black

## Abstract

Transcriptomic analysis is used to capture the molecular state of a cell or sample in many biological and medical applications. In addition to identifying alterations in activity at the level of individual genes, understanding changes in the gene networks that regulate fundamental biological mechanisms is also an important objective of molecular analysis. As a result, databases that describe biological pathways are increasingly uesad to assist with the interpretation of results from large-scale genomics studies. Incorporating information from biological pathways and gene regulatory networks into a genomic data analysis is a popular strategy, and there are many methods that provide this functionality for gene expression data. When developing or comparing such methods, it is important to gain an accurate assessment of their performance. Simulation-based validation studies are frequently used for this. This necessitates the use of simulated data that correctly accounts for pathway relationships and correlations. Here we present a versatile statistical framework to simulate correlated gene expression data from biological pathways, by sampling from a multivariate normal distribution derived from a graph structure. This procedure has been released as the `graphsim` R package on CRAN and GitHub (https://github.com/TomKellyGenetics/graphsim) and is compatible with any graph structure that can be described using the `igraph` package. This package allows the simulation of biological pathways from a graph structure based on a statistical model of gene expression.

## Introduction: inference and modelling of biological networks

Network analysis of molecular biological pathways has the potential to lead to new insights into biology and medical genetics (Barabási and Oltvai 2004; Hu, Thomas, and Brunak 2016). Since gene expression profiles capture a consistent signature of the regulatory state of a cell (Perou et al. 2000; Ozsolak and Milos 2011; Svensson, Vento-Tormo, and Teichmann 2018), they can be used to analyse complex molecular states with genome-scale data. However, biological pathways are often analysed in a reductionist paradigm as amorphous sets of genes involved in particular functions, despite the fact that the relationships defined by pathway structure could further inform gene expression analyses. In many cases, the pathway relationships are well-defined, experimentally-validated, and are available in public databases (Croft et al. 2014). As a result, network analysis techniques can play an important role in furthering our understanding of biological pathways and aiding in the interpretation of genomics studies.

Gene networks provide insights into how cells are regulated, by mapping regulatory interactions between target genes and transcription factors, enhancers, and sites of epigenetic marks or chromatin structures (Barabási and Oltvai 2004; Yamaguchi et al. 2007). Inference using these regulatory interactions genomic analysis has the potential to radically expand the range of candidate biological pathways to be further explored, or to improve the accuracy of bioinformatics and functional genomic analysis. A number of methods have been developed to utilise timecourse gene expression data (Yamaguchi et al. 2007; Arner et al. 2015) using gene regulatory modules in state-space models and recursive vector autoregressive models (Hirose et al. 2008; Shimamura et al. 2009). Various approaches to gene regulation and networks at the genome-wide scale have led to novel biological insights (Arner et al. 2015; Komatsu et al. 2013), however, inference of regulatory networks has thus far primarily relied on experimental validation or resampling-based approaches to estimate the likelihood of specific network modules being predicted (Markowetz and Spang 2007; Hawe, Theis, and Heinig 2019).

Simulating datasets that account for pathway structure are of particular interest for benchmarking regulatory network inference techniques and methods being developed for genomics data containing complex biological interactions (Schaffter, Marbach, and Floreano 2011; Saelens et al. 2019). Dynamical models using differential equations have been employed, such as by GeneNetWeaver (Schaffter, Marbach, and Floreano 2011), to generate simulated datasets specifically for benchmarking gene regulatory network inference techniques. There is also renewed interest in modelling biological pathways and simulating data for benchmarking due to the emergence of single-cell genomics technologies and the growing number of bioinformatics techniques developed to use this data (Zappia, Phipson, and Oshlack 2017; Saelens et al. 2019). Packages such as ‘splatter’ (Zappia, Phipson, and Oshlack 2017), which uses the gamma-poisson distribution, have been developed to model single-cell data. SERGIO (Dibaeinia and Sinha 2019) and dyngen (Cannoodt et al. 2020) build on this by adding gene regulatory networks and multimodality respectively. These methods have been designed based on known deterministic relationships or synthetic reaction states, to which stochasticity is then added. However, it is computationally-intensive to model these reactions at scale or run many iterations for benchmarking. In some cases, it is only necessary to model the statistical variability and “noise” of RNA-Seq data in order to evaluate methods in the presence of multivariate correlation structures.

There is a need, therefore, for a systematic framework for statistical modelling and simulation of gene expression data derived from hypothetical, inferred or known gene networks. Here we present a package to achieve this, where samples from a multivariate normal distribution are used to generate normally-distributed log-expression data, with correlations between genes derived from the structure of the underlying pathway or gene regulatory network. This methodology enables simulation of expression profiles that approximate the log-transformed and normalised data from microarray studies, as well as, bulk or single-cell RNA-Seq experiments. This procedure has been released as the `graphsim` package to enable the generation of simulated gene expression datasets containing pathway relationships from a known underlying network. These simulated datasets can be used to evaluate various bioinformatics methodologies, including statistical and network inference procedures.

## Methodology and software

Here we present a procedure to simulate gene expression data with correlation structure derived from a known graph structure. This procedure assumes that transcriptomic data have been generated and follow a log-normal distribution (i.e., *log*(*X_ij_*) ~ *MVN*(*μ, Σ*), where *μ* and Σ are the mean vector and variance-covariance matrix respectively, for gene expression data derived from a biological pathway) after appropriate normalisation (Law et al. 2014; Li et al. 2015). Log-normality of gene expression matches the assumptions of the popular `limma` package (Matthew E R. et al. 2015), which is often used for the analysis of intensity-based data from gene expression microarray studies and count-based data from RNA-Seq experiments. This approach has also been applied for modelling UMI-based count data from single-cell RNA-Seq experiments in the `DESCEND` R package (Wang et al. 2018).

In order to simulate transcriptomic data, a pathway is first constructed as a graph structure, using the `igraph` R package (Csardi and Nepusz 2006), with the status of the edge relationships defined (i.e, whether they activate or inhibit downstream pathway members). This procedure uses a graph structure such as that presented in Figure 1a. The graph can be defined by an adjacency matrix, *A* (with elements *A_ij_*), where

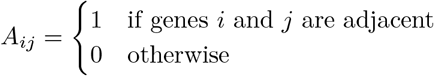

**Figure 1:**
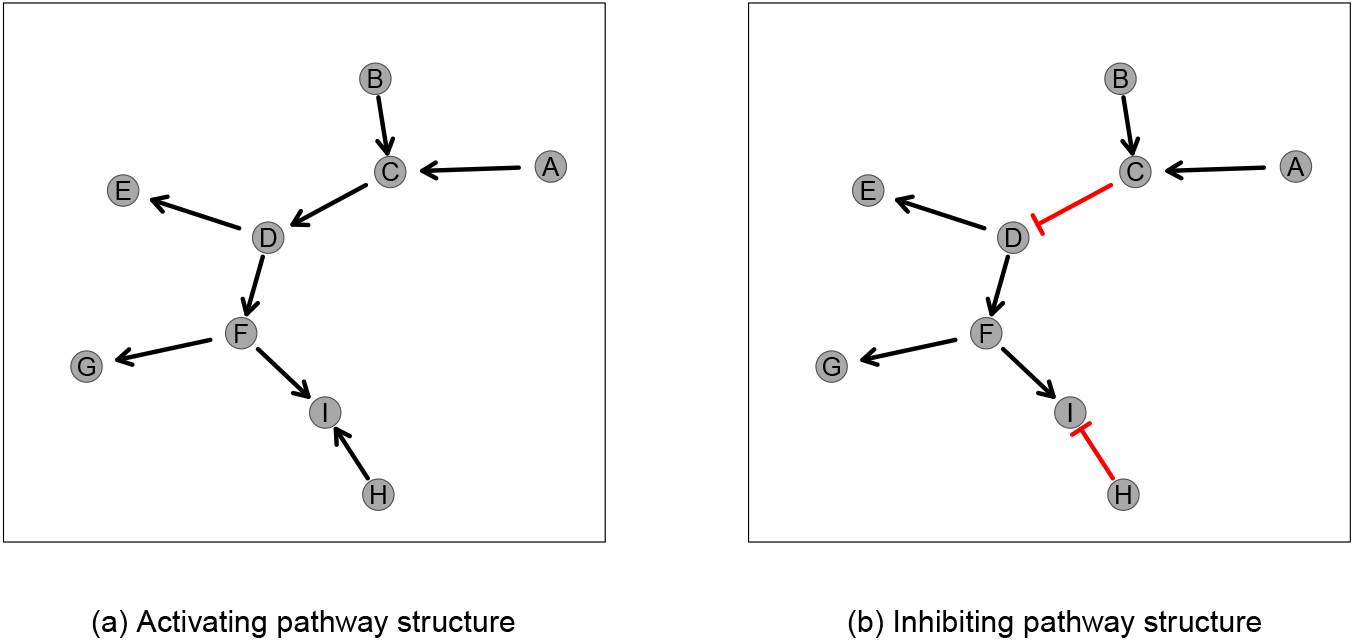
Simulated graph structures. A constructed graph structure used as an example to demonstrate the simulation procedure in Figures 2 and 3. Activating links are denoted by black arrows and inhibiting links by red edges.

A matrix, *R*, with elements *R_ij_*, is calculated based on distance (i.e., number of edges contained in the shortest path) between nodes, such that closer nodes are given more weight than more distant nodes, to define inter-node relationships. A geometrically-decreasing (relative) distance weighting is used to achieve this:

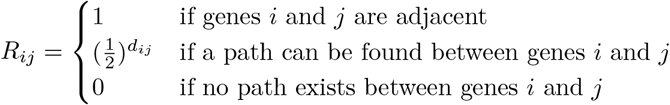

where *d_ij_* is the length of the shortest path (i.e., minimum number of edges traversed) between genes (nodes) *i* and *j* in graph *G.* Each more distant node is thus related by 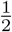 compared to the next nearest, as shown in Figure 2b. An arithmetically-decreasing (absolute) distance weighting is also supported in the `graphsim` R package which implements this procedure:

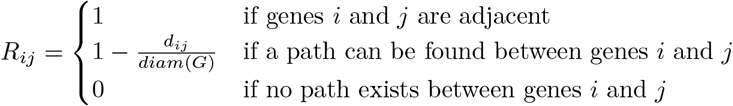

**Figure 2:**
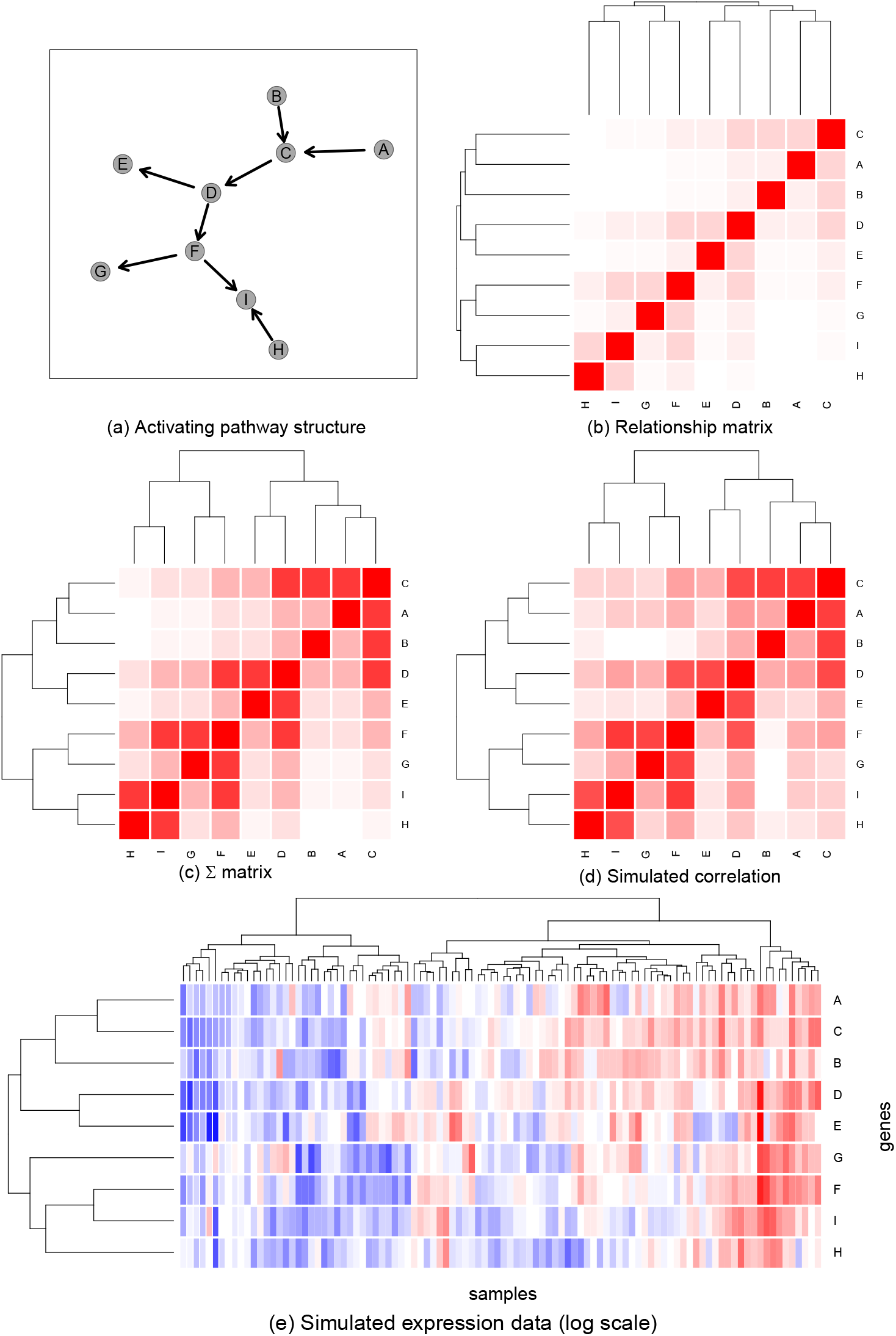
Simulating expression from a graph structure. An example of a graph structure (a) that has been used to derive a relationship matrix (b), Σ matrix (c) and correlation structure (d) from the relative distances between the nodes. Non-negative values are coloured white to red from 0 to 1 (e). The Σ matrix has been used to generate a simulated expression dataset of 100 samples (coloured blue to red from low to high) via sampling from the multivariate normal distribution. Here genes with closer relationships in the pathway structure show a higher correlation between simulated values.

Assuming a unit variance for each gene, these values can be used to derive a Σ matrix:

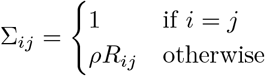

where *ρ* is the correlation between adjacent nodes. Thus covariances between adjacent nodes are assigned by a correlation parameter (*ρ*) and the remaining off-diagonal values in the matrix are based on scaling these correlations by the geometrically weighted relationship matrix (or the nearest positive definite matrix for Σ with negative correlations).

Computing the nearest positive definite matrix is necessary to ensure that the variance-covariance matrix can be inverted when used as a parameter in multivariate normal simulations, particularly when negative correlations are included for inhibitions (as shown below). Matrices that cannot be inverted occur rarely with biologically plausible graph structures but this approach allows for the computation of a plausible correlation matrix when the given graph structure is incomplete or contains loops. When required, the nearest positive definite matrix is computed using the `nearPD` function of the `Matrix` R package (Bates and Maechler 2016) to perform Higham’s algorithm (Higham 2002) on variance-covariance matrices. The `graphsim` package gives a warning when this occurs.

## Illustrations

### Generating a Graph Structure

The graph structure in Figure 1a was used to simulate correlated gene expression data by sampling from a multivariate normal distribution using the `mvtnorm` R package (Genz and Bretz 2009; Genz et al. 2016). The graph structure visualisation in Figure 1 was specifically developed for (directed) `igraph` objects in and is available in the `graphsim` package. The `plot_directed` function enables customisation of plot parameters for each node or edge, and mixed (directed) edge types for indicating activation or inhibition. These inhibition links (which occur frequently in biological pathways) are demonstrated in Figure 1b.

A graph structure can be generated and plotted using the following commands in R:

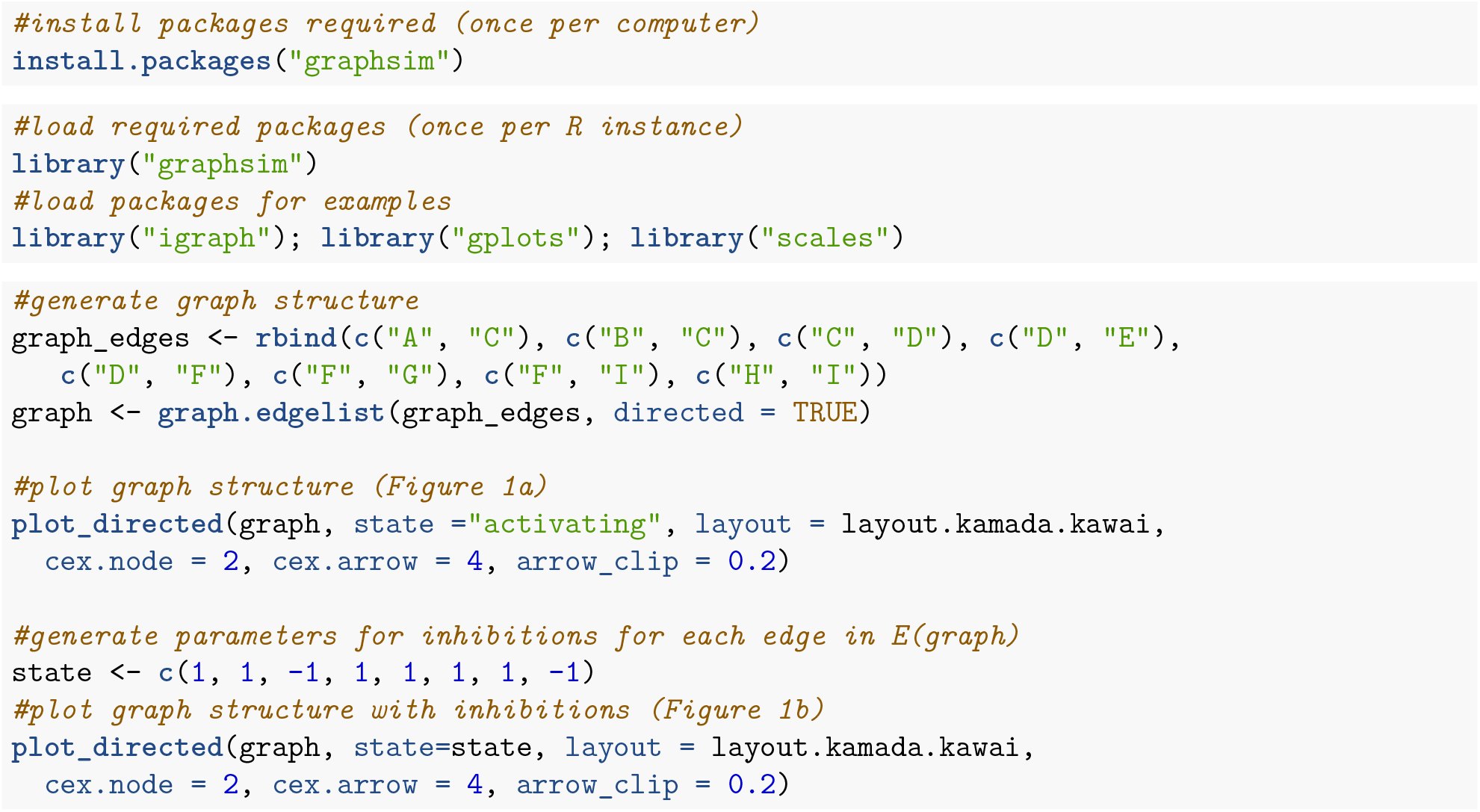

### Generating a Simulated Expression Dataset

The correlation parameter of *ρ* = 0.8 is used to demonstrate the inter-correlated datasets using a geometrically-generated relationship matrix (as used for the example in Figure 2c). This Σ matrix was then used to sample from a multivariate normal distribution such that each gene had a mean of 0, standard deviation 1, and covariance within the range [0,1] so that the off-diagonal elements of Σ represent correlations. This procedure generated a simulated (continuous normally-distributed) log-expression profile for each node (Figure 2e) with a corresponding correlation structure (Figure 2d). The simulated correlation structure closely resembled the expected correlation structure (Σ in Figure 2c) even for the relatively modest sample size (*N* = 100) illustrated in Figure 2. Once a gene expression dataset comprising multiple pathways has been generated (as in Figure 2e), it can then be used to test procedures designed for analysis of empirical gene expression data (such as those generated by microarrays or RNA-Seq) that have been normalised on a log-scale.

The simulated dataset can be generated using the following code:

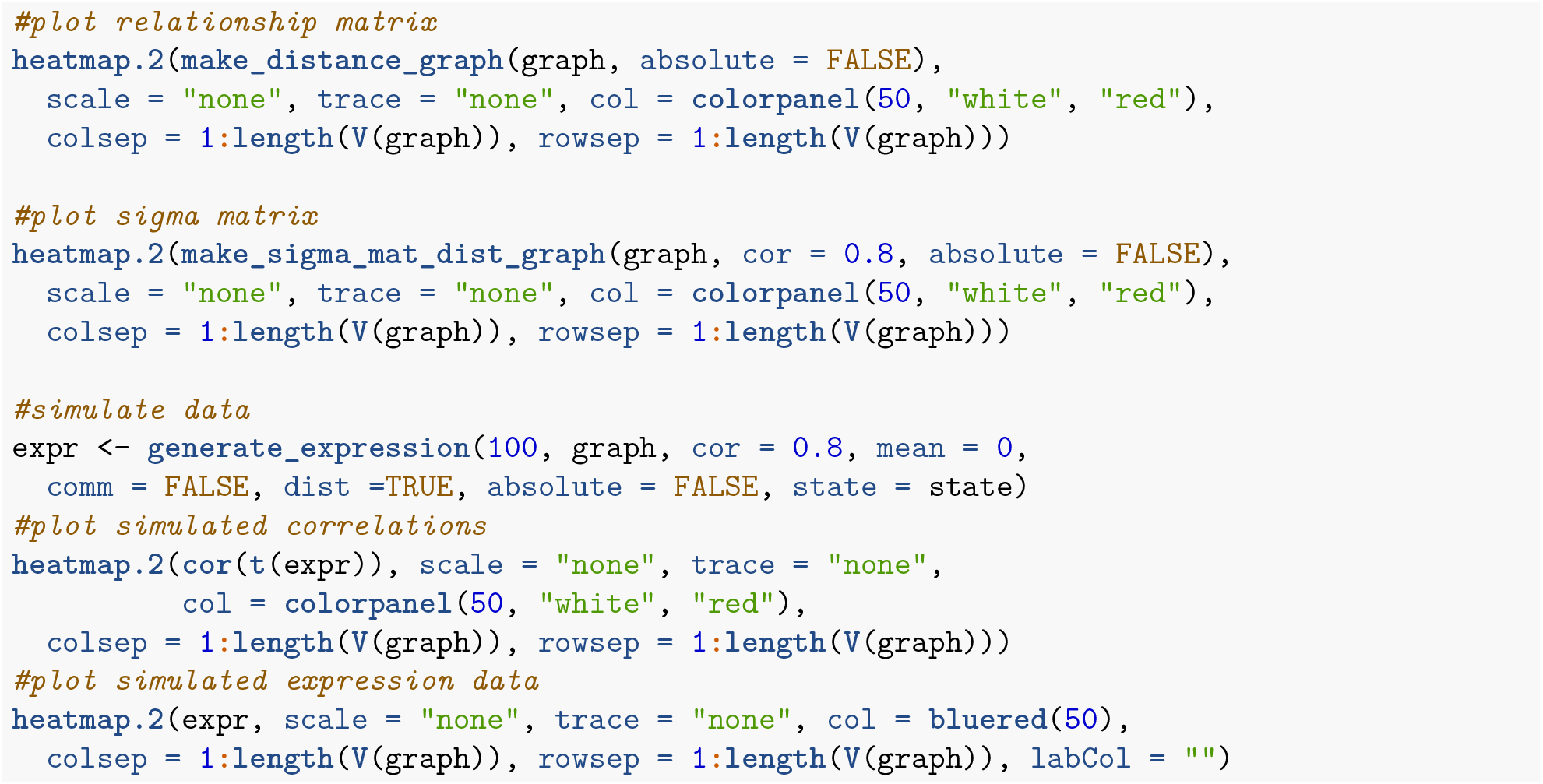

The simulation procedure (Figure 2) can similarly be used for pathways containing inhibitory links (Figure 3) with several refinements. With the inhibitory links (Figure 3a), distances are calculated in the same manner as before (Figure 3b) with inhibitions accounted for by iteratively multiplying downstream nodes by –1 to form modules with negative correlations between them (Figures 3c and 3d). A multivariate normal distribution with these negative correlations can be sampled to generate simulated data (Figure 3e).

**Figure 3:**
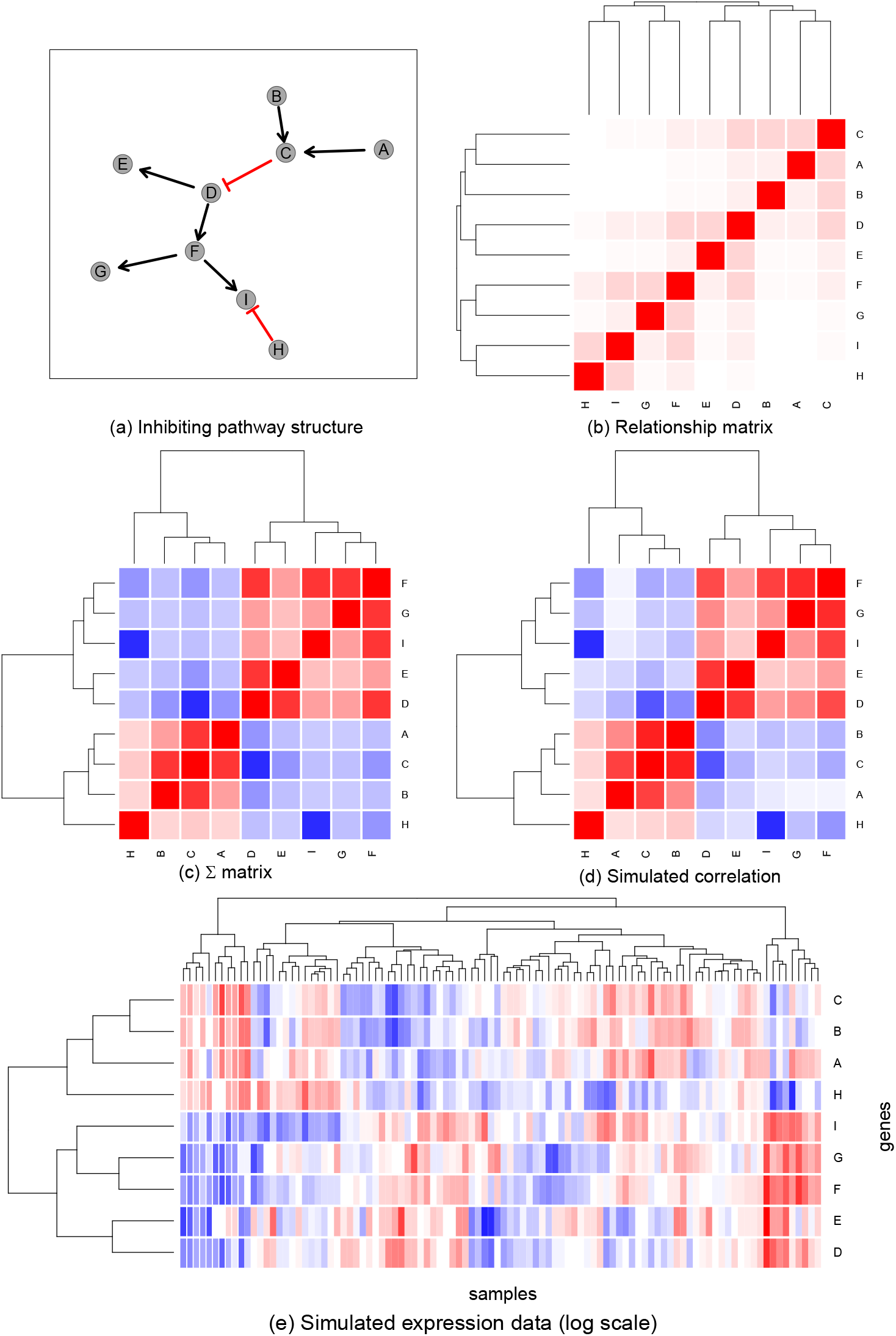
Simulating expression from graph structure with inhibitions. An example of a graph structure (a), that has been used to derive a relationship matrix (b), Σ matrix (c), and correlation structure (d), from the relative distances between the nodes (e). These values are coloured blue to red from –1 to 1. This has been used to generate a simulated expression dataset of 100 samples (coloured blue to red from low to high) via sampling from the multivariate normal distribution. Here the inhibitory relationships between genes are reflected in negatively correlated simulated values.

The following changes are needed to handle inhibitions:

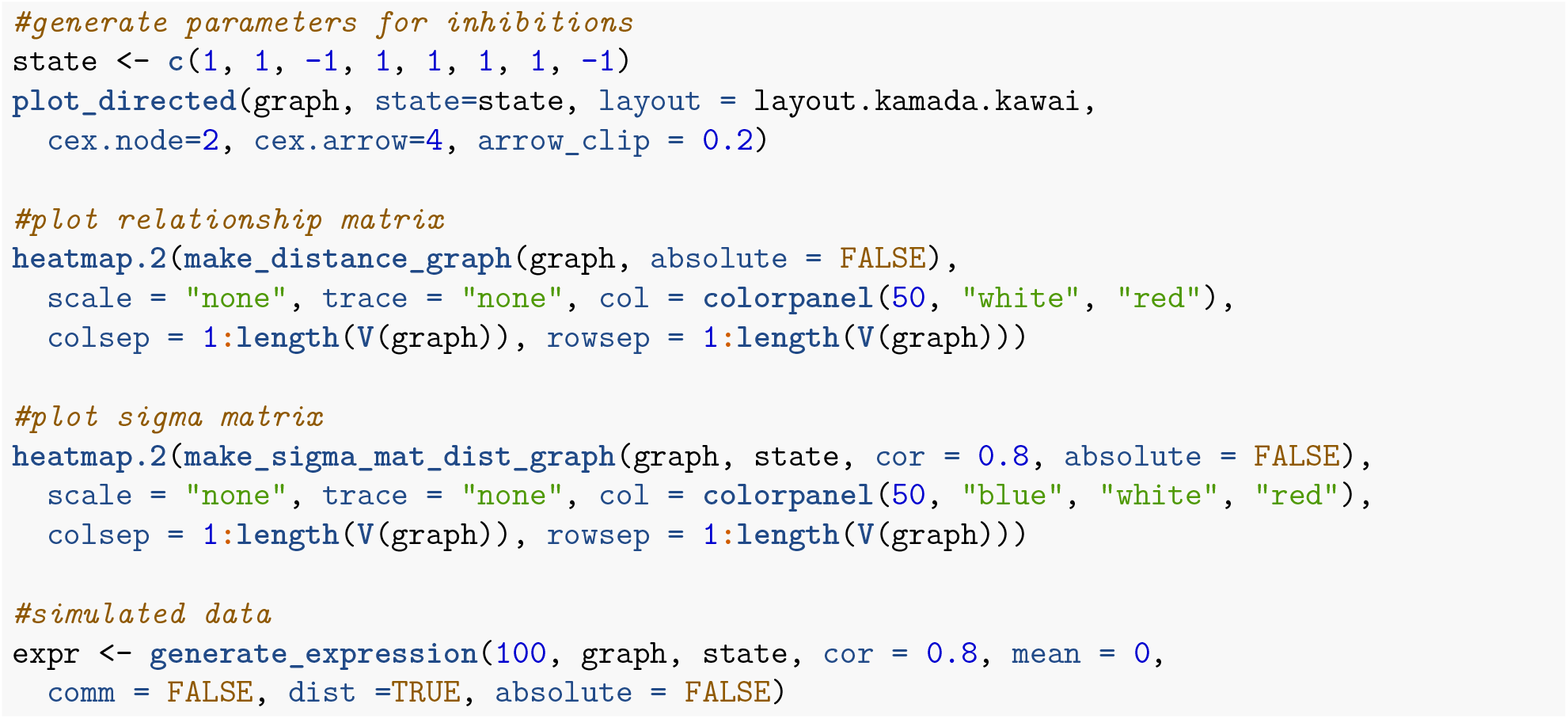

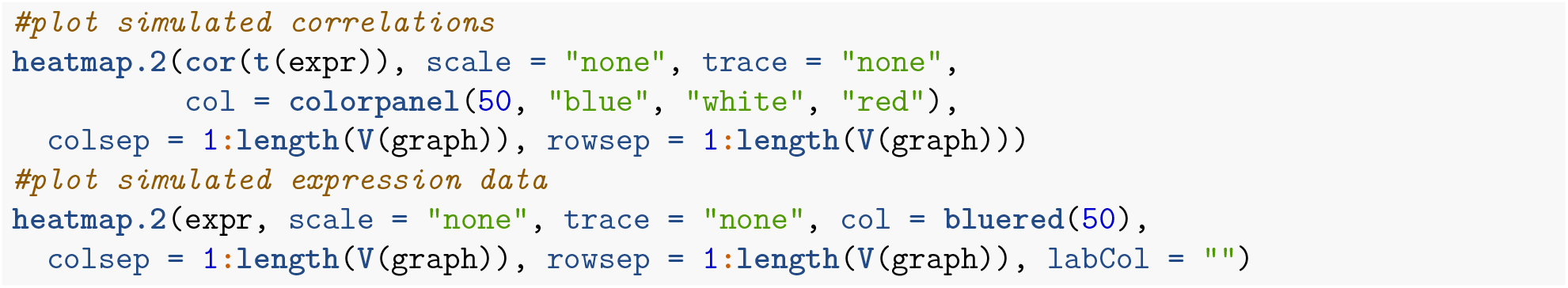

The simulation procedure is also demonstrated here (Figure 4) on a pathway structure for a known biological pathway (Reactome pathway R-HSA-2173789): “TGF-*β* receptor signaling activates SMADs” (Figure 4a) derived from the Reactome database version 52 (Croft et al. 2014). Distances are calculated in the same manner as before (Figure 4b) producing blocks of correlated genes (Figures 4c and 4d). This shows that the multivariate normal distribution can be sampled to generate simulated data to represent expression with the complexity of a biological pathway (Figure 4e). Here *SMAD7* exhibits negative correlations with the other SMADs consistent with its functions as an “inhibitor SMAD” which competitively inhibits *SMAD4.*

**Figure 4:**
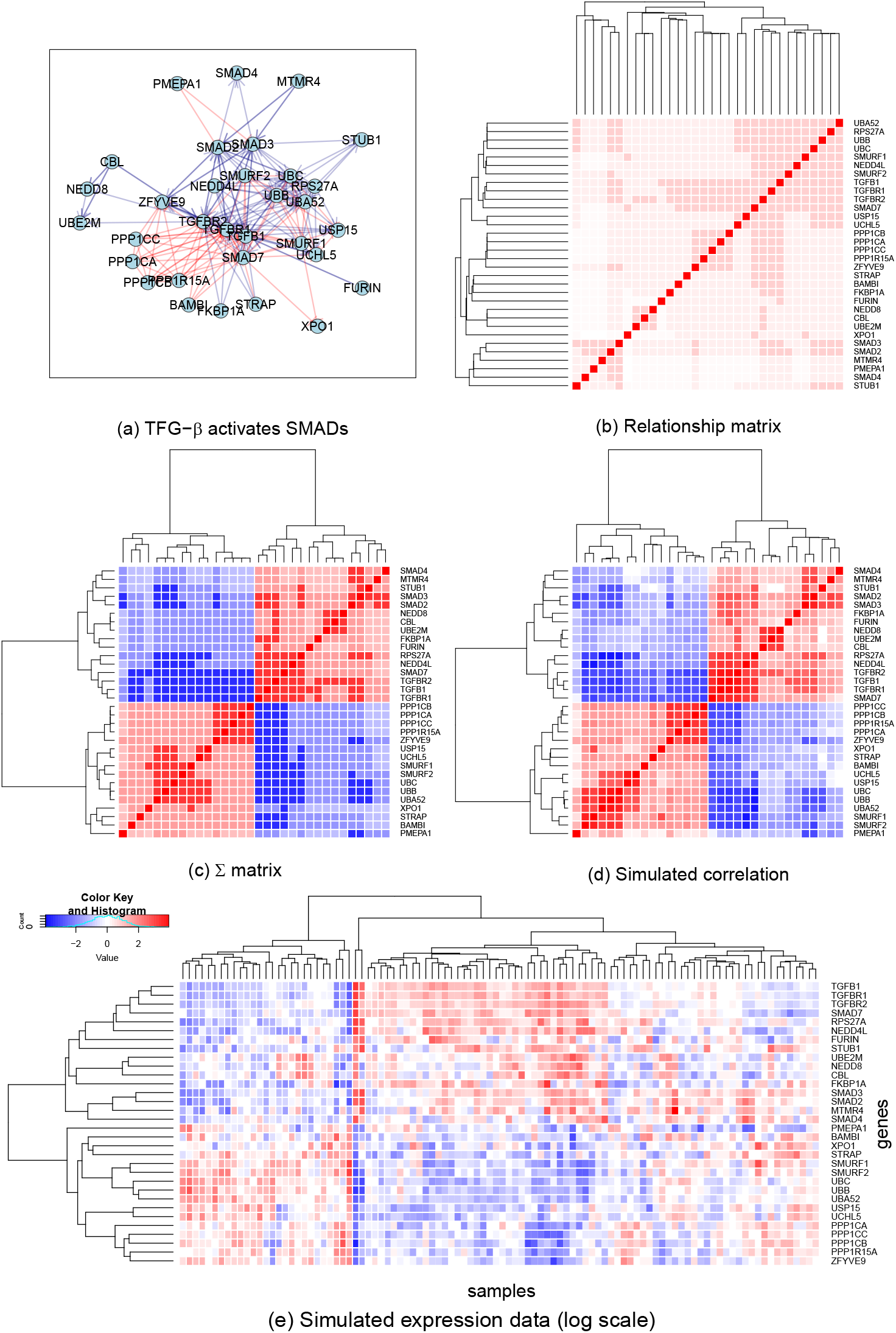
Simulating expression from a biological pathway graph structure. The graph structure (a) of a known biological pathway, “TGF-*β* receptor signaling activates SMADs” (R-HSA-2173789), was used to derive a relationship matrix (b), Σ matrix (c) and correlation structure (d) from the relative distances between the nodes. These values are coloured blue to red from –1 to 1 (e). This has been used to generate a simulated expression dataset of 100 samples (coloured blue to red from low to high) via sampling from the multivariate normal distribution. Here modules of genes with correlated expression can be clearly discerned.

We can import the graph structure into R as follows and run simulations as above:

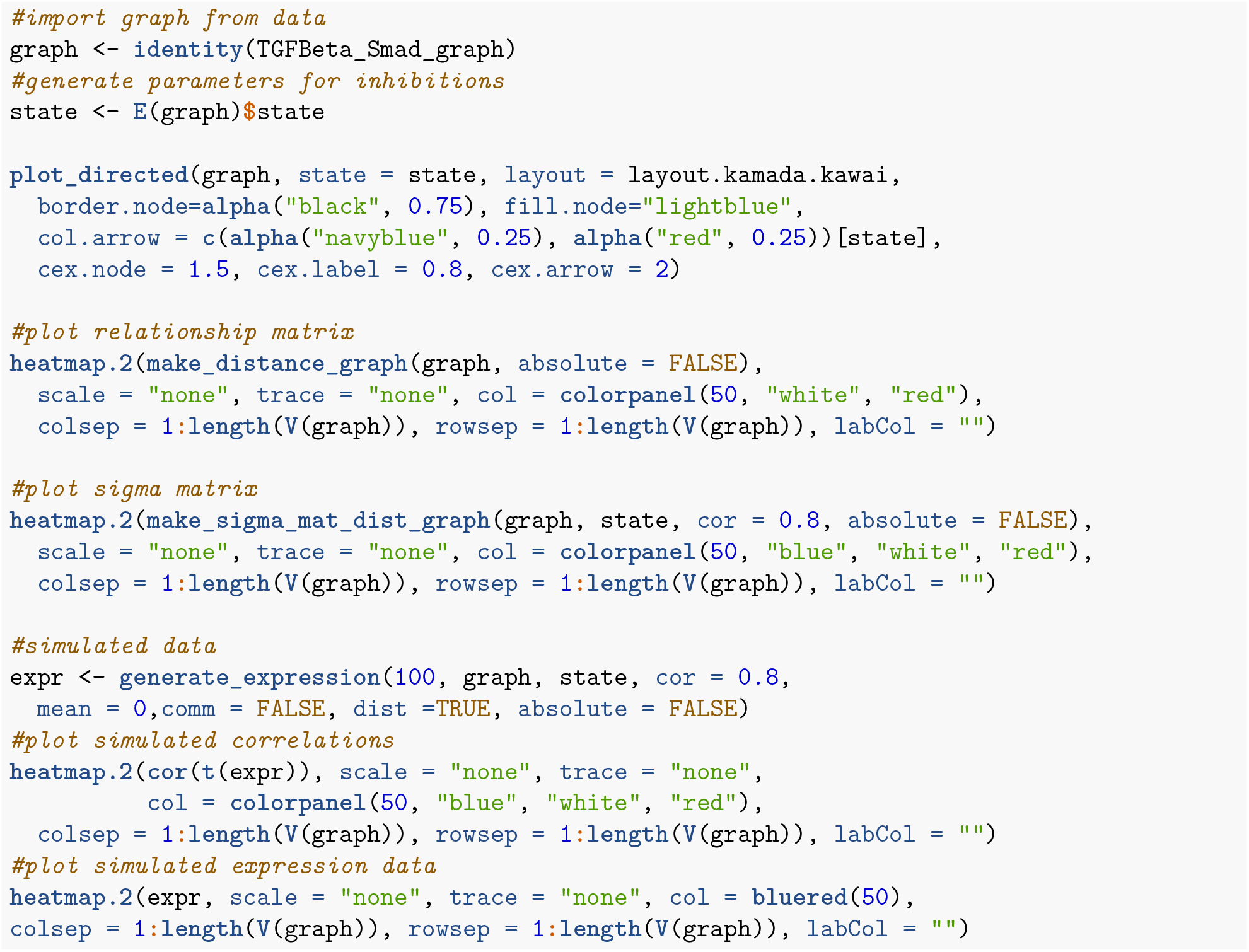

These simulated datasets can also be used for simulating gene expression data within a graph network to test genomic analysis techniques. Correlation structure can be included in datasets generated for testing whether true positive genes or samples can be detected in a sample with the background of complex pathway structure.

## Summary and discussion

Biological pathways are of fundamental importance to understanding molecular biology. In order to translate findings from genomics studies into real-world applications such as improved healthcare, the roles of genes must be studied in the context of molecular pathways. Here we present a statistical framework to simulate gene expression from biological pathways, and provide the `graphsim` package in R to generate these simulated datasets. This approach is versatile and can be fine-tuned for modelling existing biological pathways or for testing whether constructed pathways can be detected by other means. In particular, methods to infer biological pathways and gene regulatory networks from gene expression data can be tested on simulated datasets using this framework. The package also enables simulation of complex gene expression datasets to test how these pathways impact on statistical analysis of gene expression data using existing methods or novel statistical methods being developed for gene expression data analysis. This approach is intended to be applied to bulk gene expression data but could in principle be adapted to modelling single-cell or different modalities such as genome-wide epigenetic data.

## Computational details

The results in this paper were obtained using R 4.0.2 with the `igraph` 1.2.5 `Matrix` 1.2-17, `matrixcalc` 1.0-3, and mvtnorm 1.1-1 packages. R itself and all dependent packages used are available from the Comprehensive Archive Network (CRAN) at https://CRAN.R-project.org. The `graphsim` 1.0.0 package presented can be installed from CRAN and the issues can be reported to the development version on GitHub (https://github.com/TomKellyGenetics/graphsim). This package is included in the `igraph.extensions` library on GitHub (https://github.com/TomKellyGenetics/igraph.extensions) which installs various tools for `igraph` analysis. This software is cross-platform and compatible with installations on Windows, Mac, and Linux operating systems. The package GitHub repository also contains vignettes with more information and examples on running functions released in the package. Updates to the package (`graphsim` 1.0.0) will be released on CRAN.

Complete examples of code needed to produce the figures in this paper are available in the Rmarkdown version in the package GitHub repository (https://github.com/TomKellyGenetics/graphsim/paper). Further details are available in the vignettes as well.

## Supporting information

graphsim 1.0.0 R package

graphsim 1.0.0 R package

## Acknowledgements

This package was developed as part of a PhD research project funded by the Postgraduate Tassell Scholarship in Cancer Research Scholarship awarded to STK. We thank members of the Laboratory of Professor Satoru Miyano at the University of Tokyo, Institute for Medical Science, Professor Seiya Imoto, Associate Professor Rui Yamaguchi, and Dr Paul Sheridan (Assistant Professor at Hirosaki University, CSO at Tupac Bio) for helpful discussions in this field. We also thank Professor Parry Guilford at the University of Otago, Professor Cristin Print at the University of Auckland, and Dr Erik Arner at the RIKEN Center for Integrative Medical Sciences for their excellent advice during this project.

## Author Contributions

S.T.K. and M.A.B. conceived of the presented methodology. S.T.K. developed the theory and performed the computations. M.A.B. provided guidance throughout the project and gave feedback on the package. All authors discussed the package and contributed to the final manuscript.

