## Supplementary material for "graphsim: An R package for simulating gene expression data from graph structures of biological pathways": graphsim 1.0.0 R package: DESCRIPTION

Package: graphsim
Type: Package
Title: Simulate Expression Data from 'igraph' Networks
Version: 1.0.0
Date: 2020-06-15
Description: Functions to develop simulated continuous data (e.g., gene expression) from a sigma covariance matrix derived from a graph structure in 'igraph' objects. Intended to extend 'mvtnorm' to take 'igraph'  structures rather than sigma matrices as input. This allows the use of simulated data that correctly accounts for pathway relationships and correlations. This allows the use of simulated data that correctly accounts for pathway relationships and correlations. Here we present a versatile statistical framework to simulate  correlated gene expression data from biological pathways, by sampling from a multivariate normal distribution derived from a graph structure. This package allows the simulation of biological pathways from a graph structure based on a statistical model of gene expression. For example methods to infer biological pathways and gene regulatory networks from gene expression data can be tested on simulated datasets using this framework. This also allows for pathway structures to be considered as a confounding variable when simulating gene expression data to test the performance of genomics analyses.
License: GPL-3
BugReports: https://github.com/TomKellyGenetics/graphsim/issues/
Depends: R (>= 2.10)
Imports: utils, gplots, igraph, mvtnorm, Matrix, matrixcalc, graphics
Suggests: devtools, knitr (>= 1.5), prettydoc, rmarkdown, testthat,
        scales, vdiffr
LazyData: TRUE
RoxygenNote: 7.1.1
Encoding: UTF-8
VignetteBuilder: knitr
NeedsCompilation: no
Packaged: 2020-07-01 03:18:31 UTC; tom
Author: S. Thomas Kelly [ctb],
  S. Thomas Kelly [aut, cre],
  Michael A. Black [aut, ths]
Maintainer: S. Thomas Kelly <>
