## Supplementary material for "graphsim: An R package for simulating gene expression data from graph structures of biological pathways": graphsim 1.0.0 R package: plots_directed.html

graphsim: directed plots for igraph objects


### graphsim: directed plots for igraph objects

###### S. Thomas Kelly^1,2^, Michael A. Black^2^ ^1^ Department of Biochemistry, University of Otago, PO Box 56, Dunedin 9054, New Zealand ^2^ RIKEN Center for Integrative Medical Sciences, Suehiro-cho-1-7-22, Tsurumi Ward, Yokohama

###### Tuesday 30 June 2020

- 1 Directed graph plots
  - 1.1 Motivations
  - 1.2 Getting started
- 2 Toy example
  - 2.1 Set up simulated graph
  - 2.2 Plotting
    - 2.2.1 Custom aesthetics
      - 2.2.1.1 Vectorisation of aesthetics
    - 2.2.2 Arrow customisation
      - 2.2.2.1 Vectorisation of edge properties
- 3 Empirical examples
  - 3.1 RAF/MAP kinase cascade
  - 3.2 Pi3K cascade
  - 3.3 TGFβ-Smad pathway
- 4 Session info
- 5 References

**Summary**

Guide on how to plot directed graphs from `igraph` objects to display activating and repressive states.

**Package**

graphsim 1.0.0

### 1 Directed graph plots

#### 1.1 Motivations

Here we demonstrate the plotting functions that come built-in with `graphsim`, an alternative to the `plot.igraph` provided by the `igraph` package. Here we provide additional functionality for plotting directed graphs. This draws upon many functions provided by `igraph`, including layout settings (Csardi and Nepusz 2006).

In particular, graph and network representations in biology often require displaying edge properties (Barabási and Oltvai 2004). Here we have the “state” parameter which can be used to differentiate these, and allow us to represent activating and inhibiting or repressing relationships differently. We use different arrowheads as per convention in molecular biology.

#### 1.2 Getting started

To generate these plots, the following packages must be imported.

```
library("igraph")
library("scales")
library("graphsim")
```

### 2 Toy example

Here we demonstrate the plot functions on a small toy graph.

#### 2.1 Set up simulated graph

First we generate a graph object using the `igraph` package.

```
graph_edges <- rbind(c("A", "C"), c("B", "C"), c("C", "D"), c("D", "E"),
                     c("D", "F"), c("F", "G"), c("F", "I"), c("H", "I"))
graph <- graph.edgelist(graph_edges, directed = TRUE)
```

#### 2.2 Plotting

We next demonstrate the plotting function for directed graph objects. `plot_directed` with default settings uses the `layout.fruchterman.reingold` as does built-in plotting function `igraph::plot.igraph`. This function provides additional functionality to displaying directed graphs in particular.

```
plot_directed(graph)
```

Here you can see that the plotting function displays a graph in a similar layout to `plot.igraph` with different aesthetic parameters. We suggest that you choose the function that suits your needs and datasets. We demonstrate the features available for `plot_directed` below.

##### 2.2.1 Custom aesthetics

We support various aesthetic parameters to control the colour and relative size of nodes and edges.

`plot_directed` supports customised layouts:

```
plot_directed(graph, layout = layout.kamada.kawai)
```

In addition, custom colouts are supported:

```
plot_directed(graph, fill.node = "lightblue", border.node = "royalblue")
```

###### 2.2.1.1 Vectorisation of aesthetics

Colours may also be entered as a vector for each node in `V(graph)`:

```
names(V(graph))
#> [1] "A" "C" "B" "D" "E" "F" "G" "I" "H"
colour_vector <- ifelse(names(V(graph)) %in% c("A", "D", "I"), 1, 2)
plot_directed(graph, fill.node = c("lightblue", "grey")[colour_vector], border.node = c("royalblue", "black")[colour_vector])
```

This functionality allows highlighting of particular groups based on known properties of the graph. For examples `V(graph)$type` for bipartite graphs or partitions from Louvain (`igraph::cluster_louvain`) or Leiden (`leiden::leiden`) clustering algorithms.

##### 2.2.2 Arrow customisation

The `state` parameter controls whether the links are “activating” or “inhibiting”. These can denote activation and repression: foe example, positive and negative regulation of genes or kinase and phosphatase activity of proteins. These may be specified globally as either a character string or numeric:

Activating links are displated with any of the following:

- “activating”
- `1`
- `0`

```
plot_directed(graph, state = "activating")
```

Note that activating states can also be specified as follows:

```
plot_directed(graph, state = 1)
plot_directed(graph, state = 0)
```

Inhibiting links are displated with any of the following:

- “inhibiting”
- `-1`
- `2`

```
plot_directed(graph, state = "inhibiting")
```

Note that inhibiting states can also be specified as follows:

```
plot_directed(graph, state = -1)
plot_directed(graph, state = 2)
```

###### 2.2.2.1 Vectorisation of edge properties

The state parameter may also be applied as a vector to each edge in `E(graph)` respectively.

```
E(graph)
#> + 8/8 edges from 30607b1 (vertex names):
#> [1] A->C B->C C->D D->E D->F F->G F->I H->I
plot_directed(graph, state = c(1, 1, -1, -1, 1, -1, 1, -1))
```

Note that by default, inhibiting relationships are highlighted with a different `col.arrow` value, which can be controlled by the input parameter.

```
edge_properties <- c(1, 1, -1, -1, 1, -1, 1, -1)/2 + 1.5
plot_directed(graph, state = edge_properties, col.arrow = c("darkgreen", "red")[edge_properties])
```

```
edge_properties <- c(1, 1, -1, -1, 1, -1, 1, -1)/2 + 1.5
ggplot_colours <- c("#F8766D", "#CD9600", "#7CAE00", "#00BE67",
                    "#00BFC4", "#00A9FF", "#C77CFF", "#FF61CC")
plot_directed(graph, state = edge_properties,
              col.arrow = ggplot_colours, fill.node =  ggplot_colours)
```

### 3 Empirical examples

Here we demonstrate using the plotting package to display real biological pathways from the “Reactome” database (Croft *et al*. 2014). We can import these from the `data` directory included with this package. These graphs are given for examples and convenience. Any empirical data that consists of a list of directed edges can be imported as an igraph object and handled similarly. Below are some demonstrations.

#### 3.1 RAF/MAP kinase cascade

Here we plot the RAF/MAP kinase cascade pathway.

```
graph <- identity(RAF_MAP_graph)
plot_directed(graph,col.arrow = alpha("#00A9FF", 0.25),
              fill.node = "lightblue",
              layout = layout.kamada.kawai)
```

#### 3.2 Pi3K cascade

Here we plot the phosphoinositide-3-kinase (Pi3K) cascade pathway.

```
graph <- identity(Pi3K_graph)
plot_directed(graph, col.arrow = alpha("#00A9FF", 0.25),
              fill.node = "lightblue",
              layout = layout.kamada.kawai)
```

#### 3.3 TGFβ-Smad pathway

Here we plot the TGFβ-Smad pathway with inhibitions known. States are imported as edge attributes from the imported graph.

```
graph <- identity(TGFBeta_Smad_graph)
edge_properties <- E(graph)$state
plot_directed(graph, state = edge_properties,
              col.arrow = c(alpha("navyblue", 0.25), alpha("red", 0.25))[edge_properties],
              fill.node = c("lightblue"),
              layout = layout.kamada.kawai)
```

---

### 4 Session info

Here is the output of `sessionInfo()` on the system on which this document was compiled running pandoc 2.1:

```
#> R version 4.0.2 (2020-06-22)
#> Platform: x86_64-apple-darwin17.0 (64-bit)
#> Running under: macOS Mojave 10.14.6
#> 
#> Matrix products: default
#> BLAS:   /Library/Frameworks/R.framework/Versions/4.0/Resources/lib/libRblas.dylib
#> LAPACK: /Library/Frameworks/R.framework/Versions/4.0/Resources/lib/libRlapack.dylib
#> 
#> locale:
#> [1] en_GB.UTF-8/en_GB.UTF-8/en_GB.UTF-8/C/en_GB.UTF-8/en_GB.UTF-8
#> 
#> attached base packages:
#> [1] stats     graphics  grDevices utils     datasets  methods  
#> [7] base     
#> 
#> other attached packages:
#> [1] graphsim_1.0.0 scales_1.1.1   igraph_1.2.5  
#> 
#> loaded via a namespace (and not attached):
#>  [1] knitr_1.29         magrittr_1.5       munsell_0.5.0     
#>  [4] colorspace_1.4-1   lattice_0.20-41    R6_2.4.1          
#>  [7] rlang_0.4.6        stringr_1.4.0      matrixcalc_1.0-3  
#> [10] caTools_1.18.0     tools_4.0.2        grid_4.0.2        
#> [13] xfun_0.15          KernSmooth_2.23-17 htmltools_0.5.0   
#> [16] gtools_3.8.2       yaml_2.2.1         digest_0.6.25     
#> [19] lifecycle_0.2.0    Matrix_1.2-18      farver_2.0.3      
#> [22] bitops_1.0-6       prettydoc_0.3.1    evaluate_0.14     
#> [25] rmarkdown_2.3      gdata_2.18.0       stringi_1.4.6     
#> [28] compiler_4.0.2     gplots_3.0.3       mvtnorm_1.1-1     
#> [31] pkgconfig_2.0.3
```

---
