## Supplementary material for "graphsim: An R package for simulating gene expression data from graph structures of biological pathways": graphsim 1.0.0 R package: simulate_expression.html

graphsim: simulating continuous data based on network graph structures


### graphsim: simulating continuous data based on network graph structures

###### S. Thomas Kelly^1,2^, Michael A. Black^2^ ^1^ Department of Biochemistry, University of Otago, PO Box 56, Dunedin 9054, New Zealand ^2^ RIKEN Center for Integrative Medical Sciences, Suehiro-cho-1-7-22, Tsurumi Ward, Yokohama

###### Tuesday 30 June 2020

- 1 Overview of graphsim
  - 1.1 Motivations
- 2 Getting Started
  - 2.1 Install dependencies
  - 2.2 Set up simulated graphs
- 3 Generating simulated expression data from graph
  - 3.1 Minimal example
  - 3.2 How it works step-by-step
    - 3.2.1 Adjacency matrix
    - 3.2.2 Distance matrix
    - 3.2.3 Sigma (Σ) matrix
    - 3.2.4 Simulated expression and observed correlation
  - 3.3 Summary
    - 3.3.1 Inhibiting relationships
- 4 Examples
  - 4.1 Toy examples
    - 4.1.1 Diverging branches
      - 4.1.1.1 Inhibiting relationships
    - 4.1.2 Converging branches
      - 4.1.2.1 Inhibiting relationships
    - 4.1.3 Reconnecting paths
      - 4.1.3.1 Inhibiting relationships
  - 4.2 Empirical examples
    - 4.2.1 Kinase pathways
      - 4.2.1.1 RAF/MAP kinase cascade
      - 4.2.1.2 Phosphoinositide-3-kinase cascade
    - 4.2.2 Pathways with repression
      - 4.2.2.1 The Pi3K/AKT pathway
      - 4.2.2.2 The TGFβ-Smad pathway
- 5 Summary
  - 5.1 Citation
  - 5.2 Reporting issues
- 6 Session info
- 7 References

**Summary**

Guide on how to simulate gene expression data from pathway structures defined by graphs from `igraph` objects.

**Package**

graphsim 1.0.0

### 1 Overview of graphsim

This package is designed to balance user-friendliness (by providing sensible defaults and built-in functions) and flexbility (many options are available to use as needed).

If you have problems or feedback, sumbmitting an issue to the the GitHub repository is encouraged. See the DESCRIPTION and README.md for more details on how to suggest changes to the package.

#### 1.1 Motivations

Pathway and graph structures have a wide array of applications. Here we consider the simulation of (log-normalised) gene expression data from a biological pathway. If you have another use for this software you are welcome to apply it to your problem but please bear in mind that it was designed with this application in mind. In principle, normally-distributed continuous data can be generated based on any defined relationships. This package uses the graph structure to define a ∑ covariance matrix and generate simulated data by sampling from a multivariate normal distribution.

Crucially, this allows the simulation of negative correlations based on inhibitory or repressive relationships, as commonly occur in biology (Barabási and Oltvai 2004). A custom plotting function `plot_directed` is provided to visualise these relationships using the “state” parameter. This plotting function has a dedicated vignette.

For more details on the background of this package, see the paper included with the package on GitHub. This vignette provides more detail on the code needed to reproduce the figures presented in the manuscript.

### 2 Getting Started

#### 2.1 Install dependencies

The package can be installed as follows. Run the following command to install the current release from CRAN (recommended).

```
#install packages required (once per computer)
install.packages("graphsim")
```

Run the following command to install the development version from GitHub (advanced users). This will import the latest changes to the package so behaviour may be unstable.

```
#install stable release
remotes::install_github("TomKellyGenetics", ref = "master")
#install development version
remotes::install_github("TomKellyGenetics", ref = "dev")
```

Once the required packages are installed, we must load the packages required to use the package functions with `igraph` objects and to generate plots (Csardi and Nepusz 2006). Here `igraph` is required to create `igraph` objects and `gplots` is required for plotting heatmaps.

```
library("igraph")
library("gplots")
library("graphsim")
library("scales")
```

#### 2.2 Set up simulated graphs

Here we set up a simple graph to demonstrate how connections in the graph structure lead to correlations in the final output. We create a simple pathway of 9 genes with various branches.

```
graph_structure_edges <- rbind(c("A", "C"), c("B", "C"), c("C", "D"),c("D", "E"),
                               c("D", "F"), c("F", "G"), c("F", "I"), c("H", "I"))
graph_structure <- graph.edgelist(graph_structure_edges, directed = T)
plot_directed(graph_structure, layout = layout.kamada.kawai)
```

### 3 Generating simulated expression data from graph

#### 3.1 Minimal example

A simulated dataset can be generated with a single command. This is all that is required to get started.

```
expr <- generate_expression(100, graph_structure, cor = 0.8, mean = 0,
                            comm = FALSE, dist = TRUE, absolute = FALSE)
heatmap.2(expr, scale = "none", trace = "none",
          col = bluered(50), colsep = 1:4, rowsep = 1:4)
```

Here we’ve generated a simulated dataset of 100 samples with gene expression data for the genes in the graph shown above. We will show below how these are used to demonstrate what computations are being performed to generate this data from the graph structure given.

Various arguments are supported to alter how the simulated datasets are computed. The other functions used below are called internally by `generate_expression` and are not needed to compute the final dataset in the above heatmap plot. See the documentation for details.

#### 3.2 How it works step-by-step

Here we show the data generated by this graph structure. This demonstrates how several of the available options are used to compute the necessary steps.

##### 3.2.1 Adjacency matrix

The data can be summarised by an “adjacency matrix” where a one (1) is given between a row `i` and column `j` if there is an edge between genes `i` and `j`. Otherise it is a zero (0) for genes that are not connected. For an undirected graph, edges are shown in a symmetical matrix.

```
adj_mat <- make_adjmatrix_graph(graph_structure)
heatmap.2(make_adjmatrix_graph(graph_structure),
          scale = "none", trace = "none",
          col = colorpanel(3, "grey75", "white", "blue"),
          colsep = 1:4, rowsep = 1:4)
```

For a directed graph, the adjacency matrix may be assymetric. A non-zero element `adjmat[i,j]` represents the presence or weight of the edge from gene `i` (matrix rows) to gene `j` (matrix columns).

```
heatmap.2(make_adjmatrix_graph(graph_structure, directed = T),
          scale = "none", trace = "none",
          col = colorpanel(3, "grey75", "white", "blue"),
          colsep = 1:4, rowsep = 1:4)
```

We can compute the common links between each pair of genes. This shows how many genes are connected to both genes `i` and `j`.

```
comm_mat <- make_commonlink_graph(graph_structure)
heatmap.2(make_commonlink_graph(graph_structure),
          scale = "none", trace = "none",
          col = colorpanel(50, "grey75", "red"),
          colsep = 1:4, rowsep = 1:4)
```

This shows how many edges to a shared neighbour these nodes have between them. The diagonal will therefore reflect vertex degree as all edges are counted.

We define commonlinks between each pair of nodes as how many nodes are mutually connected to both of the nodes in the adjacency matrix (how many paths of length 2 exist between them). Note that this weights towards genes with a higher vertex degree (as does the Laplacian).

The Laplacian matrix has the same dimensions as the adjancency matrix. For undirected graphs it is a symmetric matrix but for directed graphs it is not. It has the same number of rows and columns as the number of nodes.

The Laplacian matrix is defined as `laplacian[i,j] = vertex_degree(i)` if `i == j` and `laplacian[i,j] = -1` if `i != j`. The wieghted Laplacian matrix is defined as `laplacian[i,j] = -wieghts(graph)[i,j]` for the off-diagonal terms.

```
laplacian_mat <- make_laplacian_graph(graph_structure)
heatmap.2(make_laplacian_graph(graph_structure),
          scale = "none", trace = "none", 
          col = bluered(50),colsep = 1:4, rowsep = 1:4)
```

As expected, the off-diagonal terms of the Laplacian are negative integer values and the diagonals reflect the vertex degree.

##### 3.2.2 Distance matrix

To compute the relationships between each gene by “distance” we first compute the shortest paths between each pair of nodes, using Dijkstra’s algorithm (Prim 1957; Dijkstra 1959).

```
shortest.paths(graph_structure)
```

```
##   A C B D E F G I H
## A 0 1 2 2 3 3 4 4 5
## C 1 0 1 1 2 2 3 3 4
## B 2 1 0 2 3 3 4 4 5
## D 2 1 2 0 1 1 2 2 3
## E 3 2 3 1 0 2 3 3 4
## F 3 2 3 1 2 0 1 1 2
## G 4 3 4 2 3 1 0 2 3
## I 4 3 4 2 3 1 2 0 1
## H 5 4 5 3 4 2 3 1 0
```

```
heatmap.2(shortest.paths(graph_structure),
          scale = "none", trace = "none",
          col = colorpanel(50, "grey75", "red"),
           thincolsep = 1:4, rowsep = 1:4)
```

```
## Warning in plot.window(...): "thincolsep" is not a graphical
## parameter
```

```
## Warning in plot.xy(xy, type, ...): "thincolsep" is not a graphical
## parameter
```

```
## Warning in title(...): "thincolsep" is not a graphical parameter
```

Here we plot the number of edges in the shortest paths between each pair of nodes in the graph (as an integrer value). Relative to the “diameter” (length of the longest shortest path between any 2 nodes), we can show which genes are more similar or different based on the graph structure.

```
diam <- diameter(graph_structure)
relative_dist <- (1+diam-shortest.paths(graph_structure))/diam
relative_dist
```

```
##      A    C    B    D    E    F    G    I    H
## A 1.25 1.00 0.75 0.75 0.50 0.50 0.25 0.25 0.00
## C 1.00 1.25 1.00 1.00 0.75 0.75 0.50 0.50 0.25
## B 0.75 1.00 1.25 0.75 0.50 0.50 0.25 0.25 0.00
## D 0.75 1.00 0.75 1.25 1.00 1.00 0.75 0.75 0.50
## E 0.50 0.75 0.50 1.00 1.25 0.75 0.50 0.50 0.25
## F 0.50 0.75 0.50 1.00 0.75 1.25 1.00 1.00 0.75
## G 0.25 0.50 0.25 0.75 0.50 1.00 1.25 0.75 0.50
## I 0.25 0.50 0.25 0.75 0.50 1.00 0.75 1.25 1.00
## H 0.00 0.25 0.00 0.50 0.25 0.75 0.50 1.00 1.25
```

```
heatmap.2(relative_dist, scale = "none", trace = "none",
          col = colorpanel(50, "grey75", "red"),
          colsep = 1:4, rowsep = 1:4)
```

These relationships are used to create a distance graph relative to the diameter. A relative geometrically decreasing distance is computed as follows. In this case every connected node is weighted in fractions of the diameter.

```
dist_mat <- make_distance_graph(graph_structure, absolute = F)
dist_mat
```

```
##            A          C          B          D          E          F
## A 1.00000000 0.20000000 0.10000000 0.10000000 0.06666667 0.06666667
## C 0.20000000 1.00000000 0.20000000 0.20000000 0.10000000 0.10000000
## B 0.10000000 0.20000000 1.00000000 0.10000000 0.06666667 0.06666667
## D 0.10000000 0.20000000 0.10000000 1.00000000 0.20000000 0.20000000
## E 0.06666667 0.10000000 0.06666667 0.20000000 1.00000000 0.10000000
## F 0.06666667 0.10000000 0.06666667 0.20000000 0.10000000 1.00000000
## G 0.05000000 0.06666667 0.05000000 0.10000000 0.06666667 0.20000000
## I 0.05000000 0.06666667 0.05000000 0.10000000 0.06666667 0.20000000
## H 0.04000000 0.05000000 0.04000000 0.06666667 0.05000000 0.10000000
##            G          I          H
## A 0.05000000 0.05000000 0.04000000
## C 0.06666667 0.06666667 0.05000000
## B 0.05000000 0.05000000 0.04000000
## D 0.10000000 0.10000000 0.06666667
## E 0.06666667 0.06666667 0.05000000
## F 0.20000000 0.20000000 0.10000000
## G 1.00000000 0.10000000 0.06666667
## I 0.10000000 1.00000000 0.20000000
## H 0.06666667 0.20000000 1.00000000
```

```
heatmap.2(dist_mat, scale = "none", trace = "none",
          col = colorpanel(50, "grey75", "red"),
          colsep = 1:4, rowsep = 1:4)
```

An arithmetically decreasing distance is computed as follows. In this case every connected node is by the length of their shortest paths relative to the diameter.

```
dist_mat <- make_distance_graph(graph_structure, absolute = T)
dist_mat
```

```
##     A   C   B   D   E   F   G   I   H
## A 1.0 0.8 0.6 0.6 0.4 0.4 0.2 0.2 0.0
## C 0.8 1.0 0.8 0.8 0.6 0.6 0.4 0.4 0.2
## B 0.6 0.8 1.0 0.6 0.4 0.4 0.2 0.2 0.0
## D 0.6 0.8 0.6 1.0 0.8 0.8 0.6 0.6 0.4
## E 0.4 0.6 0.4 0.8 1.0 0.6 0.4 0.4 0.2
## F 0.4 0.6 0.4 0.8 0.6 1.0 0.8 0.8 0.6
## G 0.2 0.4 0.2 0.6 0.4 0.8 1.0 0.6 0.4
## I 0.2 0.4 0.2 0.6 0.4 0.8 0.6 1.0 0.8
## H 0.0 0.2 0.0 0.4 0.2 0.6 0.4 0.8 1.0
```

```
heatmap.2(dist_mat, scale = "none", trace = "none",
          col = colorpanel(50, "grey75", "red"),
          colsep = 1:4, rowsep = 1:4)
```

##### 3.2.3 Sigma (Σ) matrix

The Sigma (Σ) covariance matrix defines the relationships between the simulated gene distributions. Where the diagonal is one (1), the covariance terms are correlations between each gene. Where possible these are derived from the distance relationships described above. In cases where this is not compatible, the nearest “positive definite” symmetric matrix is computed.

These can be computed directly from an adjacency matrix.

```
#sigma from adj mat
sigma_mat <- make_sigma_mat_graph(graph_structure, 0.8)
sigma_mat
```

```
##     A   C   B   D   E   F   G   I   H
## A 1.0 0.8 0.0 0.0 0.0 0.0 0.0 0.0 0.0
## C 0.8 1.0 0.8 0.8 0.0 0.0 0.0 0.0 0.0
## B 0.0 0.8 1.0 0.0 0.0 0.0 0.0 0.0 0.0
## D 0.0 0.8 0.0 1.0 0.8 0.8 0.0 0.0 0.0
## E 0.0 0.0 0.0 0.8 1.0 0.0 0.0 0.0 0.0
## F 0.0 0.0 0.0 0.8 0.0 1.0 0.8 0.8 0.0
## G 0.0 0.0 0.0 0.0 0.0 0.8 1.0 0.0 0.0
## I 0.0 0.0 0.0 0.0 0.0 0.8 0.0 1.0 0.8
## H 0.0 0.0 0.0 0.0 0.0 0.0 0.0 0.8 1.0
```

```
heatmap.2(sigma_mat, scale = "none", trace = "none",
          col = colorpanel(50, "grey75", "red"),
          colsep = 1:4, rowsep = 1:4)
```

A commonlink matrix can also be used to compute a Σ matrix.

```
#sigma from comm mat
sigma_mat <- make_sigma_mat_graph(graph_structure, 0.8, comm = T)
sigma_mat
```

```
##     A   C   B   D   E   F   G   I   H
## A 1.0 0.0 0.8 0.8 0.0 0.0 0.0 0.0 0.0
## C 0.0 1.0 0.0 0.0 0.8 0.8 0.0 0.0 0.0
## B 0.8 0.0 1.0 0.8 0.0 0.0 0.0 0.0 0.0
## D 0.8 0.0 0.8 1.0 0.0 0.0 0.8 0.8 0.0
## E 0.0 0.8 0.0 0.0 1.0 0.8 0.0 0.0 0.0
## F 0.0 0.8 0.0 0.0 0.8 1.0 0.0 0.0 0.8
## G 0.0 0.0 0.0 0.8 0.0 0.0 1.0 0.8 0.0
## I 0.0 0.0 0.0 0.8 0.0 0.0 0.8 1.0 0.0
## H 0.0 0.0 0.0 0.0 0.0 0.8 0.0 0.0 1.0
```

```
heatmap.2(sigma_mat, scale = "none", trace = "none",
          col = colorpanel(50, "grey75", "red"),
          colsep = 1:4, rowsep = 1:4)
```

It is recommended to compute the distance relationships and use these. This is supported with the built-in functions. For instance Σ from the geometrically computed distances.

```
# sigma from geometric distance matrix
make_sigma_mat_dist_graph(graph_structure, 0.8, absolute = F)
```

```
##           A         C         B         D         E         F
## A 1.0000000 0.8000000 0.4000000 0.4000000 0.2666667 0.2666667
## C 0.8000000 1.0000000 0.8000000 0.8000000 0.4000000 0.4000000
## B 0.4000000 0.8000000 1.0000000 0.4000000 0.2666667 0.2666667
## D 0.4000000 0.8000000 0.4000000 1.0000000 0.8000000 0.8000000
## E 0.2666667 0.4000000 0.2666667 0.8000000 1.0000000 0.4000000
## F 0.2666667 0.4000000 0.2666667 0.8000000 0.4000000 1.0000000
## G 0.2000000 0.2666667 0.2000000 0.4000000 0.2666667 0.8000000
## I 0.2000000 0.2666667 0.2000000 0.4000000 0.2666667 0.8000000
## H 0.1600000 0.2000000 0.1600000 0.2666667 0.2000000 0.4000000
##           G         I         H
## A 0.2000000 0.2000000 0.1600000
## C 0.2666667 0.2666667 0.2000000
## B 0.2000000 0.2000000 0.1600000
## D 0.4000000 0.4000000 0.2666667
## E 0.2666667 0.2666667 0.2000000
## F 0.8000000 0.8000000 0.4000000
## G 1.0000000 0.4000000 0.2666667
## I 0.4000000 1.0000000 0.8000000
## H 0.2666667 0.8000000 1.0000000
```

```
heatmap.2(make_sigma_mat_dist_graph(graph_structure, 0.8, absolute = F),
          scale = "none", trace = "none",
          col = colorpanel(50, "grey75", "red"),
          colsep = 1:4, rowsep = 1:4)
```

Σ can also be computed for arithmetically computed distances.

```
# sigma from absolute distance matrix
sigma_mat <- make_sigma_mat_dist_graph(graph_structure, 0.8, absolute = T)
heatmap.2(sigma_mat, scale = "none", trace = "none", 
          col = colorpanel(50, "grey75", "red"),
          colsep = 1:4, rowsep = 1:4)
```

##### 3.2.4 Simulated expression and observed correlation

Here we generate the final simulated expression dataset. Note that none of the prior steps are required. These are called internalled as needed.

For example, the adjacency matrix is derived to generate the following dataset. Note that the nearest positive definite matrix is required for the Σ matrix in this case.

```
#simulate expression data
#adj mat
expr <- generate_expression(100, graph_structure, cor = 0.8,
                            mean = 0, comm = F)
heatmap.2(expr, scale = "none", trace = "none",
          col = bluered(50), colsep = 1:4, rowsep = 1:4)
```

```
heatmap.2(cor(t(expr)), scale = "none", trace = "none",
          col = colorpanel(50, "grey75", "red"),
          colsep = 1:4, rowsep = 1:4)
```

Here we compute a simulated dataset based on common links shared to other nodes.

```
#comm mat
expr <- generate_expression(100, graph_structure, cor = 0.8, mean = 0, comm =T)
```

```
## Warning in generate_expression(100, graph_structure, cor = 0.8, mean
## = 0, : sigma matrix was not positive definite, nearest approximation
## used.
```

```
heatmap.2(expr, scale = "none", trace = "none",
          col = bluered(50), colsep = 1:4, rowsep = 1:4)
```

```
heatmap.2(cor(t(expr)), scale = "none", trace = "none",
          col = colorpanel(50, "grey75", "red"),
          colsep = 1:4, rowsep = 1:4)
```

Here we use relative distance (relationships are geometrically weighted to the diameter).

```
# relative dist
expr <- generate_expression(100, graph_structure, cor = 0.8,mean = 0,
                            comm = F, dist = T, absolute = F)
```

```
## Warning in generate_expression(100, graph_structure, cor = 0.8, mean
## = 0, : sigma matrix was not positive definite, nearest approximation
## used.
```

```
heatmap.2(expr, scale = "none", trace = "none", col = bluered(50),
          colsep = 1:4, rowsep = 1:4)
```

```
heatmap.2(cor(t(expr)), scale = "none", trace = "none",
          col = colorpanel(50, "grey75", "red"),
          colsep = 1:4, rowsep = 1:4)
```

Here we use absolute distance (relationships are arithmetically weighted to the diameter).

```
#absolute dist
expr <- generate_expression(100, graph_structure, cor = 0.8, mean = 0,
                            comm = F, dist = T, absolute = T) 
heatmap.2(expr, scale = "none", trace = "none", col = bluered(50),
          colsep = 1:4, rowsep = 1:4)
```

```
heatmap.2(cor(t(expr)), scale = "none", trace = "none",
           col = bluered(50), colsep = 1:4, rowsep = 1:4)
```

#### 3.3 Summary

In summary, we compute the following expression dataset but on these underlying relationships in the graph structure. Here we use geometrically decreasing correlations between more distant nodes in the network. This code is a complete example to compute the following plots.

```
# activating graph
state <- rep(1, length(E(graph_structure)))
plot_directed(graph_structure, state=state,
              layout = layout.kamada.kawai,
              cex.node=2, cex.arrow=4, arrow_clip = 0.2)
mtext(text = "(a) Activating pathway structure", side=1,
      line=3.5, at=0.075, adj=0.5, cex=1.75)
box()
#plot relationship matrix
heatmap.2(make_distance_graph(graph_structure, absolute = FALSE),
          scale = "none", trace = "none",
          col = colorpanel(50, "white", "red"),
          colsep = 1:length(V(graph_structure)),
          rowsep = 1:length(V(graph_structure)))
mtext(text = "(b) Relationship matrix", side=1,
      line=3.5, at=0, adj=0.5, cex=1.75)
#plot sigma matrix
sigma_mat <- make_sigma_mat_dist_graph(graph_structure,
               cor = 0.8, absolute = FALSE)
heatmap.2(sigma_mat, scale = "none", trace = "none",
          col = colorpanel(50, "white", "red"),
          colsep = 1:length(V(graph_structure)),
          rowsep = 1:length(V(graph_structure)))
mtext(text = expression(paste("(c) ", Sigma, " matrix")), side=1,
      line=3.5, at=0, adj=0.5, cex=1.75)
#simulated data
expr <- generate_expression(100, graph_structure, cor = 0.8, mean = 0,
          comm = FALSE, dist =TRUE, absolute = FALSE, state = state)
#plot simulated correlations
heatmap.2(cor(t(expr)), scale = "none", trace = "none",
          col = colorpanel(50, "white", "red"),
          colsep = 1:length(V(graph_structure)),
          rowsep = 1:length(V(graph_structure)))
mtext(text = "(d) Simulated correlation", side=1,
      line=3.5, at=0, adj=0.5, cex=1.75)
#plot simulated expression data
heatmap.2(expr, scale = "none", trace = "none", col = bluered(50),
          colsep = 1:length(V(graph_structure)),
          rowsep = 1:length(V(graph_structure)), labCol = "")
mtext(text = "samples", side=1, line=1.5, at=0.2, adj=0.5, cex=1.5)
mtext(text = "genes", side=4, line=1, at=-0.4, adj=0.5, cex=1.5)
mtext(text = "(e) Simulated expression data (log scale)", side=1,
      line=3.5, at=0, adj=0.5, cex=1.75)
```

##### 3.3.1 Inhibiting relationships

Here we simulate the same graph structure with inhibiting edges but passing the `"state"` parameter. This takes one argument for each each to identify which are inhibitory (as documented).

```
# activating graph
state <- state <- c(1, 1, -1, 1, 1, 1, 1, -1)
plot_directed(graph_structure, state=state, layout = layout.kamada.kawai,
              cex.node=2, cex.arrow=4, arrow_clip = 0.2)
mtext(text = "(a) Activating pathway structure", side=1,
      line=3.5, at=0.075, adj=0.5, cex=1.75)
box()
#plot relationship matrix
heatmap.2(make_distance_graph(graph_structure, absolute = FALSE),
          scale = "none", trace = "none",
          col = colorpanel(50, "white", "red"),
          colsep = 1:length(V(graph_structure)),
          rowsep = 1:length(V(graph_structure)))
mtext(text = "(b) Relationship matrix", side=1,
      line=3.5, at=0, adj=0.5, cex=1.75)
#plot sigma matrix
heatmap.2(make_sigma_mat_dist_graph(graph_structure, state,
                                    cor = 0.8, absolute = FALSE),
          scale = "none", trace = "none",
          col = colorpanel(50, "blue", "white", "red"),
          colsep = 1:length(V(graph_structure)),
          rowsep = 1:length(V(graph_structure)))
mtext(text = expression(paste("(c) ", Sigma, " matrix")), side=1,
      line=3.5, at=0, adj=0.5, cex=1.75)
#simulated data
expr <- generate_expression(100, graph_structure, state,
                            cor = 0.8, mean = 0,
                            comm = FALSE, dist =TRUE,
                            absolute = FALSE, state = state)
#plot simulated correlations
heatmap.2(cor(t(expr)), scale = "none", trace = "none",
          col = colorpanel(50, "blue", "white", "red"),
          colsep = 1:length(V(graph_structure)),
          rowsep = 1:length(V(graph_structure)))
mtext(text = "(d) Simulated correlation", side=1,
      line=3.5, at=0, adj=0.5, cex=1.75)
#plot simulated expression data
heatmap.2(expr, scale = "none", trace = "none", col = bluered(50),
          colsep = 1:length(V(graph_structure)),
          rowsep = 1:length(V(graph_structure)), labCol = "")
mtext(text = "samples", side=1, line=1.5, at=0.2, adj=0.5, cex=1.5)
mtext(text = "genes", side=4, line=1, at=-0.4, adj=0.5, cex=1.5)
mtext(text = "(e) Simulated expression data (log scale)", side=1,
      line=3.5, at=0, adj=0.5, cex=1.75)
```

### 4 Examples

#### 4.1 Toy examples

Here we give some toy examples to show how the simulations behave in simple cases. This serves to understand how modules within a larger graph will translate to correlations in the final simulated datasets.

##### 4.1.1 Diverging branches

```
graph_diverging_edges <- rbind(c("A", "B"), c("B", "C"), c("B", "D"))
graph_diverging <- graph.edgelist(graph_diverging_edges, directed = T)

# activating graph
state <- rep(1, length(E(graph_diverging)))
plot_directed(graph_diverging, state=state,
              layout = layout.kamada.kawai,
              cex.node=2, cex.arrow=4, arrow_clip = 0.2)
mtext(text = "(a) Activating pathway structure", side=1,
      line=3.5, at=0.075, adj=0.5, cex=1.75)
box()
#plot relationship matrix
heatmap.2(make_distance_graph(graph_diverging, absolute = FALSE),
          scale = "none", trace = "none",
          col = colorpanel(50, "white", "red"),
          colsep = 1:length(V(graph_diverging)),
          rowsep = 1:length(V(graph_diverging)))
mtext(text = "(b) Relationship matrix", side=1,
      line=3.5, at=0, adj=0.5, cex=1.75)
#plot sigma matrix
heatmap.2(make_sigma_mat_dist_graph(graph_diverging,
                                    cor = 0.8, absolute = FALSE),
          scale = "none", trace = "none",
          col = colorpanel(50, "white", "red"),
          colsep = 1:length(V(graph_diverging)),
          rowsep = 1:length(V(graph_diverging)))
mtext(text = expression(paste("(c) ", Sigma, " matrix")), side=1,
      line=3.5, at=0, adj=0.5, cex=1.75)
#simulated data
expr <- generate_expression(100, graph_diverging,
                            cor = 0.8, mean = 0,
                            comm = FALSE, dist =TRUE,
                            absolute = FALSE, state = state)
#plot simulated correlations
heatmap.2(cor(t(expr)), scale = "none", trace = "none",
          col = colorpanel(50, "white", "red"),
          colsep = 1:length(V(graph_diverging)),
          rowsep = 1:length(V(graph_diverging)))
mtext(text = "(d) Simulated correlation", side=1,
      line=3.5, at=0, adj=0.5, cex=1.75)
#plot simulated expression data
heatmap.2(expr, scale = "none", trace = "none",col = bluered(50),
          colsep = 1:length(V(graph_diverging)),
          rowsep = 1:length(V(graph_diverging)), labCol = "")
mtext(text = "samples", side=1, line=1.5, at=0.2, adj=0.5, cex=1.5)
mtext(text = "genes", side=4, line=1, at=-0.4, adj=0.5, cex=1.5)
mtext(text = "(e) Simulated expression data (log scale)", side=1,
      line=3.5, at=0, adj=0.5, cex=1.75)
```

###### 4.1.1.1 Inhibiting relationships

Here we simulate the same graph structure with inhibiting edges but passing the `"state"` parameter. This takes one argument for each each to identify which are inhibitory (as documented).

```
# activating graph
state <- state <- c(1, 1, -1)
plot_directed(graph_diverging, state=state, layout = layout.kamada.kawai,
              cex.node=2, cex.arrow=4, arrow_clip = 0.2)
mtext(text = "(a) Activating pathway structure", side=1,
      line=3.5, at=0.075, adj=0.5, cex=1.75)
box()
#plot relationship matrix
heatmap.2(make_distance_graph(graph_diverging, absolute = FALSE),
          scale = "none", trace = "none",
          col = colorpanel(50, "white", "red"),
          colsep = 1:length(V(graph_diverging)),
          rowsep = 1:length(V(graph_diverging)))
mtext(text = "(b) Relationship matrix", side=1,
      line=3.5, at=0, adj=0.5, cex=1.75)
#plot sigma matrix
heatmap.2(make_sigma_mat_dist_graph(graph_diverging, state,
                                    cor = 0.8, absolute = FALSE),
          scale = "none", trace = "none",
          col = colorpanel(50, "blue", "white", "red"),
          colsep = 1:length(V(graph_diverging)),
          rowsep = 1:length(V(graph_diverging)))
mtext(text = expression(paste("(c) ", Sigma, " matrix")), side=1,
      line=3.5, at=0, adj=0.5, cex=1.75)
#simulated data
expr <- generate_expression(100, graph_diverging, state,
                            cor = 0.8, mean = 0,
                            comm = FALSE, dist =TRUE,
                            absolute = FALSE, state = state)
#plot simulated correlations
heatmap.2(cor(t(expr)), scale = "none", trace = "none",
          col = colorpanel(50, "blue", "white", "red"),
          colsep = 1:length(V(graph_diverging)),
          rowsep = 1:length(V(graph_diverging)))
mtext(text = "(d) Simulated correlation", side=1,
      line=3.5, at=0, adj=0.5, cex=1.75)
#plot simulated expression data
heatmap.2(expr, scale = "none", trace = "none", col = bluered(50),
          colsep = 1:length(V(graph_diverging)),
          rowsep = 1:length(V(graph_diverging)), labCol = "")
mtext(text = "samples", side=1, line=1.5, at=0.2, adj=0.5, cex=1.5)
mtext(text = "genes", side=4, line=1, at=-0.4, adj=0.5, cex=1.5)
mtext(text = "(e) Simulated expression data (log scale)", side=1,
      line=3.5, at=0, adj=0.5, cex=1.75)
```

##### 4.1.2 Converging branches

```
graph_converging_edges <- rbind(c("C", "E"), c("D", "E"), c("E", "F"))
graph_converging <- graph.edgelist(graph_converging_edges, directed = T)

# activating graph
state <- rep(1, length(E(graph_converging)))
plot_directed(graph_converging, state=state,
              layout = layout.kamada.kawai,
              cex.node=2, cex.arrow=4, arrow_clip = 0.2)
mtext(text = "(a) Activating pathway structure", side=1,
      line=3.5, at=0.075, adj=0.5, cex=1.75)
box()
#plot relationship matrix
heatmap.2(make_distance_graph(graph_converging, absolute = FALSE),
          scale = "none", trace = "none",
          col = colorpanel(50, "white", "red"),
          colsep = 1:length(V(graph_converging)),
          rowsep = 1:length(V(graph_converging)))
mtext(text = "(b) Relationship matrix", side=1,
      line=3.5, at=0, adj=0.5, cex=1.75)
#plot sigma matrix
heatmap.2(make_sigma_mat_dist_graph(graph_converging, cor = 0.8,
                                    absolute = FALSE),
          scale = "none", trace = "none",
          col = colorpanel(50, "white", "red"),
          colsep = 1:length(V(graph_converging)),
          rowsep = 1:length(V(graph_converging)))
mtext(text = expression(paste("(c) ", Sigma, " matrix")), side=1,
      line=3.5, at=0, adj=0.5, cex=1.75)
#simulated data
expr <- generate_expression(100, graph_converging,
                            cor = 0.8, mean = 0,
                            comm = FALSE, dist =TRUE,
                            absolute = FALSE, state = state)
#plot simulated correlations
heatmap.2(cor(t(expr)), scale = "none", trace = "none",
          col = colorpanel(50, "white", "red"),
          colsep = 1:length(V(graph_converging)),
          rowsep = 1:length(V(graph_converging)))
mtext(text = "(d) Simulated correlation", side=1,
      line=3.5, at=0, adj=0.5, cex=1.75)
#plot simulated expression data
heatmap.2(expr, scale = "none", trace = "none", col = bluered(50),
          colsep = 1:length(V(graph_converging)),
          rowsep = 1:length(V(graph_converging)), labCol = "")
mtext(text = "samples", side=1, line=1.5, at=0.2, adj=0.5, cex=1.5)
mtext(text = "genes", side=4, line=1, at=-0.4, adj=0.5, cex=1.5)
mtext(text = "(e) Simulated expression data (log scale)", side=1,
      line=3.5, at=0, adj=0.5, cex=1.75)
```

###### 4.1.2.1 Inhibiting relationships

Here we simulate the same graph structure with inhibiting edges but passing the `"state"` parameter. This takes one argument for each each to identify which are inhibitory (as documented).

```
# activating graph
state <- state <- c(-1, 1, -1)
plot_directed(graph_converging, state=state,
              layout = layout.kamada.kawai,
              cex.node=2, cex.arrow=4, arrow_clip = 0.2)
mtext(text = "(a) Activating pathway structure", side=1,
      line=3.5, at=0.075, adj=0.5, cex=1.75)
box()
#plot relationship matrix
heatmap.2(make_distance_graph(graph_converging, absolute = FALSE),
          scale = "none", trace = "none",
          col = colorpanel(50, "white", "red"),
          colsep = 1:length(V(graph_converging)),
          rowsep = 1:length(V(graph_converging)))
mtext(text = "(b) Relationship matrix", side=1,
      line=3.5, at=0, adj=0.5, cex=1.75)
#plot sigma matrix
heatmap.2(make_sigma_mat_dist_graph(graph_converging, state,
                                    cor = 0.8, absolute = FALSE),
          scale = "none", trace = "none",
          col = colorpanel(50, "blue", "white", "red"),
          colsep = 1:length(V(graph_converging)),
          rowsep = 1:length(V(graph_converging)))
mtext(text = expression(paste("(c) ", Sigma, " matrix")), side=1,
      line=3.5, at=0, adj=0.5, cex=1.75)
#simulated data
expr <- generate_expression(100, graph_converging, state,
                            cor = 0.8, mean = 0,
                            comm = FALSE, dist =TRUE,
                            absolute = FALSE, state = state)
#plot simulated correlations
heatmap.2(cor(t(expr)), scale = "none", trace = "none",
          col = colorpanel(50, "blue", "white", "red"),
          colsep = 1:length(V(graph_converging)),
          rowsep = 1:length(V(graph_converging)))
mtext(text = "(d) Simulated correlation", side=1,
      line=3.5, at=0, adj=0.5, cex=1.75)
#plot simulated expression data
heatmap.2(expr, scale = "none", trace = "none", col = bluered(50),
          colsep = 1:length(V(graph_converging)),
          rowsep = 1:length(V(graph_converging)), labCol = "")
mtext(text = "samples", side=1, line=1.5, at=0.2, adj=0.5, cex=1.5)
mtext(text = "genes", side=4, line=1, at=-0.4, adj=0.5, cex=1.5)
mtext(text = "(e) Simulated expression data (log scale)", side=1,
      line=3.5, at=0, adj=0.5, cex=1.75)
```

##### 4.1.3 Reconnecting paths

```
graph_reconnecting_edges <- rbind(c("A", "B"), c("B", "C"), c("B", "D"),
                                  c("C", "E"), c("D", "E"), c("E", "F"))
graph_reconnecting <- graph.edgelist(graph_reconnecting_edges, directed = T)

# activating graph
state <- rep(1, length(E(graph_reconnecting)))
plot_directed(graph_reconnecting, state=state,
              layout = layout.kamada.kawai,
              cex.node=2, cex.arrow=4, arrow_clip = 0.2)
mtext(text = "(a) Activating pathway structure", side=1,
      line=3.5, at=0.075, adj=0.5, cex=1.75)
box()
#plot relationship matrix
heatmap.2(make_distance_graph(graph_reconnecting, absolute = FALSE),
          scale = "none", trace = "none",
          col = colorpanel(50, "white", "red"),
          colsep = 1:length(V(graph_reconnecting)),
          rowsep = 1:length(V(graph_reconnecting)))
mtext(text = "(b) Relationship matrix", side=1,
      line=3.5, at=0, adj=0.5, cex=1.75)
#plot sigma matrix
heatmap.2(make_sigma_mat_dist_graph(graph_reconnecting, cor = 0.8,
                                    absolute = FALSE),
          scale = "none", trace = "none",
          col = colorpanel(50, "white", "red"),
          colsep = 1:length(V(graph_reconnecting)),
          rowsep = 1:length(V(graph_reconnecting)))
mtext(text = expression(paste("(c) ", Sigma, " matrix")), side=1,
      line=3.5, at=0, adj=0.5, cex=1.75)
#simulated data
expr <- generate_expression(100, graph_reconnecting,
                            cor = 0.8, mean = 0,
                            comm = FALSE, dist =TRUE,
                            absolute = FALSE, state = state)
#plot simulated correlations
heatmap.2(cor(t(expr)), scale = "none", trace = "none",
          col = colorpanel(50, "white", "red"),
          colsep = 1:length(V(graph_reconnecting)),
          rowsep = 1:length(V(graph_reconnecting)))
mtext(text = "(d) Simulated correlation", side=1,
      line=3.5, at=0, adj=0.5, cex=1.75)
#plot simulated expression data
heatmap.2(expr, scale = "none", trace = "none", col = bluered(50),
          colsep = 1:length(V(graph_reconnecting)),
          rowsep = 1:length(V(graph_reconnecting)), labCol = "")
mtext(text = "samples", side=1, line=1.5, at=0.2, adj=0.5, cex=1.5)
mtext(text = "genes", side=4, line=1, at=-0.4, adj=0.5, cex=1.5)
mtext(text = "(e) Simulated expression data (log scale)", side=1,
      line=3.5, at=0, adj=0.5, cex=1.75)
```

###### 4.1.3.1 Inhibiting relationships

Here we simulate the same graph structure with inhibiting edges but passing the `"state"` parameter. This takes one argument for each each to identify which are inhibitory (as documented).

```
# activating graph
state <- state <- c(1, 1, -1, -1, 1, 1, 1, 1)
plot_directed(graph_reconnecting, state=state,
              layout = layout.kamada.kawai,
              cex.node=2, cex.arrow=4, arrow_clip = 0.2)
mtext(text = "(a) Activating pathway structure", side=1,
      line=3.5, at=0.075, adj=0.5, cex=1.75)
box()
#plot relationship matrix
heatmap.2(make_distance_graph(graph_reconnecting, absolute = FALSE),
          scale = "none", trace = "none",
          col = colorpanel(50, "white", "red"),
          colsep = 1:length(V(graph_reconnecting)),
          rowsep = 1:length(V(graph_reconnecting)))
mtext(text = "(b) Relationship matrix", side=1, line=3.5, at=0, adj=0.5, cex=1.75)
#plot sigma matrix
heatmap.2(make_sigma_mat_dist_graph(graph_reconnecting, state,
                                    cor = 0.8, absolute = FALSE),
          scale = "none", trace = "none",
          col = colorpanel(50, "blue", "white", "red"),
          colsep = 1:length(V(graph_reconnecting)),
          rowsep = 1:length(V(graph_reconnecting)))
mtext(text = expression(paste("(c) ", Sigma, " matrix")), side=1,
      line=3.5, at=0, adj=0.5, cex=1.75)
#simulated data
expr <- generate_expression(100, graph_reconnecting, state,
                            cor = 0.8, mean = 0,
                            comm = FALSE, dist =TRUE,
                            absolute = FALSE, state = state)
#plot simulated correlations
heatmap.2(cor(t(expr)), scale = "none", trace = "none",
          col = colorpanel(50, "blue", "white", "red"),
          colsep = 1:length(V(graph_reconnecting)),
          rowsep = 1:length(V(graph_reconnecting)))
mtext(text = "(d) Simulated correlation", side=1,
      line=3.5, at=0, adj=0.5, cex=1.75)
#plot simulated expression data
heatmap.2(expr, scale = "none", trace = "none", col = bluered(50),
          colsep = 1:length(V(graph_reconnecting)),
          rowsep = 1:length(V(graph_reconnecting)), labCol = "")
mtext(text = "samples", side=1, line=1.5, at=0.2, adj=0.5, cex=1.5)
mtext(text = "genes", side=4, line=1, at=-0.4, adj=0.5, cex=1.5)
mtext(text = "(e) Simulated expression data (log scale)", side=1,
      line=3.5, at=0, adj=0.5, cex=1.75)
```

#### 4.2 Empirical examples

Next we demonstrate the simulation procedure based on real biological pathways from the “Reactome” database (Croft *et al*. 2014). We can import these from the `data` directory included with this package. These graphs are given for examples and convenience.

##### 4.2.1 Kinase pathways

The following pathways are treated as all relationships are activating.

###### 4.2.1.1 RAF/MAP kinase cascade

Here we generate simulated data for the RAF/MAP kinase cascade pathway.

```
RAF_MAP_graph <- identity(RAF_MAP_graph)

# activating graph
state <- rep(1, length(E(RAF_MAP_graph)))
plot_directed(RAF_MAP_graph, state=state,
              layout = layout.kamada.kawai,
              cex.node=2, cex.arrow=4, arrow_clip = 0.2,
              col.arrow = "#00A9FF", fill.node = "lightblue")
mtext(text = "(a) Activating pathway structure", side=1,
      line=3.5, at=0.075, adj=0.5, cex=1.75)
box()
#plot relationship matrix
heatmap.2(make_distance_graph(RAF_MAP_graph, absolute = FALSE),
          scale = "none", trace = "none",
          col = colorpanel(50, "white", "red"),
          colsep = 1:length(V(RAF_MAP_graph)),
          rowsep = 1:length(V(RAF_MAP_graph)))
mtext(text = "(b) Relationship matrix", side=1,
      line=3.5, at=0, adj=0.5, cex=1.75)
#plot sigma matrix
heatmap.2(make_sigma_mat_dist_graph(RAF_MAP_graph, cor = 0.8,
                                    absolute = FALSE),
          scale = "none", trace = "none",
          col = colorpanel(50, "white", "red"),
          colsep = 1:length(V(RAF_MAP_graph)),
          rowsep = 1:length(V(RAF_MAP_graph)))
mtext(text = expression(paste("(c) ", Sigma, " matrix")), side=1,
      line=3.5, at=0, adj=0.5, cex=1.75)
#simulated data
expr <- generate_expression(100, RAF_MAP_graph,
                            cor = 0.8, mean = 0,
                            comm = FALSE, dist =TRUE,
                            absolute = FALSE, state = state)
#plot simulated correlations
heatmap.2(cor(t(expr)), scale = "none", trace = "none",
          col = colorpanel(50, "white", "red"),
          colsep = 1:length(V(RAF_MAP_graph)),
          rowsep = 1:length(V(RAF_MAP_graph)))
mtext(text = "(d) Simulated correlation", side=1,
      line=3.5, at=0, adj=0.5, cex=1.75)
#plot simulated expression data
heatmap.2(expr, scale = "none", trace = "none", col = bluered(50),
          colsep = 1:length(V(RAF_MAP_graph)),
          rowsep = 1:length(V(RAF_MAP_graph)), labCol = "")
mtext(text = "samples", side=1, line=1.5, at=0.2, adj=0.5, cex=1.5)
mtext(text = "genes", side=4, line=1, at=-0.4, adj=0.5, cex=1.5)
mtext(text = "(e) Simulated expression data (log scale)", side=1,
      line=3.5, at=0, adj=0.5, cex=1.75)
```

###### 4.2.1.2 Phosphoinositide-3-kinase cascade

Here we generate simulated data for the phosphoinositide-3-kinase (Pi3K) cascade pathway.

```
graph <- identity(Pi3K_graph)

# activating graph
state <- rep(1, length(E(Pi3K_graph)))
plot_directed(Pi3K_graph, state=state,
              layout = layout.kamada.kawai,
              cex.node=2, cex.arrow=4, arrow_clip = 0.2,
              col.arrow = "#00A9FF", fill.node = "lightblue")
mtext(text = "(a) Activating pathway structure", side=1,
      line=3.5, at=0.075, adj=0.5, cex=1.75)
box()
#plot relationship matrix
heatmap.2(make_distance_graph(Pi3K_graph, absolute = FALSE),
          scale = "none", trace = "none",
          col = colorpanel(50, "white", "red"),
          colsep = 1:length(V(Pi3K_graph)),
          rowsep = 1:length(V(Pi3K_graph)))
mtext(text = "(b) Relationship matrix", side=1,
      line=3.5, at=0, adj=0.5, cex=1.75)
#plot sigma matrix
heatmap.2(make_sigma_mat_dist_graph(Pi3K_graph, cor = 0.8,
                                    absolute = FALSE),
          scale = "none", trace = "none",
          col = colorpanel(50, "white", "red"),
          colsep = 1:length(V(Pi3K_graph)),
          rowsep = 1:length(V(Pi3K_graph)))
mtext(text = expression(paste("(c) ", Sigma, " matrix")), side=1,
      line=3.5, at=0, adj=0.5, cex=1.75)
#simulated data
expr <- generate_expression(100, Pi3K_graph, 
                            cor = 0.8, mean = 0,
                            comm = FALSE, dist =TRUE,
                            absolute = FALSE, state = state)
#plot simulated correlations
heatmap.2(cor(t(expr)), scale = "none", trace = "none",
          col = colorpanel(50, "white", "red"),
          colsep = 1:length(V(Pi3K_graph)),
          rowsep = 1:length(V(Pi3K_graph)))
mtext(text = "(d) Simulated correlation", side=1, line=3.5, at=0, adj=0.5, cex=1.75)
#plot simulated expression data
heatmap.2(expr, scale = "none", trace = "none", col = bluered(50),
          colsep = 1:length(V(Pi3K_graph)),
          rowsep = 1:length(V(Pi3K_graph)), labCol = "")
mtext(text = "samples", side=1, line=1.5, at=0.2, adj=0.5, cex=1.5)
mtext(text = "genes", side=4, line=1, at=-0.4, adj=0.5, cex=1.5)
mtext(text = "(e) Simulated expression data (log scale)", side=1,
      line=3.5, at=0, adj=0.5, cex=1.75)
```

##### 4.2.2 Pathways with repression

###### 4.2.2.1 The Pi3K/AKT pathway

Here we generate simulated data for the phosphoinositide-3-kinase activation of Protein kinase B (PKB) cascade (also known as Pi3k/AKT) pathway. States are imported as edge attributes from the imported graph.

```
Pi3K_AKT_graph <- identity(Pi3K_AKT_graph)
Pi3K_AKT_graph <- simplify(Pi3K_AKT_graph,
      edge.attr.comb = function(x){
         ifelse(any(x %in% list(-1, 2, "inhibiting", "inhibition")), 2, 1)
        })
Pi3K_AKT_graph <- simplify(Pi3K_AKT_graph, edge.attr.comb = "first")
edge_properties <- E(Pi3K_AKT_graph)$state

# activating graph
#state <- rep(1, length(E(Pi3K_AKT_graph)))
plot_directed(Pi3K_AKT_graph, state = edge_properties,
              layout = layout.kamada.kawai,
              cex.node=2, cex.arrow=4, arrow_clip = 0.2,
              col.arrow = c(alpha("navyblue", 0.25),
                            alpha("red", 0.25))[edge_properties],
              fill.node = "lightblue")
mtext(text = "(a) Activating pathway structure", side=1,
      line=3.5, at=0.075, adj=0.5, cex=1.75)
box()
#plot relationship matrix
heatmap.2(make_distance_graph(Pi3K_AKT_graph, absolute = FALSE),
          scale = "none", trace = "none",
          col = colorpanel(50, "white", "red"))
mtext(text = "(b) Relationship matrix", side=1, line=3.5, at=0, adj=0.5, cex=1.75)
#plot sigma matrix
heatmap.2(make_sigma_mat_dist_graph(Pi3K_AKT_graph, state = edge_properties,
                                    cor = 0.8, absolute = FALSE),
          scale = "none", trace = "none",
          col = colorpanel(50, "blue", "white", "red"))
mtext(text = expression(paste("(c) ", Sigma, " matrix")), side=1,
      line=3.5, at=0, adj=0.5, cex=1.75)
#simulated data
expr <- generate_expression(100, Pi3K_AKT_graph, cor = 0.8, mean = 0,
                            comm = FALSE, dist =TRUE, absolute = FALSE,
                            state = edge_properties)
#plot simulated correlations
heatmap.2(cor(t(expr)), scale = "none", trace = "none",
          col = colorpanel(50, "blue", "white", "red"))
mtext(text = "(d) Simulated correlation", side=1,
      line=3.5, at=0, adj=0.5, cex=1.75)
#plot simulated expression data
heatmap.2(expr, scale = "none", trace = "none",
          col = bluered(50), labCol = "")
mtext(text = "samples", side=1, line=1.5, at=0.2, adj=0.5, cex=1.5)
mtext(text = "genes", side=4, line=1, at=-0.4, adj=0.5, cex=1.5)
mtext(text = "(e) Simulated expression data (log scale)", side=1,
      line=3.5, at=0, adj=0.5, cex=1.75)
```

###### 4.2.2.2 The TGFβ-Smad pathway

Here we generate simulated data for the TGFβ-Smad gene regulatory pathway with inhibitions known. States are imported as edge attributes from the imported graph.

```
TGFBeta_Smad_graph <- identity(TGFBeta_Smad_graph)
edge_properties <- E(TGFBeta_Smad_graph)$state
plot_directed(TGFBeta_Smad_graph, state = edge_properties,
              col.arrow = c(alpha("navyblue", 0.25),
                            alpha("red", 0.25))[edge_properties],
              fill.node = c("lightblue"))
```

```
TGFBeta_Smad_graph <- identity(TGFBeta_Smad_graph)
edge_properties <- E(TGFBeta_Smad_graph)$state

# activating graph
plot_directed(TGFBeta_Smad_graph, state = edge_properties,
              layout = layout.kamada.kawai,
              cex.node=2, cex.arrow=4, arrow_clip = 0.2,
              col.arrow = c(alpha("navyblue", 0.25),
                            alpha("red", 0.25))[edge_properties],
              fill.node = "lightblue")
mtext(text = "(a) Activating pathway structure", side=1,
      line=3.5, at=0.075, adj=0.5, cex=1.75)
box()
#plot relationship matrix
heatmap.2(make_distance_graph(TGFBeta_Smad_graph, absolute = FALSE),
          scale = "none", trace = "none",
          col = colorpanel(50, "white", "red"),
          colsep = 1:length(V(TGFBeta_Smad_graph)),
          rowsep = 1:length(V(TGFBeta_Smad_graph)))
mtext(text = "(b) Relationship matrix", side=1,
      line=3.5, at=0, adj=0.5, cex=1.75)
#plot sigma matrix
heatmap.2(make_sigma_mat_dist_graph(TGFBeta_Smad_graph,
                                    state = edge_properties,
                                    cor = 0.8, absolute = FALSE),
          scale = "none", trace = "none",
          col = colorpanel(50, "blue", "white", "red"),
          colsep = 1:length(V(TGFBeta_Smad_graph)),
          rowsep = 1:length(V(TGFBeta_Smad_graph)))
mtext(text = expression(paste("(c) ", Sigma, " matrix")), side=1,
      line=3.5, at=0, adj=0.5, cex=1.75)
#simulated data
expr <- generate_expression(100, TGFBeta_Smad_graph,
                            cor = 0.8, mean = 0,
                            comm = FALSE, dist =TRUE,
                            absolute = FALSE, state = edge_properties)
#plot simulated correlations
heatmap.2(cor(t(expr)), scale = "none", trace = "none",
          col = colorpanel(50, "blue", "white", "red"),
          colsep = 1:length(V(TGFBeta_Smad_graph)),
          rowsep = 1:length(V(TGFBeta_Smad_graph)))
mtext(text = "(d) Simulated correlation", side=1,
      line=3.5, at=0, adj=0.5, cex=1.75)
#plot simulated expression data
heatmap.2(expr, scale = "none", trace = "none", col = bluered(50),
          colsep = 1:length(V(TGFBeta_Smad_graph)),
          rowsep = 1:length(V(TGFBeta_Smad_graph)), labCol = "")
mtext(text = "samples", side=1, line=1.5, at=0.2, adj=0.5, cex=1.5)
mtext(text = "genes", side=4, line=1, at=-0.4, adj=0.5, cex=1.5)
mtext(text = "(e) Simulated expression data (log scale)", side=1,
      line=3.5, at=0, adj=0.5, cex=1.75)
```

### 5 Summary

Here we have demonstrated that simulated datasets can be generated based on graph structures. These can be either constructed networks for modelling purposes or empirical networks such as those from curated biological databases. Various parameters are available and described in the documentation. You can alter these parameters in the examples given here to see the impact they have on the final network. It is encouraged to try different parameters and examine the results carefully, in addition to carefully considering which assumptions are appropriate for your investigations. A model or simulation is never correct, it is a tool to test your assumptions and find weaknesses in your technique, consider which conditions could your method struggle with and model these. Pathway structure in particular should be considered in biological datasets as correlations within a pathway can lead to false positive results and confounding.

The intended application for this package is modelling RNA-Seq gene expression data. However, other applications are encouraged, provided that they require multivariate normal simulations based on relationships in graph structures.

---

#### 5.1 Citation

If you use this package, please cite it where appropriate to recognise the efforts of the developers.

```
citation("graphsim")
```

```
## 
## To cite package 'graphsim' in publications use:
## 
##   S. Thomas Kelly and Michael A. Black (2020). graphsim:
##   Simulate Expression data from igraph networks. R package
##   version 1.0.0.
##   https://github.com/TomKellyGenetics/graphsim
##   doi:10.5281/zenodo.1313986
## 
## A BibTeX entry for LaTeX users is
## 
##   @Manual{,
##     title = {{graphsim}: Simulate Expression data from igraph networks},
##     author = {S. Thomas Kelly and Michael A. Black},
##     year = {2020},
##     note = {R package version 1.0.0},
##     url = {https://github.com/TomKellyGenetics/graphsim},
##     doi = {10.5281/zenodo.1313986},
##   }
## 
## Please also acknowledge the manuscript describing use of
## this package once it is published.
## 
## Kelly, S.T. and Black, M.A. (2020) graphsim: An R package
## for simulating gene expression data from graph structures of
## biological pathways.
## 
## @Article{, title = {{graphsim}: An {R} package for
## simulating gene expression data from graph structures of
## biological pathways}, journal = {}, author = {S. Thomas
## Kelly and Michael A. Black}, year = {2020}, volume = {},
## number = {}, pages = {}, month = {}, note = {Submitted for
## peer-review}, url =
## {https://github.com/TomKellyGenetics/graphsim}, doi = {}, }
```

#### 5.2 Reporting issues

Please see the GitHub repository for reporting problems and requesting changes. Details on how to contribute can be found in the DESCRIPTION and README.

---

### 6 Session info

Here is the output of `sessionInfo()` on the system on which this document was compiled running pandoc 2.1:

```
## R version 4.0.2 (2020-06-22)
## Platform: x86_64-apple-darwin17.0 (64-bit)
## Running under: macOS Mojave 10.14.6
## 
## Matrix products: default
## BLAS:   /Library/Frameworks/R.framework/Versions/4.0/Resources/lib/libRblas.dylib
## LAPACK: /Library/Frameworks/R.framework/Versions/4.0/Resources/lib/libRlapack.dylib
## 
## locale:
## [1] en_GB.UTF-8/en_GB.UTF-8/en_GB.UTF-8/C/en_GB.UTF-8/en_GB.UTF-8
## 
## attached base packages:
## [1] stats     graphics  grDevices utils     datasets  methods  
## [7] base     
## 
## other attached packages:
## [1] scales_1.1.1   graphsim_1.0.0 gplots_3.0.3   igraph_1.2.5  
## 
## loaded via a namespace (and not attached):
##  [1] knitr_1.29         magrittr_1.5       munsell_0.5.0     
##  [4] colorspace_1.4-1   lattice_0.20-41    R6_2.4.1          
##  [7] rlang_0.4.6        stringr_1.4.0      caTools_1.18.0    
## [10] tools_4.0.2        matrixcalc_1.0-3   grid_4.0.2        
## [13] xfun_0.15          KernSmooth_2.23-17 htmltools_0.5.0   
## [16] gtools_3.8.2       yaml_2.2.1         digest_0.6.25     
## [19] lifecycle_0.2.0    Matrix_1.2-18      farver_2.0.3      
## [22] prettydoc_0.3.1    bitops_1.0-6       evaluate_0.14     
## [25] rmarkdown_2.3      gdata_2.18.0       stringi_1.4.6     
## [28] compiler_4.0.2     mvtnorm_1.1-1      pkgconfig_2.0.3
```

---
