## Supplementary material for "graphsim: An R package for simulating gene expression data from graph structures of biological pathways": graphsim 1.0.0 R package: distance.R

```
##' @rdname make_distance
##' @aliases make_relationship
##'
##' @title Generate Distance Matrix
##'
##' @description Compute the distance matrix of using shortest paths of a (directed)
##' \code{\link[igraph]{igraph}} structure, normalising by the diameter of the network,
##' preserving node/column/row names (and direction).
##'
##' @param mat precomputed adjacency or commonlink matrix.
##' @param graph An \code{\link[igraph]{igraph}} object. May be directed or weighted.
##' @param directed logical. Whether directed information is passed to the distance matrix.
##' @param absolute logical. Whether distances are scaled as the absolute difference
##' from the diameter (maximum possible). Defaults to TRUE. The alternative is to
##' calculate a relative difference from the diameter for a geometric decay in distance.
##' @keywords graph network igraph adjacency
##' @importFrom igraph as_adjacency_matrix
##' @import igraph
##' @examples 
##' 
##' # construct a synthetic graph module
##' library("igraph")
##' graph_test_edges <- rbind(c("A", "B"), c("B", "C"), c("B", "D"))
##' graph_test <- graph.edgelist(graph_test_edges, directed = TRUE)
##' 
##' # compute adjacency matrix for toy example
##' adjacency_matrix <- make_adjmatrix_graph(graph_test)
##' # compute nodes with relationships between nodes (geometrically decreasing by default)
##' distance_matrix_geom <- make_distance_adjmat(adjacency_matrix)
##' distance_matrix_geom
##' 
##' # compute nodes with relationships between nodes (arithmetically decreasing)
##' distance_matrix_abs <- make_distance_adjmat(adjacency_matrix, absolute = TRUE)
##' distance_matrix_abs
##' 
##' # compute Laplacian matrix
##' laplacian_matrix <- make_laplacian_graph(graph_test)
##' # compute distances from Laplacian
##' distance_matrix <- make_distance_laplacian(laplacian_matrix)
##' 
##' # construct a synthetic graph network
##' graph_structure_edges <- rbind(c("A", "C"), c("B", "C"), c("C", "D"), c("D", "E"),
##'                                c("D", "F"), c("F", "G"), c("F", "I"), c("H", "I"))
##' graph_structure <- graph.edgelist(graph_structure_edges, directed = TRUE)
##' # compute adjacency matrix for toy network
##' graph_structure_adjacency_matrix <- make_adjmatrix_graph(graph_structure)
##' # compute nodes with relationships between nodes (geometrically decreasing by default)
##' graph_structure_distance_matrix_geom <- make_distance_adjmat(graph_structure_adjacency_matrix)
##' graph_structure_distance_matrix_geom
##' # visualise matrix
##' library("gplots")
##' heatmap.2(graph_structure_distance_matrix_geom, scale = "none", trace = "none",
##'           col = colorpanel(50, "white", "red"))
##' # compute nodes with relationships between nodes (arithmetically decreasing)
##' graph_structure_distance_matrix_abs <- make_distance_adjmat(graph_structure_adjacency_matrix,
##'                                                             absolute = TRUE)
##' graph_structure_distance_matrix_abs
##' # visualise matrix
##' library("gplots")
##' heatmap.2(graph_structure_distance_matrix_abs,
##'           scale = "none", trace = "none",
##'           col = colorpanel(50, "white", "red"))
##'           
##' # import graph from package for reactome pathway
##' # TGF-\eqn{\Beta} receptor signaling activates SMADs (R-HSA-2173789)
##' TGFBeta_Smad_graph <- identity(TGFBeta_Smad_graph)
##' # compute nodes with relationships between nodes (geometrically decreasing by default)
##' TGFBeta_Smad_adjacency_matrix <- make_adjmatrix_graph(TGFBeta_Smad_graph)
##' TGFBeta_Smad_distance_matrix_geom <- make_distance_adjmat(TGFBeta_Smad_adjacency_matrix)
##' # visualise matrix
##' library("gplots")
##' heatmap.2(TGFBeta_Smad_distance_matrix_geom, scale = "none", trace = "none",
##'           col = colorpanel(50, "white", "red"))
##' # compute nodes with relationships between nodes (arithmetically decreasing)
##' TGFBeta_Smad_distance_matrix_abs <- make_distance_adjmat(TGFBeta_Smad_adjacency_matrix,
##'                         absolute = TRUE)
##' # visualise matrix
##' library("gplots")
##' heatmap.2(TGFBeta_Smad_distance_matrix_abs, scale = "none", trace = "none",
##'           col = colorpanel(50, "white", "red"))
##' 
##' @return A numeric matrix of values in the range [0, 1] where higher values are closer in the network
##' 
##' @export
make_distance_graph <- function(graph, directed = FALSE, absolute = FALSE){
  if(directed == FALSE) graph <- as.undirected(graph)
  diam <- diameter(graph)
  if (absolute){
    paths <- shortest.paths(graph)
    diam <- max(diam, max(paths))
    mat <- (diam-paths)/diam
  } else {
    paths <- shortest.paths(graph)
    diam <- max(diam, max(paths))
    mat <- 1^-diam/(diam*paths)
    diag(mat) <- 1
  }
  rownames(mat) <- colnames(mat) <- names(V(graph))
  return(mat)
}

##' @rdname make_distance
##' @importFrom igraph graph_from_adjacency_matrix
##' @export
make_distance_adjmat <- function(mat, directed = FALSE, absolute = FALSE){
  diag(mat) <- 0
  graph <- graph_from_adjacency_matrix(mat, weighted = NULL, mode = ifelse(directed, "directed", "undirected"))
  diam <- diameter(graph)
  if (absolute){
    paths <- shortest.paths(graph)
    diam <- max(diam, max(paths))
    mat <- (diam-paths)/diam
  } else {
    paths <- shortest.paths(graph)
    diam <- max(diam, max(paths))
    mat <- 1^-diam/(diam*paths)
    diag(mat) <- 1
  }
  rownames(mat) <- colnames(mat) <- names(V(graph))
  return(mat)
}

##' @rdname make_distance
##' @importFrom igraph graph_from_adjacency_matrix laplacian_matrix
##' @export
make_distance_laplacian <- function(mat, directed = FALSE, absolute = FALSE){
  diag(mat) <- 0
  adj_mat <- ifelse(mat < 0, abs(mat), 0)
  graph <- graph_from_adjacency_matrix(adj_mat, weighted = TRUE, mode = ifelse(directed, "directed", "undirected"))
  diam <- diameter(graph)
  if (absolute){
    paths <- shortest.paths(graph)
    diam <- max(diam, max(paths))
    mat <- (diam-paths)/diam
  } else {
    paths <- shortest.paths(graph)
    diam <- max(diam, max(paths))
    mat <- 1^-diam/(diam*paths)
    diag(mat) <- 1
  }
  rownames(mat) <- colnames(mat) <- names(V(graph))
  return(mat)
}
```
