## Supplementary material for "graphsim: An R package for simulating gene expression data from graph structures of biological pathways": graphsim 1.0.0 R package: test_distance.R

```
library("graphsim")
library("igraph")
context("Make Distance Matrix")

test_that("Generate distance relation matrix from adjacency matrix", {
  graph_test1_edges <- rbind(c("A", "B"), c("B", "C"), c("B", "D"))
  graph_test1 <- graph.edgelist(graph_test1_edges, directed = TRUE)
  adjacency_matrix1 <- make_adjmatrix_graph(graph_test1)
  distance_matrix1 <- make_distance_adjmat(adjacency_matrix1)
  expect_equal(isSymmetric(distance_matrix1), TRUE)
  expect_true(all(diag(distance_matrix1) == rep(1, nrow(adjacency_matrix1))))
  expect_equal(nrow(distance_matrix1), length(V(graph_test1)))
  expect_equal(ncol(distance_matrix1), length(V(graph_test1)))
  expect_equal(sum(distance_matrix1), 8.5)
  expect_true(all(adjacency_matrix1 == cbind(c(0, 1, 0, 0), c(1, 0, 1, 1), c(0, 1, 0, 0), c(0, 1, 0, 0))))
  expect_true(all(distance_matrix1 == 1/cbind(c(1, 2, 4, 4), c(2, 1, 2, 2), c(4, 2, 1, 4), c(4, 2, 4, 1))))
  expect_true(all(distance_matrix1 %in% c(0, 1/2^c(0:100))))
})

test_that("Generate distance relation matrix from commonlink matrix", {
  graph_test1_edges <- rbind(c("A", "B"), c("B", "C"), c("B", "D"))
  graph_test1 <- graph.edgelist(graph_test1_edges, directed = TRUE)
  adjacency_matrix1 <- make_adjmatrix_graph(graph_test1)
  common_link_matrix1 <- make_commonlink_adjmat(adjacency_matrix1)
  distance_matrix1 <- make_distance_adjmat(common_link_matrix1)
  expect_equal(isSymmetric(distance_matrix1), TRUE)
  expect_true(all(diag(distance_matrix1) == rep(1, nrow(adjacency_matrix1))))
  expect_equal(nrow(distance_matrix1), length(V(graph_test1)))
  expect_equal(ncol(distance_matrix1), length(V(graph_test1)))
  expect_equal(sum(distance_matrix1), 10)
  expect_true(all(adjacency_matrix1 == cbind(c(0, 1, 0, 0), c(1, 0, 1, 1), c(0, 1, 0, 0), c(0, 1, 0, 0))))
  expect_true(all(common_link_matrix1 == cbind(c(1, 0, 1, 1), c(0, 3, 0, 0), c(1, 0, 1, 1), c(1, 0, 1, 1))))
  expect_true(all(distance_matrix1 == cbind(c(1, 0, 1, 1), c(0, 1, 0, 0), c(1, 0, 1, 1), c(1, 0, 1, 1))))
  expect_true(all(distance_matrix1 %in% c(0, 1/2^c(0:100))))
})


test_that("Generate distance relation matrix matrix from graph structure", {
  graph_test1_edges <- rbind(c("A", "B"), c("B", "C"), c("B", "D"))
  graph_test1 <- graph.edgelist(graph_test1_edges, directed = TRUE)
  adjacency_matrix1 <- make_adjmatrix_graph(graph_test1)
  common_link_matrix1 <- make_commonlink_graph(graph_test1)
  distance_matrix1 <- make_distance_graph(graph_test1)
  expect_equal(isSymmetric(distance_matrix1), TRUE)
  expect_true(all(diag(distance_matrix1) == rep(1, nrow(adjacency_matrix1))))
  expect_equal(nrow(distance_matrix1), length(V(graph_test1)))
  expect_equal(ncol(distance_matrix1), length(V(graph_test1)))
  expect_equal(sum(distance_matrix1), 8.5)
  expect_true(all(adjacency_matrix1 == cbind(c(0, 1, 0, 0), c(1, 0, 1, 1), c(0, 1, 0, 0), c(0, 1, 0, 0))))
  expect_true(all(distance_matrix1 == 1/cbind(c(1, 2, 4, 4), c(2, 1, 2, 2), c(4, 2, 1, 4), c(4, 2, 4, 1))))
  expect_true(all(distance_matrix1 %in% c(0, 1/2^c(0:100))))
})
```
