## Supplementary material for "graphsim: An R package for simulating gene expression data from graph structures of biological pathways": graphsim 1.0.0 R package: test_laplacian.R

```
library("graphsim")
library("igraph")
context("Make Commonlink Matrix")

test_that("Generate Laplacian matrix from adjacency matrix", {
  graph_test1_edges <- rbind(c("A", "B"), c("B", "C"), c("B", "D"))
  graph_test1 <- graph.edgelist(graph_test1_edges, directed = TRUE)
  adjacency_matrix1 <- make_adjmatrix_graph(graph_test1)
  laplacian_matrix1 <- make_laplacian_adjmat(adjacency_matrix1)
  expect_equal(isSymmetric(laplacian_matrix1), TRUE)
  expect_equal(diag(laplacian_matrix1), degree(graph_test1))
  expect_equal(sum(diag(laplacian_matrix1)), sum(degree(graph_test1)))
  expect_equal(nrow(laplacian_matrix1), length(V(graph_test1)))
  expect_equal(ncol(laplacian_matrix1), length(V(graph_test1)))
  expect_equal(sum(laplacian_matrix1), 0)
  expect_true(all(adjacency_matrix1 == cbind(c(0, 1, 0, 0), c(1, 0, 1, 1), c(0, 1, 0, 0), c(0, 1, 0, 0))))
  expect_equal(all(is.matrix(laplacian_matrix1)), TRUE)
  expect_true(all(laplacian_matrix1 == cbind(c(1, -1, 0, 0), c(-1, 3, -1, -1), c(0, -1, 1, 0), c(0,-1, 0, 1))))
})

test_that("Generate Laplacian matrix from graph structure", {
  graph_test1_edges <- rbind(c("A", "B"), c("B", "C"), c("B", "D"))
  graph_test1 <- graph.edgelist(graph_test1_edges, directed = TRUE)
  laplacian_matrix1 <- make_laplacian_graph(graph_test1)
  expect_equal(isSymmetric(laplacian_matrix1), TRUE)
  expect_equal(diag(laplacian_matrix1), degree(graph_test1))
  expect_equal(sum(diag(laplacian_matrix1)), sum(degree(graph_test1)))
  expect_equal(nrow(laplacian_matrix1), length(V(graph_test1)))
  expect_equal(ncol(laplacian_matrix1), length(V(graph_test1)))
  expect_equal(sum(laplacian_matrix1), 0)
  expect_equal(all(is.matrix(laplacian_matrix1)), TRUE)
  expect_true(all(laplacian_matrix1 == cbind(c(1, -1, 0, 0), c(-1, 3, -1, -1), c(0, -1, 1, 0), c(0,-1, 0, 1))))
})
```
