## Supplementary material for "graphsim: An R package for simulating gene expression data from graph structures of biological pathways": graphsim 1.0.0 R package: plots_directed.R

```
## ---- include = FALSE---------------------------------------------
knitr::opts_chunk$set(collapse = TRUE, comment = "#>", width = 68)
knitr::opts_chunk$set(fig.cap = "", fig.path = "Plot")
knitr::opts_chunk$set(fig.align = "center")
options(width = 68, cli.unicode = FALSE, cli.width = 68)
#par(mai=c(2.82, 2.82, 0.82, 0.82)-0.82)
par(mar=c(7, 10, 4, 2) + 0.1)
par(bty="o")
#captioner::captioner(prefix = "Fig.")

## ---- message=F---------------------------------------------------
library("igraph")
library("scales")
library("graphsim")

## -----------------------------------------------------------------
graph_edges <- rbind(c("A", "C"), c("B", "C"), c("C", "D"), c("D", "E"),
                     c("D", "F"), c("F", "G"), c("F", "I"), c("H", "I"))
graph <- graph.edgelist(graph_edges, directed = TRUE)

## ----  fig.align='center', out.width="80%",fig.height = 6, fig.width = 6, fig.retina=1.5----
plot_directed(graph)

## ----  fig.align='center', out.width="80%",fig.height = 6, fig.width = 6, fig.retina=1.5----
plot_directed(graph, layout = layout.kamada.kawai)

## ----  fig.align='center', out.width="80%",fig.height = 6, fig.width = 6, fig.retina=1.5----
plot_directed(graph, fill.node = "lightblue", border.node = "royalblue")

## ----  fig.align='center', out.width="80%",fig.height = 6, fig.width = 6, fig.retina=1.5----
names(V(graph))
colour_vector <- ifelse(names(V(graph)) %in% c("A", "D", "I"), 1, 2)
plot_directed(graph, fill.node = c("lightblue", "grey")[colour_vector], border.node = c("royalblue", "black")[colour_vector])

## ----  fig.align='center', out.width="80%",fig.height = 6, fig.width = 6, fig.retina=1.5----
plot_directed(graph, state = "activating")

## ----  fig.align='center', out.width="80%", fig.height = 6, fig.width = 6, fig.retina=6, eval=FALSE----
#  plot_directed(graph, state = 1)
#  plot_directed(graph, state = 0)

## ----  fig.align='center', out.width="80%",fig.height = 6, fig.width = 6, fig.retina=1.5----
plot_directed(graph, state = "inhibiting")

## ----  fig.align='center', out.width="80%", fig.height = 6, fig.width = 6, fig.retina=6, eval=FALSE----
#  plot_directed(graph, state = -1)
#  plot_directed(graph, state = 2)

## ----  fig.align='center', out.width="80%",fig.height = 6, fig.width = 6, fig.retina=1.5----
E(graph)
plot_directed(graph, state = c(1, 1, -1, -1, 1, -1, 1, -1))

## ----  fig.align='center', out.width="80%",fig.height = 6, fig.width = 6, fig.retina=1.5----
edge_properties <- c(1, 1, -1, -1, 1, -1, 1, -1)/2 + 1.5
plot_directed(graph, state = edge_properties, col.arrow = c("darkgreen", "red")[edge_properties])

## ----  fig.align='center', out.width="80%",fig.height = 6, fig.width = 6, fig.retina=1.5----
edge_properties <- c(1, 1, -1, -1, 1, -1, 1, -1)/2 + 1.5
ggplot_colours <- c("#F8766D", "#CD9600", "#7CAE00", "#00BE67",
                    "#00BFC4", "#00A9FF", "#C77CFF", "#FF61CC")
plot_directed(graph, state = edge_properties,
              col.arrow = ggplot_colours, fill.node =  ggplot_colours)

## ----  fig.align='center', out.width="80%", fig.height = 5, fig.width = 5, warning=FALSE, message = FALSE----
graph <- identity(RAF_MAP_graph)
plot_directed(graph,col.arrow = alpha("#00A9FF", 0.25),
              fill.node = "lightblue",
              layout = layout.kamada.kawai)

## ----  fig.align='center', out.width="80%",fig.height = 6, fig.width = 6, fig.retina=1.5----
graph <- identity(Pi3K_graph)
plot_directed(graph, col.arrow = alpha("#00A9FF", 0.25),
              fill.node = "lightblue",
              layout = layout.kamada.kawai)

## ----  fig.align='center', out.width="80%",fig.height = 6, fig.width = 6, fig.retina=1.5----
graph <- identity(TGFBeta_Smad_graph)
edge_properties <- E(graph)$state
plot_directed(graph, state = edge_properties,
              col.arrow = c(alpha("navyblue", 0.25), alpha("red", 0.25))[edge_properties],
              fill.node = c("lightblue"),
              layout = layout.kamada.kawai)

## ---- echo = FALSE------------------------------------------------
sessionInfo()
```
