## Supplementary material for "graphsim: An R package for simulating gene expression data from graph structures of biological pathways": graphsim 1.0.0 R package: simulate_expression.html

```
## R version 4.0.2 (2020-06-22)
## Platform: x86_64-apple-darwin17.0 (64-bit)
## Running under: macOS Mojave 10.14.6
## 
## Matrix products: default
## BLAS:   /Library/Frameworks/R.framework/Versions/4.0/Resources/lib/libRblas.dylib
## LAPACK: /Library/Frameworks/R.framework/Versions/4.0/Resources/lib/libRlapack.dylib
## 
## locale:
## [1] en_GB.UTF-8/en_GB.UTF-8/en_GB.UTF-8/C/en_GB.UTF-8/en_GB.UTF-8
## 
## attached base packages:
## [1] stats     graphics  grDevices utils     datasets  methods  
## [7] base     
## 
## other attached packages:
## [1] scales_1.1.1   graphsim_1.0.0 gplots_3.0.3   igraph_1.2.5  
## 
## loaded via a namespace (and not attached):
##  [1] knitr_1.29         magrittr_1.5       munsell_0.5.0     
##  [4] colorspace_1.4-1   lattice_0.20-41    R6_2.4.1          
##  [7] rlang_0.4.6        stringr_1.4.0      caTools_1.18.0    
## [10] tools_4.0.2        matrixcalc_1.0-3   grid_4.0.2        
## [13] xfun_0.15          KernSmooth_2.23-17 htmltools_0.5.0   
## [16] gtools_3.8.2       yaml_2.2.1         digest_0.6.25     
## [19] lifecycle_0.2.0    Matrix_1.2-18      farver_2.0.3      
## [22] codetools_0.2-16   prettydoc_0.3.1    bitops_1.0-6      
## [25] evaluate_0.14      rmarkdown_2.3      gdata_2.18.0      
## [28] stringi_1.4.6      compiler_4.0.2     mvtnorm_1.1-1     
## [31] pkgconfig_2.0.3
```

---
