## Supplementary material for "graphsim: An R package for simulating gene expression data from graph structures of biological pathways": graphsim 1.0.0 R package: simulate_expression.R

```
## ---- include = FALSE---------------------------------------------
#knitr::opts_chunk$set(collapse = TRUE, comment = "#>", width = 68)
knitr::opts_chunk$set(fig.cap = "", fig.path = "Plot")
options(width = 68, cli.unicode = FALSE, cli.width = 68)
#par(mai=c(2.82, 2.82, 0.82, 0.82)-0.82)
par(mar=c(7, 10, 4, 2) + 0.1)
par(bty="o")
#captioner::captioner(prefix = "Fig.")

## ---- eval = FALSE------------------------------------------------
#  #install packages required (once per computer)
#  install.packages("graphsim")

## ---- eval = FALSE------------------------------------------------
#  #install stable release
#  remotes::install_github("TomKellyGenetics", ref = "master")
#  #install development version
#  remotes::install_github("TomKellyGenetics", ref = "dev")

## ---- warning=FALSE, results='hide', message=FALSE----------------
library("igraph")
library("gplots")
library("graphsim")
library("scales")

## ---- out.width = '50%', out.height  = '50%', fig.align='center', fig.height = 6, fig.width = 6, fig.retina=1.5----
graph_structure_edges <- rbind(c("A", "C"), c("B", "C"), c("C", "D"),c("D", "E"),
                               c("D", "F"), c("F", "G"), c("F", "I"), c("H", "I"))
graph_structure <- graph.edgelist(graph_structure_edges, directed = T)
plot_directed(graph_structure, layout = layout.kamada.kawai)

## ---- out.width = '50%', out.height  = '50%', fig.align='center', dpi=36, fig.retina=1,message=FALSE, warning=FALSE----
expr <- generate_expression(100, graph_structure, cor = 0.8, mean = 0,
                            comm = FALSE, dist = TRUE, absolute = FALSE)
heatmap.2(expr, scale = "none", trace = "none",
          col = bluered(50), colsep = 1:4, rowsep = 1:4)

## ---- out.width = '50%', out.height  = '50%', fig.align='center', dpi=54----
adj_mat <- make_adjmatrix_graph(graph_structure)
heatmap.2(make_adjmatrix_graph(graph_structure),
          scale = "none", trace = "none",
          col = colorpanel(3, "grey75", "white", "blue"),
          colsep = 1:4, rowsep = 1:4)

## ---- out.width = '50%', out.height  = '50%', fig.align='center', dpi=54----
heatmap.2(make_adjmatrix_graph(graph_structure, directed = T),
          scale = "none", trace = "none",
          col = colorpanel(3, "grey75", "white", "blue"),
          colsep = 1:4, rowsep = 1:4)

## ---- out.width = '50%', out.height  = '50%', fig.align='center', dpi=54----
comm_mat <- make_commonlink_graph(graph_structure)
heatmap.2(make_commonlink_graph(graph_structure),
          scale = "none", trace = "none",
          col = colorpanel(50, "grey75", "red"),
          colsep = 1:4, rowsep = 1:4)

## ---- out.width = '50%', out.height  = '50%', fig.align='center', dpi=54----
laplacian_mat <- make_laplacian_graph(graph_structure)
heatmap.2(make_laplacian_graph(graph_structure),
          scale = "none", trace = "none", 
          col = bluered(50),colsep = 1:4, rowsep = 1:4)

## ---- out.width = '50%', out.height  = '50%', fig.align='center', dpi=54----
shortest.paths(graph_structure)
heatmap.2(shortest.paths(graph_structure),
          scale = "none", trace = "none",
          col = colorpanel(50, "grey75", "red"),
           thincolsep = 1:4, rowsep = 1:4)

## ---- out.width = '50%', out.height  = '50%', fig.align='center', dpi=54----
diam <- diameter(graph_structure)
relative_dist <- (1+diam-shortest.paths(graph_structure))/diam
relative_dist
heatmap.2(relative_dist, scale = "none", trace = "none",
          col = colorpanel(50, "grey75", "red"),
          colsep = 1:4, rowsep = 1:4)

## ---- out.width = '50%', out.height  = '50%', fig.align='center', dpi=54----
dist_mat <- make_distance_graph(graph_structure, absolute = F)
dist_mat
heatmap.2(dist_mat, scale = "none", trace = "none",
          col = colorpanel(50, "grey75", "red"),
          colsep = 1:4, rowsep = 1:4)

## ---- out.width = '50%', out.height  = '50%', fig.align='center', dpi=54----
dist_mat <- make_distance_graph(graph_structure, absolute = T)
dist_mat
heatmap.2(dist_mat, scale = "none", trace = "none",
          col = colorpanel(50, "grey75", "red"),
          colsep = 1:4, rowsep = 1:4)

## ---- out.width = '50%', out.height  = '50%', fig.align='center', dpi=54----
#sigma from adj mat
sigma_mat <- make_sigma_mat_graph(graph_structure, 0.8)
sigma_mat
heatmap.2(sigma_mat, scale = "none", trace = "none",
          col = colorpanel(50, "grey75", "red"),
          colsep = 1:4, rowsep = 1:4)

## ---- out.width = '50%', out.height  = '50%', fig.align='center', dpi=54----
#sigma from comm mat
sigma_mat <- make_sigma_mat_graph(graph_structure, 0.8, comm = T)
sigma_mat
heatmap.2(sigma_mat, scale = "none", trace = "none",
          col = colorpanel(50, "grey75", "red"),
          colsep = 1:4, rowsep = 1:4)

## ---- out.width = '50%', out.height  = '50%', fig.align='center', dpi=54----
# sigma from geometric distance matrix
make_sigma_mat_dist_graph(graph_structure, 0.8, absolute = F)
heatmap.2(make_sigma_mat_dist_graph(graph_structure, 0.8, absolute = F),
          scale = "none", trace = "none",
          col = colorpanel(50, "grey75", "red"),
          colsep = 1:4, rowsep = 1:4)

## ---- out.width = '50%', out.height  = '50%', fig.align='center', dpi=54----
# sigma from absolute distance matrix
sigma_mat <- make_sigma_mat_dist_graph(graph_structure, 0.8, absolute = T)
heatmap.2(sigma_mat, scale = "none", trace = "none", 
          col = colorpanel(50, "grey75", "red"),
          colsep = 1:4, rowsep = 1:4)

## ---- out.width = '50%', out.height  = '50%', fig.align='center', dpi=54, warning=FALSE----
#simulate expression data
#adj mat
expr <- generate_expression(100, graph_structure, cor = 0.8,
                            mean = 0, comm = F)
heatmap.2(expr, scale = "none", trace = "none",
          col = bluered(50), colsep = 1:4, rowsep = 1:4)
heatmap.2(cor(t(expr)), scale = "none", trace = "none",
          col = colorpanel(50, "grey75", "red"),
          colsep = 1:4, rowsep = 1:4)

## ---- out.width = '50%', out.height  = '50%', fig.align='center', dpi=54----
#comm mat
expr <- generate_expression(100, graph_structure, cor = 0.8, mean = 0, comm =T)
heatmap.2(expr, scale = "none", trace = "none",
          col = bluered(50), colsep = 1:4, rowsep = 1:4)
heatmap.2(cor(t(expr)), scale = "none", trace = "none",
          col = colorpanel(50, "grey75", "red"),
          colsep = 1:4, rowsep = 1:4)

## ---- out.width = '50%', out.height  = '50%', fig.align='center', dpi=54----
# relative dist
expr <- generate_expression(100, graph_structure, cor = 0.8,mean = 0,
                            comm = F, dist = T, absolute = F)
heatmap.2(expr, scale = "none", trace = "none", col = bluered(50),
          colsep = 1:4, rowsep = 1:4)
heatmap.2(cor(t(expr)), scale = "none", trace = "none",
          col = colorpanel(50, "grey75", "red"),
          colsep = 1:4, rowsep = 1:4)

## ---- include = FALSE---------------------------------------------
set.seed(9000)

## ---- out.width = '50%', out.height  = '50%', fig.align='center', dpi=54, warning=FALSE----
#absolute dist
expr <- generate_expression(100, graph_structure, cor = 0.8, mean = 0,
                            comm = F, dist = T, absolute = T) 
heatmap.2(expr, scale = "none", trace = "none", col = bluered(50),
          colsep = 1:4, rowsep = 1:4)
heatmap.2(cor(t(expr)), scale = "none", trace = "none",
           col = bluered(50), colsep = 1:4, rowsep = 1:4)

## ---- include = FALSE---------------------------------------------
set.seed(9000)

## ----simulation_activating_hide, fig.show='hold', out.width = '50%', out.height  = '50%', fig.align='center', dpi=36, fig.retina=1, fig.margin = FALSE, fig.ncol = 2, warning=FALSE, message=FALSE, cache=TRUE----
# activating graph
state <- rep(1, length(E(graph_structure)))
plot_directed(graph_structure, state=state,
              layout = layout.kamada.kawai,
              cex.node=2, cex.arrow=4, arrow_clip = 0.2)
mtext(text = "(a) Activating pathway structure", side=1,
      line=3.5, at=0.075, adj=0.5, cex=1.75)
box()
#plot relationship matrix
heatmap.2(make_distance_graph(graph_structure, absolute = FALSE),
          scale = "none", trace = "none",
          col = colorpanel(50, "white", "red"),
          colsep = 1:length(V(graph_structure)),
          rowsep = 1:length(V(graph_structure)))
mtext(text = "(b) Relationship matrix", side=1,
      line=3.5, at=0, adj=0.5, cex=1.75)
#plot sigma matrix
sigma_mat <- make_sigma_mat_dist_graph(graph_structure,
               cor = 0.8, absolute = FALSE)
heatmap.2(sigma_mat, scale = "none", trace = "none",
          col = colorpanel(50, "white", "red"),
          colsep = 1:length(V(graph_structure)),
          rowsep = 1:length(V(graph_structure)))
mtext(text = expression(paste("(c) ", Sigma, " matrix")), side=1,
      line=3.5, at=0, adj=0.5, cex=1.75)
#simulated data
expr <- generate_expression(100, graph_structure, cor = 0.8, mean = 0,
          comm = FALSE, dist =TRUE, absolute = FALSE, state = state)
#plot simulated correlations
heatmap.2(cor(t(expr)), scale = "none", trace = "none",
          col = colorpanel(50, "white", "red"),
          colsep = 1:length(V(graph_structure)),
          rowsep = 1:length(V(graph_structure)))
mtext(text = "(d) Simulated correlation", side=1,
      line=3.5, at=0, adj=0.5, cex=1.75)
#plot simulated expression data
heatmap.2(expr, scale = "none", trace = "none", col = bluered(50),
          colsep = 1:length(V(graph_structure)),
          rowsep = 1:length(V(graph_structure)), labCol = "")
mtext(text = "samples", side=1, line=1.5, at=0.2, adj=0.5, cex=1.5)
mtext(text = "genes", side=4, line=1, at=-0.4, adj=0.5, cex=1.5)
mtext(text = "(e) Simulated expression data (log scale)", side=1,
      line=3.5, at=0, adj=0.5, cex=1.75)

## ---- include = FALSE---------------------------------------------
set.seed(9000)

## ----simulation_inhibiting_hide, fig.show='hold', out.width = '50%', out.height  = '50%', fig.align='center', dpi=36, fig.retina=1, fig.margin = FALSE, fig.ncol = 2, warning=FALSE, message=FALSE, cache=TRUE----
# activating graph
state <- state <- c(1, 1, -1, 1, 1, 1, 1, -1)
plot_directed(graph_structure, state=state, layout = layout.kamada.kawai,
              cex.node=2, cex.arrow=4, arrow_clip = 0.2)
mtext(text = "(a) Activating pathway structure", side=1,
      line=3.5, at=0.075, adj=0.5, cex=1.75)
box()
#plot relationship matrix
heatmap.2(make_distance_graph(graph_structure, absolute = FALSE),
          scale = "none", trace = "none",
          col = colorpanel(50, "white", "red"),
          colsep = 1:length(V(graph_structure)),
          rowsep = 1:length(V(graph_structure)))
mtext(text = "(b) Relationship matrix", side=1,
      line=3.5, at=0, adj=0.5, cex=1.75)
#plot sigma matrix
heatmap.2(make_sigma_mat_dist_graph(graph_structure, state,
                                    cor = 0.8, absolute = FALSE),
          scale = "none", trace = "none",
          col = colorpanel(50, "blue", "white", "red"),
          colsep = 1:length(V(graph_structure)),
          rowsep = 1:length(V(graph_structure)))
mtext(text = expression(paste("(c) ", Sigma, " matrix")), side=1,
      line=3.5, at=0, adj=0.5, cex=1.75)
#simulated data
expr <- generate_expression(100, graph_structure, state,
                            cor = 0.8, mean = 0,
                            comm = FALSE, dist =TRUE,
                            absolute = FALSE, state = state)
#plot simulated correlations
heatmap.2(cor(t(expr)), scale = "none", trace = "none",
          col = colorpanel(50, "blue", "white", "red"),
          colsep = 1:length(V(graph_structure)),
          rowsep = 1:length(V(graph_structure)))
mtext(text = "(d) Simulated correlation", side=1,
      line=3.5, at=0, adj=0.5, cex=1.75)
#plot simulated expression data
heatmap.2(expr, scale = "none", trace = "none", col = bluered(50),
          colsep = 1:length(V(graph_structure)),
          rowsep = 1:length(V(graph_structure)), labCol = "")
mtext(text = "samples", side=1, line=1.5, at=0.2, adj=0.5, cex=1.5)
mtext(text = "genes", side=4, line=1, at=-0.4, adj=0.5, cex=1.5)
mtext(text = "(e) Simulated expression data (log scale)", side=1,
      line=3.5, at=0, adj=0.5, cex=1.75)


## ---- include = FALSE---------------------------------------------
set.seed(9000)

## ----simulation_graph_diverging_hide, fig.show='hold', out.width = '50%', out.height  = '50%', fig.align='center', dpi=36, fig.retina=1, fig.margin = FALSE, fig.ncol = 2, warning=FALSE, message=FALSE, cache=TRUE----
graph_diverging_edges <- rbind(c("A", "B"), c("B", "C"), c("B", "D"))
graph_diverging <- graph.edgelist(graph_diverging_edges, directed = T)

# activating graph
state <- rep(1, length(E(graph_diverging)))
plot_directed(graph_diverging, state=state,
              layout = layout.kamada.kawai,
              cex.node=2, cex.arrow=4, arrow_clip = 0.2)
mtext(text = "(a) Activating pathway structure", side=1,
      line=3.5, at=0.075, adj=0.5, cex=1.75)
box()
#plot relationship matrix
heatmap.2(make_distance_graph(graph_diverging, absolute = FALSE),
          scale = "none", trace = "none",
          col = colorpanel(50, "white", "red"),
          colsep = 1:length(V(graph_diverging)),
          rowsep = 1:length(V(graph_diverging)))
mtext(text = "(b) Relationship matrix", side=1,
      line=3.5, at=0, adj=0.5, cex=1.75)
#plot sigma matrix
heatmap.2(make_sigma_mat_dist_graph(graph_diverging,
                                    cor = 0.8, absolute = FALSE),
          scale = "none", trace = "none",
          col = colorpanel(50, "white", "red"),
          colsep = 1:length(V(graph_diverging)),
          rowsep = 1:length(V(graph_diverging)))
mtext(text = expression(paste("(c) ", Sigma, " matrix")), side=1,
      line=3.5, at=0, adj=0.5, cex=1.75)
#simulated data
expr <- generate_expression(100, graph_diverging,
                            cor = 0.8, mean = 0,
                            comm = FALSE, dist =TRUE,
                            absolute = FALSE, state = state)
#plot simulated correlations
heatmap.2(cor(t(expr)), scale = "none", trace = "none",
          col = colorpanel(50, "white", "red"),
          colsep = 1:length(V(graph_diverging)),
          rowsep = 1:length(V(graph_diverging)))
mtext(text = "(d) Simulated correlation", side=1,
      line=3.5, at=0, adj=0.5, cex=1.75)
#plot simulated expression data
heatmap.2(expr, scale = "none", trace = "none",col = bluered(50),
          colsep = 1:length(V(graph_diverging)),
          rowsep = 1:length(V(graph_diverging)), labCol = "")
mtext(text = "samples", side=1, line=1.5, at=0.2, adj=0.5, cex=1.5)
mtext(text = "genes", side=4, line=1, at=-0.4, adj=0.5, cex=1.5)
mtext(text = "(e) Simulated expression data (log scale)", side=1,
      line=3.5, at=0, adj=0.5, cex=1.75)


## ---- include = FALSE---------------------------------------------
set.seed(9000)

## ----simulation_graph_diverging_inhibiting_hide, fig.show='hold', out.width = '50%', out.height  = '50%', fig.align='center', dpi=36, fig.retina=1, fig.margin = FALSE, fig.ncol = 2, warning=FALSE, message=FALSE, cache=TRUE----
# activating graph
state <- state <- c(1, 1, -1)
plot_directed(graph_diverging, state=state, layout = layout.kamada.kawai,
              cex.node=2, cex.arrow=4, arrow_clip = 0.2)
mtext(text = "(a) Activating pathway structure", side=1,
      line=3.5, at=0.075, adj=0.5, cex=1.75)
box()
#plot relationship matrix
heatmap.2(make_distance_graph(graph_diverging, absolute = FALSE),
          scale = "none", trace = "none",
          col = colorpanel(50, "white", "red"),
          colsep = 1:length(V(graph_diverging)),
          rowsep = 1:length(V(graph_diverging)))
mtext(text = "(b) Relationship matrix", side=1,
      line=3.5, at=0, adj=0.5, cex=1.75)
#plot sigma matrix
heatmap.2(make_sigma_mat_dist_graph(graph_diverging, state,
                                    cor = 0.8, absolute = FALSE),
          scale = "none", trace = "none",
          col = colorpanel(50, "blue", "white", "red"),
          colsep = 1:length(V(graph_diverging)),
          rowsep = 1:length(V(graph_diverging)))
mtext(text = expression(paste("(c) ", Sigma, " matrix")), side=1,
      line=3.5, at=0, adj=0.5, cex=1.75)
#simulated data
expr <- generate_expression(100, graph_diverging, state,
                            cor = 0.8, mean = 0,
                            comm = FALSE, dist =TRUE,
                            absolute = FALSE, state = state)
#plot simulated correlations
heatmap.2(cor(t(expr)), scale = "none", trace = "none",
          col = colorpanel(50, "blue", "white", "red"),
          colsep = 1:length(V(graph_diverging)),
          rowsep = 1:length(V(graph_diverging)))
mtext(text = "(d) Simulated correlation", side=1,
      line=3.5, at=0, adj=0.5, cex=1.75)
#plot simulated expression data
heatmap.2(expr, scale = "none", trace = "none", col = bluered(50),
          colsep = 1:length(V(graph_diverging)),
          rowsep = 1:length(V(graph_diverging)), labCol = "")
mtext(text = "samples", side=1, line=1.5, at=0.2, adj=0.5, cex=1.5)
mtext(text = "genes", side=4, line=1, at=-0.4, adj=0.5, cex=1.5)
mtext(text = "(e) Simulated expression data (log scale)", side=1,
      line=3.5, at=0, adj=0.5, cex=1.75)


## ---- include = FALSE---------------------------------------------
set.seed(9000)

## ----simulation_graph_converging_hide, fig.show='hold', out.width = '50%', out.height  = '50%', fig.align='center', dpi=36, fig.retina=1, fig.margin = FALSE, fig.ncol = 2, warning=FALSE, message=FALSE, cache=TRUE----
graph_converging_edges <- rbind(c("C", "E"), c("D", "E"), c("E", "F"))
graph_converging <- graph.edgelist(graph_converging_edges, directed = T)

# activating graph
state <- rep(1, length(E(graph_converging)))
plot_directed(graph_converging, state=state,
              layout = layout.kamada.kawai,
              cex.node=2, cex.arrow=4, arrow_clip = 0.2)
mtext(text = "(a) Activating pathway structure", side=1,
      line=3.5, at=0.075, adj=0.5, cex=1.75)
box()
#plot relationship matrix
heatmap.2(make_distance_graph(graph_converging, absolute = FALSE),
          scale = "none", trace = "none",
          col = colorpanel(50, "white", "red"),
          colsep = 1:length(V(graph_converging)),
          rowsep = 1:length(V(graph_converging)))
mtext(text = "(b) Relationship matrix", side=1,
      line=3.5, at=0, adj=0.5, cex=1.75)
#plot sigma matrix
heatmap.2(make_sigma_mat_dist_graph(graph_converging, cor = 0.8,
                                    absolute = FALSE),
          scale = "none", trace = "none",
          col = colorpanel(50, "white", "red"),
          colsep = 1:length(V(graph_converging)),
          rowsep = 1:length(V(graph_converging)))
mtext(text = expression(paste("(c) ", Sigma, " matrix")), side=1,
      line=3.5, at=0, adj=0.5, cex=1.75)
#simulated data
expr <- generate_expression(100, graph_converging,
                            cor = 0.8, mean = 0,
                            comm = FALSE, dist =TRUE,
                            absolute = FALSE, state = state)
#plot simulated correlations
heatmap.2(cor(t(expr)), scale = "none", trace = "none",
          col = colorpanel(50, "white", "red"),
          colsep = 1:length(V(graph_converging)),
          rowsep = 1:length(V(graph_converging)))
mtext(text = "(d) Simulated correlation", side=1,
      line=3.5, at=0, adj=0.5, cex=1.75)
#plot simulated expression data
heatmap.2(expr, scale = "none", trace = "none", col = bluered(50),
          colsep = 1:length(V(graph_converging)),
          rowsep = 1:length(V(graph_converging)), labCol = "")
mtext(text = "samples", side=1, line=1.5, at=0.2, adj=0.5, cex=1.5)
mtext(text = "genes", side=4, line=1, at=-0.4, adj=0.5, cex=1.5)
mtext(text = "(e) Simulated expression data (log scale)", side=1,
      line=3.5, at=0, adj=0.5, cex=1.75)


## ---- include = FALSE---------------------------------------------
set.seed(9000)

## ----simulation_graph_converging_inhibiting_hide, fig.show='hold', out.width = '50%', out.height  = '50%', fig.align='center', dpi=36, fig.retina=1, fig.margin = FALSE, fig.ncol = 2, warning=FALSE, message=FALSE, cache=TRUE----
# activating graph
state <- state <- c(-1, 1, -1)
plot_directed(graph_converging, state=state,
              layout = layout.kamada.kawai,
              cex.node=2, cex.arrow=4, arrow_clip = 0.2)
mtext(text = "(a) Activating pathway structure", side=1,
      line=3.5, at=0.075, adj=0.5, cex=1.75)
box()
#plot relationship matrix
heatmap.2(make_distance_graph(graph_converging, absolute = FALSE),
          scale = "none", trace = "none",
          col = colorpanel(50, "white", "red"),
          colsep = 1:length(V(graph_converging)),
          rowsep = 1:length(V(graph_converging)))
mtext(text = "(b) Relationship matrix", side=1,
      line=3.5, at=0, adj=0.5, cex=1.75)
#plot sigma matrix
heatmap.2(make_sigma_mat_dist_graph(graph_converging, state,
                                    cor = 0.8, absolute = FALSE),
          scale = "none", trace = "none",
          col = colorpanel(50, "blue", "white", "red"),
          colsep = 1:length(V(graph_converging)),
          rowsep = 1:length(V(graph_converging)))
mtext(text = expression(paste("(c) ", Sigma, " matrix")), side=1,
      line=3.5, at=0, adj=0.5, cex=1.75)
#simulated data
expr <- generate_expression(100, graph_converging, state,
                            cor = 0.8, mean = 0,
                            comm = FALSE, dist =TRUE,
                            absolute = FALSE, state = state)
#plot simulated correlations
heatmap.2(cor(t(expr)), scale = "none", trace = "none",
          col = colorpanel(50, "blue", "white", "red"),
          colsep = 1:length(V(graph_converging)),
          rowsep = 1:length(V(graph_converging)))
mtext(text = "(d) Simulated correlation", side=1,
      line=3.5, at=0, adj=0.5, cex=1.75)
#plot simulated expression data
heatmap.2(expr, scale = "none", trace = "none", col = bluered(50),
          colsep = 1:length(V(graph_converging)),
          rowsep = 1:length(V(graph_converging)), labCol = "")
mtext(text = "samples", side=1, line=1.5, at=0.2, adj=0.5, cex=1.5)
mtext(text = "genes", side=4, line=1, at=-0.4, adj=0.5, cex=1.5)
mtext(text = "(e) Simulated expression data (log scale)", side=1,
      line=3.5, at=0, adj=0.5, cex=1.75)


## ---- include = FALSE---------------------------------------------
set.seed(9000)

## ----simulation_graph_reconnecting_hide, fig.show='hold', out.width = '50%', out.height  = '50%', fig.align='center', dpi=36, fig.retina=1, fig.margin = FALSE, fig.ncol = 2, warning=FALSE, message=FALSE, cache=TRUE----
graph_reconnecting_edges <- rbind(c("A", "B"), c("B", "C"), c("B", "D"),
                                  c("C", "E"), c("D", "E"), c("E", "F"))
graph_reconnecting <- graph.edgelist(graph_reconnecting_edges, directed = T)

# activating graph
state <- rep(1, length(E(graph_reconnecting)))
plot_directed(graph_reconnecting, state=state,
              layout = layout.kamada.kawai,
              cex.node=2, cex.arrow=4, arrow_clip = 0.2)
mtext(text = "(a) Activating pathway structure", side=1,
      line=3.5, at=0.075, adj=0.5, cex=1.75)
box()
#plot relationship matrix
heatmap.2(make_distance_graph(graph_reconnecting, absolute = FALSE),
          scale = "none", trace = "none",
          col = colorpanel(50, "white", "red"),
          colsep = 1:length(V(graph_reconnecting)),
          rowsep = 1:length(V(graph_reconnecting)))
mtext(text = "(b) Relationship matrix", side=1,
      line=3.5, at=0, adj=0.5, cex=1.75)
#plot sigma matrix
heatmap.2(make_sigma_mat_dist_graph(graph_reconnecting, cor = 0.8,
                                    absolute = FALSE),
          scale = "none", trace = "none",
          col = colorpanel(50, "white", "red"),
          colsep = 1:length(V(graph_reconnecting)),
          rowsep = 1:length(V(graph_reconnecting)))
mtext(text = expression(paste("(c) ", Sigma, " matrix")), side=1,
      line=3.5, at=0, adj=0.5, cex=1.75)
#simulated data
expr <- generate_expression(100, graph_reconnecting,
                            cor = 0.8, mean = 0,
                            comm = FALSE, dist =TRUE,
                            absolute = FALSE, state = state)
#plot simulated correlations
heatmap.2(cor(t(expr)), scale = "none", trace = "none",
          col = colorpanel(50, "white", "red"),
          colsep = 1:length(V(graph_reconnecting)),
          rowsep = 1:length(V(graph_reconnecting)))
mtext(text = "(d) Simulated correlation", side=1,
      line=3.5, at=0, adj=0.5, cex=1.75)
#plot simulated expression data
heatmap.2(expr, scale = "none", trace = "none", col = bluered(50),
          colsep = 1:length(V(graph_reconnecting)),
          rowsep = 1:length(V(graph_reconnecting)), labCol = "")
mtext(text = "samples", side=1, line=1.5, at=0.2, adj=0.5, cex=1.5)
mtext(text = "genes", side=4, line=1, at=-0.4, adj=0.5, cex=1.5)
mtext(text = "(e) Simulated expression data (log scale)", side=1,
      line=3.5, at=0, adj=0.5, cex=1.75)


## ---- include = FALSE---------------------------------------------
set.seed(9000)

## ----simulation_graph_reconnecting_inhibiting_hide, fig.show='hold', out.width = '50%', out.height  = '50%', fig.align='center', dpi=36, fig.retina=1, fig.margin = FALSE, fig.ncol = 2, warning=FALSE, message=FALSE, cache=TRUE----
# activating graph
state <- state <- c(1, 1, -1, -1, 1, 1, 1, 1)
plot_directed(graph_reconnecting, state=state,
              layout = layout.kamada.kawai,
              cex.node=2, cex.arrow=4, arrow_clip = 0.2)
mtext(text = "(a) Activating pathway structure", side=1,
      line=3.5, at=0.075, adj=0.5, cex=1.75)
box()
#plot relationship matrix
heatmap.2(make_distance_graph(graph_reconnecting, absolute = FALSE),
          scale = "none", trace = "none",
          col = colorpanel(50, "white", "red"),
          colsep = 1:length(V(graph_reconnecting)),
          rowsep = 1:length(V(graph_reconnecting)))
mtext(text = "(b) Relationship matrix", side=1, line=3.5, at=0, adj=0.5, cex=1.75)
#plot sigma matrix
heatmap.2(make_sigma_mat_dist_graph(graph_reconnecting, state,
                                    cor = 0.8, absolute = FALSE),
          scale = "none", trace = "none",
          col = colorpanel(50, "blue", "white", "red"),
          colsep = 1:length(V(graph_reconnecting)),
          rowsep = 1:length(V(graph_reconnecting)))
mtext(text = expression(paste("(c) ", Sigma, " matrix")), side=1,
      line=3.5, at=0, adj=0.5, cex=1.75)
#simulated data
expr <- generate_expression(100, graph_reconnecting, state,
                            cor = 0.8, mean = 0,
                            comm = FALSE, dist =TRUE,
                            absolute = FALSE, state = state)
#plot simulated correlations
heatmap.2(cor(t(expr)), scale = "none", trace = "none",
          col = colorpanel(50, "blue", "white", "red"),
          colsep = 1:length(V(graph_reconnecting)),
          rowsep = 1:length(V(graph_reconnecting)))
mtext(text = "(d) Simulated correlation", side=1,
      line=3.5, at=0, adj=0.5, cex=1.75)
#plot simulated expression data
heatmap.2(expr, scale = "none", trace = "none", col = bluered(50),
          colsep = 1:length(V(graph_reconnecting)),
          rowsep = 1:length(V(graph_reconnecting)), labCol = "")
mtext(text = "samples", side=1, line=1.5, at=0.2, adj=0.5, cex=1.5)
mtext(text = "genes", side=4, line=1, at=-0.4, adj=0.5, cex=1.5)
mtext(text = "(e) Simulated expression data (log scale)", side=1,
      line=3.5, at=0, adj=0.5, cex=1.75)


## ----  fig.align='center', out.width="80%", fig.height = 5, fig.width = 5, eval = FALSE, echo = FALSE, warning=FALSE, message = FALSE----
#  #RAF_MAP_graph <- identity(RAF_MAP_graph)
#  #plot_directed(graph, col.arrow = "#00A9FF", fill.node = "lightblue")

## ---- include = FALSE---------------------------------------------
set.seed(9000)

## ----simulation_RAF_MAP_graph_hide, fig.show='hold', out.width = '50%', out.height  = '50%', fig.align='center', dpi=36, fig.retina=1, fig.margin = FALSE, fig.ncol = 2, warning=FALSE, message=FALSE----
RAF_MAP_graph <- identity(RAF_MAP_graph)

# activating graph
state <- rep(1, length(E(RAF_MAP_graph)))
plot_directed(RAF_MAP_graph, state=state,
              layout = layout.kamada.kawai,
              cex.node=2, cex.arrow=4, arrow_clip = 0.2,
              col.arrow = "#00A9FF", fill.node = "lightblue")
mtext(text = "(a) Activating pathway structure", side=1,
      line=3.5, at=0.075, adj=0.5, cex=1.75)
box()
#plot relationship matrix
heatmap.2(make_distance_graph(RAF_MAP_graph, absolute = FALSE),
          scale = "none", trace = "none",
          col = colorpanel(50, "white", "red"),
          colsep = 1:length(V(RAF_MAP_graph)),
          rowsep = 1:length(V(RAF_MAP_graph)))
mtext(text = "(b) Relationship matrix", side=1,
      line=3.5, at=0, adj=0.5, cex=1.75)
#plot sigma matrix
heatmap.2(make_sigma_mat_dist_graph(RAF_MAP_graph, cor = 0.8,
                                    absolute = FALSE),
          scale = "none", trace = "none",
          col = colorpanel(50, "white", "red"),
          colsep = 1:length(V(RAF_MAP_graph)),
          rowsep = 1:length(V(RAF_MAP_graph)))
mtext(text = expression(paste("(c) ", Sigma, " matrix")), side=1,
      line=3.5, at=0, adj=0.5, cex=1.75)
#simulated data
expr <- generate_expression(100, RAF_MAP_graph,
                            cor = 0.8, mean = 0,
                            comm = FALSE, dist =TRUE,
                            absolute = FALSE, state = state)
#plot simulated correlations
heatmap.2(cor(t(expr)), scale = "none", trace = "none",
          col = colorpanel(50, "white", "red"),
          colsep = 1:length(V(RAF_MAP_graph)),
          rowsep = 1:length(V(RAF_MAP_graph)))
mtext(text = "(d) Simulated correlation", side=1,
      line=3.5, at=0, adj=0.5, cex=1.75)
#plot simulated expression data
heatmap.2(expr, scale = "none", trace = "none", col = bluered(50),
          colsep = 1:length(V(RAF_MAP_graph)),
          rowsep = 1:length(V(RAF_MAP_graph)), labCol = "")
mtext(text = "samples", side=1, line=1.5, at=0.2, adj=0.5, cex=1.5)
mtext(text = "genes", side=4, line=1, at=-0.4, adj=0.5, cex=1.5)
mtext(text = "(e) Simulated expression data (log scale)", side=1,
      line=3.5, at=0, adj=0.5, cex=1.75)


## ---- include = FALSE---------------------------------------------
set.seed(9000)

## ----simulation_Pi3K_graph_hide, fig.show='hold', out.width = '50%', out.height  = '50%', fig.align='center', dpi=36, fig.retina=1, fig.margin = FALSE, fig.ncol = 2, warning=FALSE, message=FALSE, cache=TRUE----
graph <- identity(Pi3K_graph)

# activating graph
state <- rep(1, length(E(Pi3K_graph)))
plot_directed(Pi3K_graph, state=state,
              layout = layout.kamada.kawai,
              cex.node=2, cex.arrow=4, arrow_clip = 0.2,
              col.arrow = "#00A9FF", fill.node = "lightblue")
mtext(text = "(a) Activating pathway structure", side=1,
      line=3.5, at=0.075, adj=0.5, cex=1.75)
box()
#plot relationship matrix
heatmap.2(make_distance_graph(Pi3K_graph, absolute = FALSE),
          scale = "none", trace = "none",
          col = colorpanel(50, "white", "red"),
          colsep = 1:length(V(Pi3K_graph)),
          rowsep = 1:length(V(Pi3K_graph)))
mtext(text = "(b) Relationship matrix", side=1,
      line=3.5, at=0, adj=0.5, cex=1.75)
#plot sigma matrix
heatmap.2(make_sigma_mat_dist_graph(Pi3K_graph, cor = 0.8,
                                    absolute = FALSE),
          scale = "none", trace = "none",
          col = colorpanel(50, "white", "red"),
          colsep = 1:length(V(Pi3K_graph)),
          rowsep = 1:length(V(Pi3K_graph)))
mtext(text = expression(paste("(c) ", Sigma, " matrix")), side=1,
      line=3.5, at=0, adj=0.5, cex=1.75)
#simulated data
expr <- generate_expression(100, Pi3K_graph, 
                            cor = 0.8, mean = 0,
                            comm = FALSE, dist =TRUE,
                            absolute = FALSE, state = state)
#plot simulated correlations
heatmap.2(cor(t(expr)), scale = "none", trace = "none",
          col = colorpanel(50, "white", "red"),
          colsep = 1:length(V(Pi3K_graph)),
          rowsep = 1:length(V(Pi3K_graph)))
mtext(text = "(d) Simulated correlation", side=1, line=3.5, at=0, adj=0.5, cex=1.75)
#plot simulated expression data
heatmap.2(expr, scale = "none", trace = "none", col = bluered(50),
          colsep = 1:length(V(Pi3K_graph)),
          rowsep = 1:length(V(Pi3K_graph)), labCol = "")
mtext(text = "samples", side=1, line=1.5, at=0.2, adj=0.5, cex=1.5)
mtext(text = "genes", side=4, line=1, at=-0.4, adj=0.5, cex=1.5)
mtext(text = "(e) Simulated expression data (log scale)", side=1,
      line=3.5, at=0, adj=0.5, cex=1.75)


## ----  fig.align='center', out.width="80%", fig.height = 5, fig.width = 5, warning=FALSE, eval = FALSE, echo = FALSE, message = FALSE----
#  Pi3K_AKT_graph <- identity(Pi3K_AKT_graph)
#  edge_properties <- E(Pi3K_AKT_graph)$state
#  plot_directed(Pi3K_AKT_graph, state = edge_properties,
#    col.arrow = c(alpha("navyblue", 0.25), alpha("red", 0.25))[edge_properties],
#    fill.node = c("lightblue"))

## ---- include = FALSE---------------------------------------------
set.seed(9000)

## ----simulation_Pi3K_AKT_graph_hide, fig.show='hold', out.width = '50%', out.height  = '50%', fig.align='center', dpi=36, fig.retina=1, fig.margin = FALSE, fig.ncol = 2, warning=FALSE, message=FALSE, cache=TRUE, eval = FALSE----
#  Pi3K_AKT_graph <- identity(Pi3K_AKT_graph)
#  Pi3K_AKT_graph <- simplify(Pi3K_AKT_graph,
#        edge.attr.comb = function(x){
#           ifelse(any(x %in% list(-1, 2, "inhibiting", "inhibition")), 2, 1)
#          })
#  Pi3K_AKT_graph <- simplify(Pi3K_AKT_graph, edge.attr.comb = "first")
#  edge_properties <- E(Pi3K_AKT_graph)$state
#  
#  # activating graph
#  #state <- rep(1, length(E(Pi3K_AKT_graph)))
#  plot_directed(Pi3K_AKT_graph, state = edge_properties,
#                layout = layout.kamada.kawai,
#                cex.node=2, cex.arrow=4, arrow_clip = 0.2,
#                col.arrow = c(alpha("navyblue", 0.25),
#                              alpha("red", 0.25))[edge_properties],
#                fill.node = "lightblue")
#  mtext(text = "(a) Activating pathway structure", side=1,
#        line=3.5, at=0.075, adj=0.5, cex=1.75)
#  box()
#  #plot relationship matrix
#  heatmap.2(make_distance_graph(Pi3K_AKT_graph, absolute = FALSE),
#            scale = "none", trace = "none",
#            col = colorpanel(50, "white", "red"))
#  mtext(text = "(b) Relationship matrix", side=1, line=3.5, at=0, adj=0.5, cex=1.75)
#  #plot sigma matrix
#  heatmap.2(make_sigma_mat_dist_graph(Pi3K_AKT_graph, state = edge_properties,
#                                      cor = 0.8, absolute = FALSE),
#            scale = "none", trace = "none",
#            col = colorpanel(50, "blue", "white", "red"))
#  mtext(text = expression(paste("(c) ", Sigma, " matrix")), side=1,
#        line=3.5, at=0, adj=0.5, cex=1.75)
#  #simulated data
#  expr <- generate_expression(100, Pi3K_AKT_graph, cor = 0.8, mean = 0,
#                              comm = FALSE, dist =TRUE, absolute = FALSE,
#                              state = edge_properties)
#  #plot simulated correlations
#  heatmap.2(cor(t(expr)), scale = "none", trace = "none",
#            col = colorpanel(50, "blue", "white", "red"))
#  mtext(text = "(d) Simulated correlation", side=1,
#        line=3.5, at=0, adj=0.5, cex=1.75)
#  #plot simulated expression data
#  heatmap.2(expr, scale = "none", trace = "none",
#            col = bluered(50), labCol = "")
#  mtext(text = "samples", side=1, line=1.5, at=0.2, adj=0.5, cex=1.5)
#  mtext(text = "genes", side=4, line=1, at=-0.4, adj=0.5, cex=1.5)
#  mtext(text = "(e) Simulated expression data (log scale)", side=1,
#        line=3.5, at=0, adj=0.5, cex=1.75)
#  

## ----  fig.align='center', out.width="80%", fig.height = 5, fig.width = 5----
TGFBeta_Smad_graph <- identity(TGFBeta_Smad_graph)
edge_properties <- E(TGFBeta_Smad_graph)$state
plot_directed(TGFBeta_Smad_graph, state = edge_properties,
              col.arrow = c(alpha("navyblue", 0.25),
                            alpha("red", 0.25))[edge_properties],
              fill.node = c("lightblue"))

## ---- include = FALSE---------------------------------------------
set.seed(9000)

## ----simulation_TGFBeta_Smad_graph_hide, fig.show='hold', out.width = '50%', out.height  = '50%', fig.align='center', dpi=36, fig.retina=1, fig.margin = FALSE, fig.ncol = 2, warning=FALSE, message=FALSE, cache=TRUE----
TGFBeta_Smad_graph <- identity(TGFBeta_Smad_graph)
edge_properties <- E(TGFBeta_Smad_graph)$state

# activating graph
plot_directed(TGFBeta_Smad_graph, state = edge_properties,
              layout = layout.kamada.kawai,
              cex.node=2, cex.arrow=4, arrow_clip = 0.2,
              col.arrow = c(alpha("navyblue", 0.25),
                            alpha("red", 0.25))[edge_properties],
              fill.node = "lightblue")
mtext(text = "(a) Activating pathway structure", side=1,
      line=3.5, at=0.075, adj=0.5, cex=1.75)
box()
#plot relationship matrix
heatmap.2(make_distance_graph(TGFBeta_Smad_graph, absolute = FALSE),
          scale = "none", trace = "none",
          col = colorpanel(50, "white", "red"),
          colsep = 1:length(V(TGFBeta_Smad_graph)),
          rowsep = 1:length(V(TGFBeta_Smad_graph)))
mtext(text = "(b) Relationship matrix", side=1,
      line=3.5, at=0, adj=0.5, cex=1.75)
#plot sigma matrix
heatmap.2(make_sigma_mat_dist_graph(TGFBeta_Smad_graph,
                                    state = edge_properties,
                                    cor = 0.8, absolute = FALSE),
          scale = "none", trace = "none",
          col = colorpanel(50, "blue", "white", "red"),
          colsep = 1:length(V(TGFBeta_Smad_graph)),
          rowsep = 1:length(V(TGFBeta_Smad_graph)))
mtext(text = expression(paste("(c) ", Sigma, " matrix")), side=1,
      line=3.5, at=0, adj=0.5, cex=1.75)
#simulated data
expr <- generate_expression(100, TGFBeta_Smad_graph,
                            cor = 0.8, mean = 0,
                            comm = FALSE, dist =TRUE,
                            absolute = FALSE, state = edge_properties)
#plot simulated correlations
heatmap.2(cor(t(expr)), scale = "none", trace = "none",
          col = colorpanel(50, "blue", "white", "red"),
          colsep = 1:length(V(TGFBeta_Smad_graph)),
          rowsep = 1:length(V(TGFBeta_Smad_graph)))
mtext(text = "(d) Simulated correlation", side=1,
      line=3.5, at=0, adj=0.5, cex=1.75)
#plot simulated expression data
heatmap.2(expr, scale = "none", trace = "none", col = bluered(50),
          colsep = 1:length(V(TGFBeta_Smad_graph)),
          rowsep = 1:length(V(TGFBeta_Smad_graph)), labCol = "")
mtext(text = "samples", side=1, line=1.5, at=0.2, adj=0.5, cex=1.5)
mtext(text = "genes", side=4, line=1, at=-0.4, adj=0.5, cex=1.5)
mtext(text = "(e) Simulated expression data (log scale)", side=1,
      line=3.5, at=0, adj=0.5, cex=1.75)


## ----echo=TRUE, message=TRUE, warning=FALSE-----------------------
citation("graphsim")

## ---- echo = FALSE------------------------------------------------
sessionInfo()
```
