## Supplementary material for "graphsim: An R package for simulating gene expression data from graph structures of biological pathways": graphsim 1.0.0 R package: NEWS.md

# graphsim 1.0.0

* Major stable release: note changes to results are possible! (legacy code should run without breaking)

* Expanded documentation and examples

* Resolves errors handling inhibiting edges

* Efficiently compute a state matrix from a vector of edge properties from paths

* Enables passing "sd" (standard deviation) to alter covariance of Sigma matrix

* Adds methods for computing using Laplacian matrices

* Adds function to compute simulations directly from an adjacency matrix

* Migrates computing states to sigma (these matrices include inhibitions)

# graphsim 0.1.1

* Initial CRAN release

* Unit testing for all functions

* Checking for compatible inputs

* Passing layout parameters to plotting function

# graphsim 0.1.1

* Full vignette of biological pathway

* Merge plot_directed from function from plot.igraph to avoid github dependencies

* Added example data for reactome pathways

# graphsim 0.1.0

* Initial version of the package for generating simulated data
