## Supplementary material for "graphsim: An R package for simulating gene expression data from graph structures of biological pathways": graphsim 1.0.0 R package: adjmat.R

```
##' @name make_adjmatrix
##' @aliases make_adjmatrix_graph
##' @rdname make_adjmatrix_graph
##'
##' @title Generate Adjacency Matrix
##' 
##' @description Compute the adjacency matrix of a (directed) \code{\link[igraph]{igraph}}
##' structure, preserving node/column/row names (and direction).
##'
##' @param graph An \code{\link[igraph]{igraph}} object. May be directed or weighted.
##' @param directed logical. Whether directed information is passed to the adjacency matrix.
##' @keywords graph network igraph adjacency
##' @importFrom igraph as_adjacency_matrix graph.edgelist as.undirected
##' @import igraph
##' @examples 
##' 
##' # construct a synthetic graph module
##' library("igraph")
##' graph_test_edges <- rbind(c("A", "B"), c("B", "C"), c("B", "D"))
##' graph_test <- graph.edgelist(graph_test_edges, directed = TRUE)
##' 
##' # compute adjacency matrix for toy example
##' adjacency_matrix <- make_adjmatrix_graph(graph_test)
##' adjacency_matrix
##' 
##' # construct a synthetic graph network
##' graph_structure_edges <- rbind(c("A", "C"), c("B", "C"), c("C", "D"), c("D", "E"),
##'                                c("D", "F"), c("F", "G"), c("F", "I"), c("H", "I"))
##' graph_structure <- graph.edgelist(graph_structure_edges, directed = TRUE)
##' # compute adjacency matrix for toy network
##' graph_structure_adjacency_matrix <- make_adjmatrix_graph(graph_structure)
##' graph_structure_adjacency_matrix
##' 
##' # import graph from package for reactome pathway
##' # TGF-\eqn{\Beta} receptor signaling activates SMADs (R-HSA-2173789)
##' TGFBeta_Smad_graph <- identity(TGFBeta_Smad_graph)
##' 
##' # compute adjacency matrix for TGF-\eqn{\Beta} receptor signaling activates SMADs
##' TGFBeta_Smad_adjacency_matrix <- make_adjmatrix_graph(TGFBeta_Smad_graph)
##' dim(TGFBeta_Smad_adjacency_matrix)
##' TGFBeta_Smad_adjacency_matrix[1:12, 1:12]
##' 
##' @return An adjacency matrix compatible with generating an expression matrix
##' 
##' @export make_adjmatrix_graph
make_adjmatrix_graph <- function(graph, directed = FALSE){
  if(directed == FALSE) graph <- as.undirected(graph)
  adj_mat <- as.matrix(as_adjacency_matrix(graph))
  rownames(adj_mat) <- colnames(adj_mat) <- names(V(graph))
  return(adj_mat)
}
```
