## Supplementary material for "graphsim: An R package for simulating gene expression data from graph structures of biological pathways": graphsim 1.0.0 R package: commonlink.R

```
##' @name make_commonlink
##' @rdname make_commonlink
##'
##' @title Generate Common Link Matrix
##'
##' @description Compute the common link matrix of a (directed) \code{\link[igraph]{igraph}}
##' structure, preserving node / column / row names (and direction). We can compute the common 
##' inks between each pair of nodes. This shows how many nodes are mutually connected to both
##' of the nodes in the matrix (how many paths of length 2 exist between them).
##'
##' @param adj_mat precomputed adjacency matrix.
##' @param graph An \code{\link[igraph]{igraph}} object. May be directed or weighted.
##' @param directed logical. Whether directed information is passed to the adjacency matrix.
##' @keywords graph network igraph neighbourhood
##' @import igraph
##' @examples 
##' 
##' # construct a synthetic graph module
##' library("igraph")
##' graph_test_edges <- rbind(c("A", "B"), c("B", "C"), c("B", "D"))
##' graph_test <- graph.edgelist(graph_test_edges, directed = TRUE)
##' 
##' # compute adjacency matrix for toy example
##' adjacency_matrix <- make_adjmatrix_graph(graph_test)
##' # compute nodes with shared edges to a 3rd node
##' common_link_matrix <- make_commonlink_adjmat(adjacency_matrix)
##' common_link_matrix
##' 
##' # construct a synthetic graph network
##' graph_structure_edges <- rbind(c("A", "C"), c("B", "C"), c("C", "D"), c("D", "E"),
##'                                c("D", "F"), c("F", "G"), c("F", "I"), c("H", "I"))
##' graph_structure <- graph.edgelist(graph_structure_edges, directed = TRUE)
##' # compute adjacency matrix for toy network
##' graph_structure_adjacency_matrix <- make_adjmatrix_graph(graph_structure)
##' # compute nodes with shared edges to a 3rd node
##' graph_structure_common_link_matrix <- make_commonlink_adjmat(graph_structure_adjacency_matrix)
##' graph_structure_common_link_matrix
##' 
##' # import graph from package for reactome pathway
##' # TGF-\eqn{\Beta} receptor signaling activates SMADs (R-HSA-2173789)
##' TGFBeta_Smad_graph <- identity(TGFBeta_Smad_graph)
##' # compute nodes with shared edges to a 3rd node 
##' TGFBeta_Smad_adjacency_matrix <- make_adjmatrix_graph(TGFBeta_Smad_graph)
##' TGFBeta_Smad_common_link_matrix <- make_commonlink_adjmat(TGFBeta_Smad_adjacency_matrix)
##' # we show summary statistics as the graph is large
##' dim(TGFBeta_Smad_common_link_matrix)
##' TGFBeta_Smad_common_link_matrix[1:12, 1:12]
##' # visualise matrix
##' library("gplots")
##' heatmap.2(TGFBeta_Smad_common_link_matrix, scale = "none", trace = "none",
##'           col = colorpanel(50, "white", "red"))
##' 
##' @return An integer matrix of number of links shared between nodes
##' 
##' @export
make_commonlink_adjmat <- function(adj_mat){
  comm_mat <- matrix(NA, nrow(adj_mat), ncol(adj_mat))
  for(ii in 1:nrow(adj_mat)){
    for(jj in 1:ncol(adj_mat)){
      comm_mat[ii, jj] <- sum(adj_mat[ii,] & adj_mat[,jj])
    }
  }
  rownames(comm_mat) <- rownames(adj_mat)
  colnames(comm_mat) <- colnames(adj_mat)
  return(comm_mat)
}

##' @rdname make_commonlink
##' @export
make_commonlink_graph <- function(graph, directed = FALSE){
  adj_mat <- make_adjmatrix_graph(graph, directed = directed)
  comm_mat <- make_commonlink_adjmat(adj_mat)
  return(comm_mat)
}
```
