## Supplementary material for "graphsim: An R package for simulating gene expression data from graph structures of biological pathways": graphsim 1.0.0 R package: generate.R

```
##' @name generate_expression
##' @rdname generate_expression
##'
##' @title Generate Simulated Expression
##'
##' @description Compute simulated continuous expression data from a graph 
##' network structure. Requires an \code{\link[igraph]{igraph}} pathway 
##' structure and a matrix of states (1 for activating and -1 for 
##' inhibiting) for link signed correlations, from a vector of edge states 
##' to a signed adjacency matrix for use in 
##' \code{\link[graphsim]{generate_expression}}. 
##' Uses graph structure to pass a sigma covariance matrix from 
##' \code{\link[graphsim]{make_sigma_mat_dist_graph}} or 
##' \code{\link[graphsim]{make_sigma_mat_graph}} on to 
##' \code{\link[mvtnorm]{rmvnorm}}. By default data is generated with a mean of
##'  0 and standard deviation of 1 for each gene (with correlations between 
##'  derived from the graph structure).
##'
##' @param n number of observations (simulated samples).
##' @param mat precomputed adjacency, laplacian, commonlink, or scaled 
##' distance matrix.
##' @param graph An \code{\link[igraph]{igraph}} object. May must be 
##' directed if states are used.
##' @param state numeric vector. Vector of length E(graph). Sign used
##' to calculate state matrix, may be an integer state or inferred directly
##' from expected correlations for each edge. May be applied a scalar across
##' all edges or as a vector for each edge respectively. May also be entered
##' as text for "activating" or "inhibiting" or as integers for activating (0,1)
##' or inhibiting (-1,2). Compatible with inputs for \code{\link[graphsim]{plot_directed}}.
##' Also takes a pre-computed state matrix from \code{\link[graphsim]{make_state_matrix}}
##' if applied to the same graph multiple times.
##' @param cor numeric. Simulated maximum correlation/covariance of two 
##' adjacent nodes. Default to 0.8.
##' @param mean mean value of each simulated gene. Defaults to 0.
##' May be entered as a scalar applying to 
##' all genes or a vector with a separate value for each.
##' @param sd	 standard deviations of each gene. Defaults to 1.
##' May be entered as a scalar applying to 
##' all genes or a vector with a separate value for each.
##' @param dist logical. Whether a graph distance 
##' (\code{\link[graphsim]{make_sigma_mat_dist_graph}}) or derived matrix
##'  (\code{\link[graphsim]{make_sigma_mat_graph}}) is used to compute the
##'   sigma matrix.
##' @param comm,absolute,laplacian logical. Parameters for Sigma matrix
##' generation. Passed on to \code{\link[graphsim]{make_sigma_mat_dist_graph}} 
##' or \code{\link[graphsim]{make_sigma_mat_graph}}.
##' @keywords graph network igraph mvtnorm simulation
##' @importFrom  mvtnorm rmvnorm
##' @importFrom igraph as_adjacency_matrix graph.edgelist
##' @import igraph
##' @importFrom igraph is_igraph E V
##' @importFrom Matrix nearPD
##' @importFrom matrixcalc is.symmetric.matrix is.positive.definite
##' @importFrom gplots heatmap.2
##' @examples
##' 
##' # construct a synthetic graph module
##' library("igraph")
##' graph_test_edges <- rbind(c("A", "B"), c("B", "C"), c("B", "D"))
##' graph_test <- graph.edgelist(graph_test_edges, directed = TRUE)
##' 
##' # compute a simulated dataset for toy example
##' # n = 100 samples
##' # cor = 0.8 max correlation between samples
##' # absolute = FALSE (geometric distance by default)
##' test_data <- generate_expression(100, graph_test, cor = 0.8)
##' ##' # visualise matrix
##' library("gplots")
##' # expression data
##' heatmap.2(test_data, scale = "none", trace = "none",
##'           col = colorpanel(50, "blue", "white", "red"))
##' # correlations
##' heatmap.2(cor(t(test_data)), scale = "none", trace = "none",
##'           col = colorpanel(50, "white", "red"))
##' # expected correlations (\eqn{\Sigma})
##' sigma_matrix <- make_sigma_mat_graph(graph_test, cor = 0.8)
##' heatmap.2(make_sigma_mat_graph(graph_test, cor = 0.8),
##'           scale = "none", trace = "none", 
##'           col = colorpanel(50, "white", "red"))
##' 
##' # compute adjacency matrix for toy example
##' adjacency_matrix <- make_adjmatrix_graph(graph_test)
##' # generate simulated data from adjacency matrix input
##' test_data <- generate_expression_mat(100, adjacency_matrix, cor = 0.8)
##' 
##' # compute a simulated dataset for toy example
##' # n = 100 samples
##' # cor = 0.8 max correlation between samples
##' # absolute = TRUE (arithmetic distance)
##' test_data <- generate_expression(100, graph_test, cor = 0.8, absolute = TRUE)
##' ##' # visualise matrix
##' library("gplots")
##' # expression data
##' heatmap.2(test_data, scale = "none", trace = "none",
##'           col = colorpanel(50, "blue", "white", "red"))
##' # correlations
##' heatmap.2(cor(t(test_data)),
##'           scale = "none", trace = "none",
##'           col = colorpanel(50, "white", "red"))
##' # expected correlations (\eqn{\Sigma})
##' sigma_matrix <- make_sigma_mat_graph(graph_test, cor = 0.8)
##' heatmap.2(make_sigma_mat_graph(graph_test, cor = 0.8),
##'           scale = "none", trace = "none",
##'           col = colorpanel(50, "white", "red"))
##' 
##' # construct a synthetic graph network
##' graph_structure_edges <- rbind(c("A", "C"), c("B", "C"), c("C", "D"), c("D", "E"),
##'                                c("D", "F"), c("F", "G"), c("F", "I"), c("H", "I"))
##' graph_structure <- graph.edgelist(graph_structure_edges, directed = TRUE)
##' 
##' # compute a simulated dataset for toy network
##' # n = 250 samples
##' # state = edge_state (properties of each edge)
##' # cor = 0.95 max correlation between samples
##' # absolute = FALSE (geometric distance by default)
##' edge_state <- c(1, 1, -1, 1, 1, 1, 1, -1)
##' structure_data <- generate_expression(250, graph_structure,
##'                                       state = edge_state, cor = 0.95)
##' ##' # visualise matrix
##' library("gplots")
##' # expression data
##' heatmap.2(structure_data, scale = "none", trace = "none",
##'           col = colorpanel(50, "blue", "white", "red"))
##' # correlations
##' heatmap.2(cor(t(structure_data)), scale = "none", trace = "none",
##'           col = colorpanel(50, "blue", "white", "red"))
##' # expected correlations (\eqn{\Sigma})
##' sigma_matrix <- make_sigma_mat_graph(graph_structure,
##'                                      state = edge_state, cor = 0.8)
##' heatmap.2(make_sigma_mat_graph(graph_structure,
##'                                state = edge_state, cor = 0.8),
##'           scale = "none", trace = "none",
##'           col = colorpanel(50, "blue", "white", "red"))
##' 
##' # compute adjacency matrix for toy network
##' graph_structure_adjacency_matrix <- make_adjmatrix_graph(graph_structure)
##' # define states for for each edge
##' edge_state <- c(1, 1, -1, 1, 1, 1, 1, -1)
##' # generate simulated data from adjacency matrix input
##' structure_data <- generate_expression_mat(250, graph_structure_adjacency_matrix,
##'                                           state = edge_state, cor = 0.8)
##' 
##' # compute a simulated dataset for toy network
##' # n = 1000 samples
##' # state = TGFBeta_Smad_state (properties of each edge)
##' # cor = 0.75 max correlation between samples
##' # absolute = FALSE (geometric distance by default)
##'  # compute states directly from graph attributes for TGF-\eqn{\Beta} pathway
##' TGFBeta_Smad_state <- E(TGFBeta_Smad_graph)$state
##' table(TGFBeta_Smad_state)
##' # generate simulated data
##' TGFBeta_Smad_data <- generate_expression(1000, TGFBeta_Smad_graph, cor = 0.75)
##' ##' # visualise matrix
##' library("gplots")
##' # expression data
##' heatmap.2(TGFBeta_Smad_data, scale = "none", trace = "none",
##'           col = colorpanel(50, "blue", "white", "red"))
##' # correlations
##' heatmap.2(cor(t(TGFBeta_Smad_data)), scale = "none", trace = "none",
##'           dendrogram = "none", Rowv = FALSE, Colv = FALSE,
##'           col = colorpanel(50, "blue", "white", "red"))
##' # expected correlations (\eqn{\Sigma})
##' sigma_matrix <- make_sigma_mat_dist_graph(TGFBeta_Smad_graph, cor = 0.75)
##' heatmap.2(make_sigma_mat_dist_graph(TGFBeta_Smad_graph, cor = 0.75),
##'           scale = "none", trace = "none",
##'           dendrogram = "none", Rowv = FALSE, Colv = FALSE,
##'           col = colorpanel(50, "blue", "white", "red"))
##' 
##' 
##' # generate simulated data (absolute distance and shared edges)
##' TGFBeta_Smad_data <- generate_expression(1000, TGFBeta_Smad_graph,
##'                                          cor = 0.75, absolute = TRUE, comm = TRUE)
##' ##' # visualise matrix
##' library("gplots")
##' # expression data
##' heatmap.2(TGFBeta_Smad_data, scale = "none", trace = "none",
##'           col = colorpanel(50, "blue", "white", "red"))
##' # correlations
##' heatmap.2(cor(t(TGFBeta_Smad_data)), scale = "none", trace = "none",
##'           dendrogram = "none", Rowv = FALSE, Colv = FALSE,
##'           col = colorpanel(50, "blue", "white", "red"))
##' # expected correlations (\eqn{\Sigma})
##' sigma_matrix <- make_sigma_mat_graph(TGFBeta_Smad_graph,
##'                                      cor = 0.75, comm = TRUE)
##' heatmap.2(make_sigma_mat_graph(TGFBeta_Smad_graph, cor = 0.75, comm = TRUE),
##'           scale = "none", trace = "none",
##'           dendrogram = "none", Rowv = FALSE, Colv = FALSE,
##'           col = colorpanel(50, "blue", "white", "red"))
##' 
##' @return numeric matrix of simulated data (log-normalised counts)
##' 
##' @export
generate_expression <- function(n, graph, state = NULL, cor = 0.8, mean = 0, sd = 1, comm = FALSE, dist = FALSE, absolute = FALSE, laplacian = FALSE){
  if(missing(graph)){
    warning(paste("object must be defined for graph input"))
    get(graph)
    stop()
  }
  if(!exists("graph")){
    warning(paste("object must be defined for graph input"))
    get("graph")
    stop()
  }
  if(!is_igraph(graph)) stop("graph must be an igraph class")
  if(!is.null(get.edge.attribute(graph, "state"))){
    state <- get.edge.attribute(graph, "state")
  } else {
    # add default state if not specified
    if(is.null(state)){
      state <- rep(1, length(E(graph)))
    }
  }
  if(is.null(get.edge.attribute(graph, "state"))){
    E(graph)$state <-  state
  }
  # this could also be done with igraph::simplify(graph, remove.multiple = TRUE, remove.loops = TRUE, edge.attr.comb = igraph_opt("edge.attr.comb"))
  ## to do: migrate to this
  graph <- as.undirected(graph, mode = "collapse", edge.attr.comb = function(x) ifelse(any(x %in% list(-1, 2, "inhibiting", "inhibition")), -1, 1))
  # remove duplicate edges (and corresponding states)
  graph <- as.directed(graph, mode = "arbitrary")
  state <- get.edge.attribute(graph, "state")
  if(!is.integer(n)){
    if(is.numeric(n)){
      if(floor(n) == n){
        n <- as.integer(n)
      } else{
        n <- floor(n)
        print(paste("rounding to sample size", n))
      }
    } else{
      stop("sample size n must be an integer of length 1")
    }
  }
  if(length(n) > 1) stop("sample size n must be an integer of length 1")
  if(length(state) == 1) state <- rep(state, length(E(graph)))
  if(!(is.vector(mean)) || length(mean) == 1 ) mean <- rep(mean,length(V(graph)))
  if(!(is.vector(sd)) || length(sd) == 1 ) sd <- rep(sd,length(V(graph)))
  if(dist){
    sig <- make_sigma_mat_dist_graph(graph, state = state, cor, sd = sd, absolute = absolute)
  } else {
    sig <- make_sigma_mat_graph(graph, state = state, cor, sd = sd, comm = comm, laplacian = laplacian)
  }
  if(is.symmetric.matrix(sig) == FALSE) {
    warning("sigma matrix was not positive definite, nearest approximation used.")
    sig <- as.matrix(nearPD(sig, corr=T, keepDiag = TRUE)$mat) #postive definite correction
  }
  if(is.positive.definite(sig) == FALSE) {
    warning("sigma matrix was not positive definite, nearest approximation used.")
    sig <- as.matrix(nearPD(sig, corr=T, keepDiag = TRUE)$mat) #postive definite correction
  }
  expr_mat <- t(rmvnorm(n,mean=mean, sigma=sig))
  rownames(expr_mat) <- names(V(graph))
  colnames(expr_mat) <- paste0("sample_", 1:n)
  return(expr_mat)
}


##' @rdname generate_expression
##' @importFrom igraph graph_from_adjacency_matrix set_edge_attr E
##' @export
generate_expression_mat <- function(n, mat, state = NULL, cor = 0.8, mean = 0, sd = 1, comm = FALSE, dist = FALSE, absolute = FALSE, laplacian = FALSE){
  if(is.null(state)){
    state <- 1
  }
  if(!is.integer(n)){
    if(is.numeric(n)){
      if(floor(n) == n){
        n <- as.integer(n)
      } else{
        n <- floor(n)
        print(paste("rounding to sample size", n))
      }
    } else{
      stop("sample size n must be an integer of length 1")
    }
  }
  if(length(n) > 1) stop("sample size n must be an integer of length 1")
  if(!(is.vector(mean)) || length(mean) == 1 ) mean <- rep(mean,ncol(mat))
  if(!(is.vector(sd)) || length(sd) == 1 ) sd <- rep(sd,ncol(mat))
  if(!is.matrix(mat)) stop("graph must be an igraph class")
  if(is.vector(state) || length(state) == 1){
    graph <- graph_from_adjacency_matrix(mat, mode = "undirected")
    if(length(state) == 1) state <- rep(state, length(E(graph)))
    graph <- set_edge_attr(graph, "state", value = state)
  }
  if(!(is.vector(mean)) || length(mean) == 1 ) mean <- rep(mean,ncol(mat))
  if(dist){
    distmat <- make_distance_adjmat(mat)
    sig <- make_sigma_mat_dist_adjmat(distmat, state = state, cor, sd = sd, absolute = absolute)
  } else {
    if(comm && laplacian){
      warning("Error: only one of commonlink or laplacian can be used")
      stop()
    }
    if(!comm && !laplacian) sig <- make_sigma_mat_adjmat(mat, state = state, cor)
    if(comm){
      mat <- make_commonlink_adjmat(mat)
      sig <- make_sigma_mat_comm(mat, state = state, cor, sd = sd)
    }
    if(laplacian){
      mat <- make_laplacian_adjmat(mat)
      sig <- make_sigma_mat_laplacian(mat, state = state, cor, sd = sd)
    }
  }
  if(is.symmetric.matrix(sig) == FALSE) {
    warning("sigma matrix was not positive definite, nearest approximation used.")
    sig <- as.matrix(nearPD(sig, corr=T, keepDiag = TRUE)$mat) #postive definite correction
  }
  if(is.positive.definite(sig) == FALSE) {
    warning("sigma matrix was not positive definite, nearest approximation used.")
    sig <- as.matrix(nearPD(sig, corr=T, keepDiag = TRUE)$mat) #postive definite correction
  }
  expr_mat <- t(rmvnorm(n,mean=mean, sigma=sig))
  rownames(expr_mat) <- names(colnames(mat))
  colnames(expr_mat) <- paste0("sample_", 1:n)
  return(expr_mat)
}
```
