## Supplementary material for "graphsim: An R package for simulating gene expression data from graph structures of biological pathways": graphsim 1.0.0 R package: pi3k_akt_graph.R

```
#' PI3K/AKT activation
#'
#' Reactome pathway R-HSA-198203 for the interactions in the phosphoinositide-3-kinase activation of Protein kinase B (PKB), also known as Akt
#' 
#' @format A graph object of 275 vertices and 21106 edges:
#' \describe{
#'   \item{V}{gene symbol (human)}
#'   \item{E}{directed relationship for pathway}
#'   \item{state}{type of relationship (activating or inhibiting) as edge attribute}
#' }
#' @source PathwayCommons \url{https://reactome.org/content/detail/R-HSA-198203}
"Pi3K_AKT_graph"
```
