## Supplementary material for "graphsim: An R package for simulating gene expression data from graph structures of biological pathways": graphsim 1.0.0 R package: pi3k_graph.R

```
#' PI3K Cascade
#'
#' Reactome pathway R-HSA-109704 for the interactions in the phosphoinositide-3-kinase cascade
#' 
#' @format A graph object of 35 vertices and 251 edges:
#' \describe{
#'   \item{V}{gene symbol (human)}
#'   \item{E}{directed relationship for pathway}
#'   \item{state}{type of relationship (activating or inhibiting) as edge attribute}
#' }
#' @source PathwayCommons \url{https://reactome.org/content/detail/R-HSA-109704}
"Pi3K_graph"
```
