## Supplementary material for "graphsim: An R package for simulating gene expression data from graph structures of biological pathways": graphsim 1.0.0 R package: plot_directed.R

```
# You can learn more about package authoring with RStudio at:
#
#   http://r-pkgs.had.co.nz/
#
# Some useful keyboard shortcuts for package authoring:
#
#   Build and Reload Package:  'Ctrl + Shift + B'
#   Check Package:             'Ctrl + Shift + E'
#   Test Package:              'Ctrl + Shift + T'
##' @name plot_directed
##' @rdname plot_directed
##' @aliases plot.directed
##'
##' @title Extensions to igraph for Customising plots
##'
##' @description Functions to plot_directed or graph structures including customised colours, layout, states, arrows. Uses graphs functions as an extension of \code{\link[igraph]{igraph}}. Designed for plotting directed graphs.
##'
##' @param graph An \code{\link[igraph]{igraph}} object. Must be directed with known states.
##' @param state character or integer. Defaults to "activating" if no "state" edge attribute 
##' found. May be applied a scalar across all edges or as a vector for each edge respectively. 
##' Accepts non-integer values for weighted edges provided that the sign indicates whether links
##'  are activating (positive) or inhibiting (negative). May also be entered as text for 
##'  "activating" or "inhibiting" or as integers for activating (0,1) or inhibiting (-1,2). 
##'  Compatible with inputs for make_state_matrix or generate_expression_graph in the graphsim 
##'  package \url{https://github.com/TomKellyGenetics/graphsim}. Vector input is supported 
##' @param labels character vector. For labels to plot nodes. Defaults to vertex names in 
##' graph object. Entering "" would yield unlabelled nodes.
##' @param layout function. Layout function as selected from \code{\link[igraph]{layout_}}. 
##' Defaults to layout.fruchterman.reingold. Alternatives include layout.kamada.kawai, 
##' layout.reingold.tilford, layout.sugiyama, and layout.davidson.harel. A 2-column 
##' layout matrix giving x and y co-ordinates of each node can be given.
##' @param cex.node numeric. Defaults to 1.
##' @param cex.label numeric. Defaults to 0.75.
##' @param cex.main numeric. Defaults to 0.8.
##' @param cex.sub numeric. Defaults to 0.8.
##' @param cex.arrow numeric Defaults to 1.25. May take a scalar applied to all edges 
##' or a vector with values for each edge respectively.
##' @param col.label character. Specfies the colours of node labels passed to plot. 
##' Defaults to par("fg").
##' @param arrow_clip numeric Defaults to 0.075 (7.5\%).
##' @param pch parameter passed to plot. Defaults to 21. Recommends using selecting 
##' between 21-25 to preserve colour behaviour. Otherwise entire node will inherit 
##' border.node as it's colour, in which case a light colour is recommended to see labels.
##' @param border.node character. Specifies the colours of node border passed to plot.
##'  Defaults to grey33. Applies to whole node shape if pch has only one colour.
##' @param fill.node character. Specfies the colours of node fill passed to plot. 
##' Defaults to grey66.
##' @param col.arrow character. Specfies the colours of arrows passed to plot. 
##' Defaults to par("fg").  May take a scalar applied to all edges or a vector
##'  with colours for each edge respectively.
##' @param main,sub,xlab,ylab Plotting parameters to specify plot titles or axes labels
##' @param frame.plot logical. Whether to frame plot with a box. Defaults to FALSE.
##' @keywords graph igraph igraph plot
##' @import igraph graphics
##' @examples
##'
##' # generate example graphs
##' library("igraph")
##' graph_structure_edges <- rbind(c("A", "C"), c("B", "C"), c("C", "D"), c("D", "E"),
##'                            c("D", "F"), c("F", "G"), c("F", "I"), c("H", "I"))
##' graph_structure <- graph.edgelist(graph_structure_edges, directed = TRUE)
##'
##' # plots with igraph defaults
##' plot(graph_structure, layout = layout.fruchterman.reingold)
##' plot(graph_structure, layout = layout.kamada.kawai)
##'
##' # plots with scalar states
##' plot_directed(graph_structure, state="activating")
##' plot_directed(graph_structure, state="inhibiting")
##'
##' # plots with vector states
##' plot_directed(graph_structure, state = c(1, 1, 1, 1, -1, 1, 1, 1))
##' plot_directed(graph_structure, state = c(1, 1, -1, 1, -1, 1, -1, 1))
##' plot_directed(graph_structure, state = c(1, 1, -1, 1, 1, 1, 1, -1))
##' 
##' # plots states with graph attributes
##' E(graph_structure)$state <- 1
##' plot_directed(graph_structure)
##' E(graph_structure)$state <- c(1, 1, -1, 1, -1, 1, -1, 1)
##' plot_directed(graph_structure)
##'
##' # plot layout customised
##' plot_directed(graph_structure, state=c(1, 1, -1, 1, -1, 1, -1, 1), layout = layout.kamada.kawai)
##' 
##' @return base R graphics
##' 
##' @export
plot_directed <- function(graph, state = NULL, labels = NULL, layout = layout.fruchterman.reingold, cex.node = 1, cex.label = 0.75, cex.arrow=1.25, cex.main=0.8, cex.sub=0.8, arrow_clip = 0.075, pch=21, border.node="grey33", fill.node="grey66", col.label = NULL, col.arrow=NULL, main=NULL, sub=NULL, xlab="", ylab="", frame.plot=F){
  if(is.function(layout)){
    L <-  layout(graph)
  } else {
    if(is.matrix(layout) && nrow(layout) == length(V(graph)) && ncol(layout) == 2){
      L <- as.matrix(layout)
    } else {
      warning(paste0("layout must be specified as an igraph function or ", length(V(graph)), " by 2 matrix"))
    }
  }
  vs <- V(graph)
  es <- as.data.frame(get.edgelist(graph))
  Nv <- length(vs)
  Ne <- length(es[1]$V1)
  Xn <- L[,1]
  Yn <- L[,2]
  plot(Xn, Yn, xaxt="n", yaxt="n", xlab=xlab, ylab=ylab, xlim = mean(Xn)+c(-1,1 )*1.2*(max(Xn)-min(Xn))/2, ylim = mean(Yn)+c(-1,1 )*1.2*(max(Yn)-min(Yn))/2, frame.plot=frame.plot, cex = 2 * cex.node, pch=1, col=par()$bg, main=main, sub=sub, cex.main=cex.main, cex.sub=cex.sub)
  if(!is.null(get.edge.attribute(graph, "state"))){
    state <- get.edge.attribute(graph, "state")
  } else {
    # add default state if not specified
    if(is.null(state)){
      state <- "activating"
    }
  }
  if(is.numeric(state)){
    state <- as.integer(state)
    if(!all(state %in% -1:2)){
      state <- sign(state)
      warning("state inferred from non-integer weighted edges")
    }
    if(all(state %in% -1:2)){
      state[state == -1] <- 2
      state[state == 0] <- 1
      if(is.null(col.arrow)){
        col.arrow <- c(par("fg"), "red")[state]
      }
      state <- c("activating", "inhibiting")[state]
    } else {
      state <- sign(state) # coerce to vector or 1 and -1 if not already
      warning("Please give numeric states as integers: 0 or 1 for activating, -1 or 2 for inhibiting")
    }
  }
  if(length(state) == 1){
    if(state == "activate" || state == "activation" || state == "active" || state == "positive"){
      state <- "activating"
    }
    if(state == "inhibit" || state == "inhibition" || state == "inhibitory" || state == "negative"){
      state <- "inhibiting"
    }
    if(state == "activating"){
      if(is.null(col.arrow)) col.arrow <- par("fg")
      arrows(x0 = (1-arrow_clip) * Xn[match(as.character(es$V1), names(vs))] + arrow_clip * Xn[match(as.character(es$V2), names(vs))], y0 = (1-arrow_clip) * Yn[match(as.character(es$V1), names(vs))] + arrow_clip * Yn[match(as.character(es$V2), names(vs))],  x1 = (1-arrow_clip) * Xn[match(as.character(es$V2), names(vs))] + arrow_clip * Xn[match(as.character(es$V1), names(vs))],  y1 = (1-arrow_clip) * Yn[match(as.character(es$V2), names(vs))]  + arrow_clip * Yn[match(as.character(es$V1), names(vs))], lwd=cex.arrow, col=col.arrow, length=0.15)
    } else if (state =="inhibiting"){
      if(is.null(col.arrow)) col.arrow <- "red"
      arrows(x0 = (1-arrow_clip) * Xn[match(as.character(es$V1), names(vs))] + arrow_clip * Xn[match(as.character(es$V2), names(vs))], y0 = (1-arrow_clip) * Yn[match(as.character(es$V1), names(vs))] + arrow_clip * Yn[match(as.character(es$V2), names(vs))],  x1 = (1-arrow_clip) * Xn[match(as.character(es$V2), names(vs))] + arrow_clip * Xn[match(as.character(es$V1), names(vs))],  y1 = (1-arrow_clip) * Yn[match(as.character(es$V2), names(vs))]  + arrow_clip * Yn[match(as.character(es$V1), names(vs))], lwd=cex.arrow, col=col.arrow, length=0.1, angle=90)
    } else{
      warning("Please give state as a scalar or vector of length(E(graph)): input must be 'activating', 'inhibiting' or an integer")
      stop()
    }
  } else{
    if(length(col.arrow)==1) col.arrow <- rep(col.arrow, Ne)
    if(length(cex.arrow)==1) cex.arrow <- rep(cex.arrow, Ne)
    for(i in 1:Ne){
      v0 <- es[i, ]$V1
      v1 <- es[i, ]$V2
      if(state[i] == "activate" || state[i] == "activation" || state[i] == "active" || state[i] == "positive"){
        state[i] <- "activating"
      }
      if(state[i] == "inhibit" || state[i] == "inhibition" || state[i] == "inhibitory" || state[i] == "negative"){
        state[i] <- "inhibiting"
      }
      if(state[i] == "activating"){
        if(is.null(col.arrow[i])) col.arrow[i] <- par("fg")
        arrows(x0 = (1-arrow_clip) * Xn[match(as.character(v0), names(vs))] + arrow_clip * Xn[match(as.character(v1), names(vs))], y0 = (1-arrow_clip) * Yn[match(as.character(v0), names(vs))] + arrow_clip * Yn[match(as.character(v1), names(vs))],  x1 = (1-arrow_clip) * Xn[match(as.character(v1), names(vs))] + arrow_clip * Xn[match(as.character(v0), names(vs))],  y1 = (1-arrow_clip) * Yn[match(as.character(v1), names(vs))]  + arrow_clip * Yn[match(as.character(v0), names(vs))], lwd=cex.arrow[i], col=col.arrow[i], length=0.15)
      } else if (state[i] =="inhibiting"){
        if(is.null(col.arrow[i])) col.arrow[i] <- "red"
        arrows(x0 = (1-arrow_clip) * Xn[match(as.character(v0), names(vs))] + arrow_clip * Xn[match(as.character(v1), names(vs))], y0 = (1-arrow_clip) * Yn[match(as.character(v0), names(vs))] + arrow_clip * Yn[match(as.character(v1), names(vs))],  x1 = (1-arrow_clip) * Xn[match(as.character(v1), names(vs))] + arrow_clip * Xn[match(as.character(v0), names(vs))],  y1 = (1-arrow_clip) * Yn[match(as.character(v1), names(vs))]  + arrow_clip * Yn[match(as.character(v0), names(vs))], lwd=cex.arrow[i], col=col.arrow[i], length=0.1, angle=90)
      } else{
        warning("please give state as a scalar or vector of length(E(graph))")
        stop()
      }
    }
  }
  if(is.null(labels)) labels <- names(vs)
  points(Xn, Yn, xaxt="n", yaxt="n", xlab=xlab, ylab=ylab, cex = 2 * cex.node, pch=21, col=border.node, bg=fill.node, main=main, sub=sub, cex.main=cex.main)
  text(Xn, Yn, labels=labels, cex = cex.label*cex.node, col=col.label)
}

plot.directed <- plot_directed
```
