## Supplementary material for "graphsim: An R package for simulating gene expression data from graph structures of biological pathways": graphsim 1.0.0 R package: raf_mapk_graph.R

```
#' #' RAF/MAP kinase cascade 
#'
#' Reactome pathway R-HSA-5673001 for the interactions in the RAF/MAP kinase cascade
#' 
#' @format A graph object of 17 vertices and 121 edges:
#' \describe{
#'   \item{V}{gene symbol (human)}
#'   \item{E}{directed relationship for pathway}
#' }
#' @source PathwayCommons \url{https://reactome.org/content/detail/R-HSA-5673001}
"RAF_MAP_graph"
```
