## Supplementary material for "graphsim: An R package for simulating gene expression data from graph structures of biological pathways": graphsim 1.0.0 R package: sigma.R

```
##' @name make_sigma
##' @rdname make_sigma
##'
##' @title Generate Sigma (\eqn{\Sigma}) Matrix
##'
##' @description Compute the Sigma (\eqn{\Sigma}) matrix from an \code{\link[igraph]{igraph}} structure 
##' or pre-computed matrix. These are compatible with \code{\link[mvtnorm]{rmvnorm}} and
##'  \code{\link[graphsim]{generate_expression}}.
##' By default data is generated with a mean of 0 and standard deviation of 1 for 
##' each gene (with correlations between derived from the graph structure).
##' Thus where the Sigma (\eqn{\Sigma}) matrix has diagonals of 1 (for the variance of each gene)
##' then the symmetric non-diagonal terms (for covariance) determine the correlations
##' between each gene in the output from \code{\link[graphsim]{generate_expression}}.
##'
##' @param mat precomputed adjacency, laplacian, commonlink, or scaled distance matrix.
##' @param graph An \code{\link[igraph]{igraph}} object. May be directed or weighted.
##' @param state numeric vector. Vector of length E(graph). Sign used to calculate 
##' state matrix, may be an integer state or inferred directly from expected correlations
##' for each edge. May be applied a scalar across all edges or as a vector for each edge
##' respectively. May also be entered as text for "activating" or "inhibiting" or as
##' integers for activating (0,1) or inhibiting (-1,2). Compatible with inputs for 
##' \code{\link[graphsim]{plot_directed}}. Also takes a pre-computed state matrix from
##' \code{\link[graphsim]{make_state_matrix}} if applied to the same graph multiple times.
##' @param cor numeric. Simulated maximum correlation/covariance of two adjacent nodes.
##' Default to 0.8.
##' @param sd	 standard deviations of each gene. Defaults to 1. May be entered as a scalar
##' applying to all genes or a vector with a separate value for each. 
##' @param directed logical. Whether directed information is passed to the distance matrix.
##' @param comm logical whether a common link matrix is used to compute sigma.
##' Defaults to FALSE (adjacency matrix).
##' @param laplacian logical whether a Laplacian matrix is used to compute sigma.
##' Defaults to FALSE (adjacency matrix).
##' @param absolute logical. Whether distances are scaled as the absolute difference from
##' the diameter (maximum possible). Defaults to TRUE. The alternative is to calculate a
##' relative difference from the diameter for a geometric decay in distance.
##' @keywords graph network igraph mvtnorm
##' @importFrom igraph as_adjacency_matrix
##' @importFrom igraph graph_from_adjacency_matrix set_edge_attr V E
##' @examples 
##' 
##' # construct a synthetic graph module
##' library("igraph")
##' graph_test_edges <- rbind(c("A", "B"), c("B", "C"), c("B", "D"))
##' graph_test <- graph.edgelist(graph_test_edges, directed = TRUE)
##' # compute sigma (\eqn{\Sigma}) matrix for toy example
##' sigma_matrix <- make_sigma_mat_graph(graph_test, cor = 0.8)
##' sigma_matrix
##' 
##' # compute sigma (\eqn{\Sigma}) matrix  from adjacency matrix for toy example
##' adjacency_matrix <- make_adjmatrix_graph(graph_test)
##' sigma_matrix <- make_sigma_mat_adjmat(adjacency_matrix, cor = 0.8)
##' sigma_matrix
##' 
##' # compute sigma (\eqn{\Sigma}) matrix from shared edges for toy example
##' common_link_matrix <- make_commonlink_graph(graph_test)
##' sigma_matrix <- make_sigma_mat_comm(common_link_matrix, cor = 0.8)
##' sigma_matrix
##' 
##' # compute sigma (\eqn{\Sigma}) matrix from Laplacian for toy example
##' laplacian_matrix <- make_laplacian_graph(graph_test)
##' sigma_matrix <- make_sigma_mat_laplacian(laplacian_matrix, cor = 0.8)
##' sigma_matrix
##' 
##' # compute sigma (\eqn{\Sigma}) matrix from distance matrix for toy example
##' distance_matrix <- make_distance_graph(graph_test, absolute = FALSE)
##' sigma_matrix <- make_sigma_mat_dist_adjmat(distance_matrix, cor = 0.8)
##' sigma_matrix
##' 
##' # compute sigma (\eqn{\Sigma}) matrix from geometric distance directly from toy example graph
##' sigma_matrix <- make_sigma_mat_dist_graph(graph_test, cor = 0.8)
##' sigma_matrix
##' 
##' # compute sigma (\eqn{\Sigma}) matrix from absolute distance directly from toy example graph
##' sigma_matrix <- make_sigma_mat_dist_graph(graph_test, cor = 0.8, absolute = TRUE)
##' sigma_matrix
##' 
##' # compute sigma (\eqn{\Sigma}) matrix from geometric distance with sd = 2
##' sigma_matrix <- make_sigma_mat_dist_graph(graph_test, cor = 0.8, sd = 2)
##' sigma_matrix
##' 
##' # construct a synthetic graph network
##' graph_structure_edges <- rbind(c("A", "C"), c("B", "C"), c("C", "D"), c("D", "E"),
##'                                c("D", "F"), c("F", "G"), c("F", "I"), c("H", "I"))
##' graph_structure <- graph.edgelist(graph_structure_edges, directed = TRUE)
##' 
##' # compute sigma (\eqn{\Sigma}) matrix from geometric distance directly from synthetic graph network
##' sigma_matrix_graph_structure <- make_sigma_mat_dist_graph(graph_structure,
##'                                                           cor = 0.8, absolute = FALSE)
##' sigma_matrix_graph_structure
##' # visualise matrix
##' library("gplots")
##' heatmap.2(sigma_matrix_graph_structure, scale = "none", trace = "none",
##'                      col = colorpanel(50, "white", "red"))
##' 
##' # compute sigma (\eqn{\Sigma}) matrix from geometric distance directly from
##' # synthetic graph network with inhibitions
##' edge_state <- c(1, 1, -1, 1, 1, 1, 1, -1)
##' # pass edge state as a parameter
##' sigma_matrix_graph_structure_inhib <- make_sigma_mat_dist_graph(graph_structure, 
##'                                                                 state = edge_state,
##'                                                                 cor = 0.8,
##'                                                                 absolute = FALSE)
##' sigma_matrix_graph_structure_inhib
##' # visualise matrix
##' library("gplots")
##' heatmap.2(sigma_matrix_graph_structure_inhib, scale = "none", trace = "none",
##'           col = colorpanel(50, "blue", "white", "red"))
##' 
##' # compute sigma (\eqn{\Sigma}) matrix from geometric distance directly from 
##' # synthetic graph network with inhibitions
##' E(graph_structure)$state <-  c(1, 1, -1, 1, 1, 1, 1, -1)
##' # pass edge state as a graph attribute
##' sigma_matrix_graph_structure_inhib <- make_sigma_mat_dist_graph(graph_structure,
##'                                                                 cor = 0.8,
##'                                                                 absolute = FALSE)
##' sigma_matrix_graph_structure_inhib
##' # visualise matrix
##' library("gplots")
##' heatmap.2(sigma_matrix_graph_structure_inhib, scale = "none", trace = "none",
##'           col = colorpanel(50, "blue", "white", "red"))
##' 
##' # import graph from package for reactome pathway
##' # TGF-\eqn{\Beta} receptor signaling activates SMADs (R-HSA-2173789)
##' TGFBeta_Smad_graph <- identity(TGFBeta_Smad_graph)
##' 
##' # compute sigma (\eqn{\Sigma}) matrix from geometric distance directly from TGF-\eqn{\Beta} pathway
##' TFGBeta_Smad_state <- E(TGFBeta_Smad_graph)$state
##' table(TFGBeta_Smad_state)
##' # states are edge attributes
##'  sigma_matrix_TFGBeta_Smad_inhib <- make_sigma_mat_dist_graph(TGFBeta_Smad_graph,
##'                                                               cor = 0.8,
##'                                                               absolute = FALSE)
##' # visualise matrix
##' library("gplots")
##' heatmap.2(sigma_matrix_TFGBeta_Smad_inhib, scale = "none", trace = "none",
##'           col = colorpanel(50, "blue", "white", "red"))
##' 
##' # compute sigma (\eqn{\Sigma}) matrix from geometric distance directly from TGF-\eqn{\Beta} pathway
##' TGFBeta_Smad_graph <- remove.edge.attribute(TGFBeta_Smad_graph, "state")
##' # compute with states removed (all negative)
##' sigma_matrix_TFGBeta_Smad <- make_sigma_mat_dist_graph(TGFBeta_Smad_graph,
##'                                                        state = -1,
##'                                                        cor = 0.8,
##'                                                        absolute = FALSE)
##' # visualise matrix
##' library("gplots")
##' heatmap.2(sigma_matrix_TFGBeta_Smad, scale = "none", trace = "none",
##'           col = colorpanel(50, "white", "red"))
##' # compute with states removed (all positive)
##' sigma_matrix_TFGBeta_Smad <- make_sigma_mat_dist_graph(TGFBeta_Smad_graph,
##'                                                        state = 1,
##'                                                        cor = 0.8,
##'                                                        absolute = FALSE)
##' # visualise matrix
##' library("gplots")
##' heatmap.2(sigma_matrix_TFGBeta_Smad, scale = "none", trace = "none",
##'           col = colorpanel(50, "white", "red"))
##' 
##' #restore edge attributes
##' TGFBeta_Smad_graph <- set_edge_attr(TGFBeta_Smad_graph, "state",
##'                                     value = TFGBeta_Smad_state)
##' TFGBeta_Smad_state <- E(TGFBeta_Smad_graph)$state
##' # states are edge attributes
##'  sigma_matrix_TFGBeta_Smad_inhib <- make_sigma_mat_dist_graph(TGFBeta_Smad_graph,
##'                                                               cor = 0.8,
##'                                                               absolute = FALSE)
##' # visualise matrix
##' library("gplots")
##' heatmap.2(sigma_matrix_TFGBeta_Smad_inhib, scale = "none", trace = "none",
##'           col = colorpanel(50, "blue", "white", "red"))
##' 
##' @return a numeric covariance matrix of values in the range [-1, 1]
##' @export
make_sigma_mat_adjmat <- function(mat, state = NULL, cor = 0.8, sd = 1){
  sig <- ifelse(mat > 0, cor, 0)
  diag(sig) <- 1
  if(!is.null(state)){
    #pass state parameters sign of sigma
    graph <- graph_from_adjacency_matrix(mat, mode = "undirected")
    if(length(state) == 1) state <- rep(state, length(E(graph)))
    graph <- set_edge_attr(graph, "state", value = state)
    state_mat <- make_state_matrix(graph, state)
    sig <- sig * sign(state_mat)
  }
  if(sd[1] != 1 || length(sd) != 1){
    if(!(is.vector(sd)) || length(sd) == 1 ) sd <- rep(sd,ncol(mat))
    sig <- t(sd * t(sig)) * sd
  }
  rownames(sig) <- rownames(mat)
  colnames(sig) <- colnames(mat)
  return(sig)
}

##' @rdname make_sigma
##' @export
make_sigma_mat_comm <- function(mat, state = NULL, cor = 0.8, sd = 1){
  mat <- abs(mat)
  mat <- apply(mat, 1, function(x) x/max(x))
  mat <- apply(mat, 2, function(x) x/max(x))
  sig <- ifelse(mat>0, cor*mat/max(mat), 0)
  diag(sig) <- 1
  if(!is.null(state)){
    #pass state parameters sign of sigma
    graph <- graph_from_adjacency_matrix(mat, mode = "undirected")
    if(length(state) == 1) state <- rep(state, length(E(graph)))
    graph <- set_edge_attr(graph, "state", value = state)
    state_mat <- make_state_matrix(graph, state)
    sig <- sig * sign(state_mat)
  }
  if(sd[1] != 1 || length(sd) != 1){
    if(!(is.vector(sd)) || length(sd) == 1 ) sd <- rep(sd,ncol(mat))
    sig <- t(sd * t(sig)) * sd
  }
  rownames(sig) <- rownames(mat)
  colnames(sig) <- colnames(mat)
  return(sig)
}

##' @rdname make_sigma
##' @export
make_sigma_mat_laplacian <- function(mat, state = NULL, cor = 0.8, sd = 1){
  mat <- abs(mat)
  diag(mat)[diag(mat) == 0] <- 1
  mat <- apply(mat, 1, function(x) x/max(x))
  mat <- apply(mat, 2, function(x) x/max(x))
  sig <- ifelse(mat>0, cor*mat/max(mat), 0)
  diag(sig) <- 1
  if(!is.null(state)){
    #pass state parameters sign of sigma
    graph <- graph_from_adjacency_matrix(mat, mode = "undirected")
    if(length(state) == 1) state <- rep(state, length(E(graph)))
    graph <- set_edge_attr(graph, "state", value = state)
    state_mat <- make_state_matrix(graph, state)
    sig <- sig * sign(state_mat)
  }
  if(sd[1] != 1 || length(sd) != 1){
    if(!(is.vector(sd)) || length(sd) == 1 ) sd <- rep(sd,ncol(mat))
    sig <- t(sd * t(sig)) * sd
  }
  rownames(sig) <- rownames(mat)
  colnames(sig) <- colnames(mat)
  return(sig)
}

##' @rdname make_sigma
##' @export
##' 
make_sigma_mat_graph <- function(graph, state = NULL, cor = 0.8, sd = 1, comm = FALSE, laplacian = FALSE, directed = FALSE){
  if(!is.null(get.edge.attribute(graph, "state"))){
    state <- get.edge.attribute(graph, "state")
  } else {
    # add default state if not specified
    if(is.null(state)){
      state <- "activating"
    }
  }
  if(comm && laplacian){
    warning("Error: only one of commonlink or laplacian can be used")
    stop()
  }
  if(!laplacian) mat <- make_adjmatrix_graph(graph, directed = directed)
  if(comm){
    mat <- make_commonlink_adjmat(mat)
    diag(mat) <-  max(mat) + 1
  }
  if(laplacian){
    mat <- make_laplacian_graph(graph, directed = directed)
    mat <- abs(mat)
    diag(mat)[diag(mat) == 0] <- 1
    mat <- apply(mat, 1, function(x) x/max(x))
    mat <- apply(mat, 2, function(x) x/max(x))
  } else{
    diag(mat) <- 1
  }
  sig <- ifelse(mat>0, cor*mat/max(mat), 0)
  diag(sig) <- 1
  if(!is.null(state)){
    #pass state parameters sign of sigma
    state_mat <- make_state_matrix(graph, state)
    sig <- sig * sign(state_mat)
  }
  if(sd[1] != 1 || length(sd) != 1){
    if(!(is.vector(sd)) || length(sd) == 1 ) sd <- rep(sd,length(V(graph)))
    sig <- t(sd * t(sig)) * sd
  }
  rownames(sig) <- rownames(mat)
  colnames(sig) <- colnames(mat)
  return(sig)
}

##' @rdname make_sigma
##' @export
make_sigma_mat_dist_adjmat <- function(mat, state = NULL, cor = 0.8, sd = 1, absolute = FALSE){
  if(!(all(diag(mat) == 1))) stop("distance matrix must have diagonal of zero")
  if(!(max(mat[mat != 1]) > 0) || !(max(mat[mat!=1]) <= 1)) stop("distance matrix expected, not adjacency matrix")
  sig <- mat/max(mat[mat != 1]) * cor
  sig <- ifelse(sig > 0, sig, 0)
  diag(sig) <- 1
  if(!is.null(state)){
    #adjacency matrix from distance
    adjmat <- ifelse(mat == max(mat[mat != 1]), 1, 0)
    graph <- graph_from_adjacency_matrix(adjmat, mode = "undirected")
    graph <- as.directed(graph, mode = "arbitrary")
    #pass state parameters sign of sigma
    state_mat <- make_state_matrix(graph, state)
    sig <- sig * sign(state_mat)
  }
  if(sd[1] != 1 || length(sd) != 1){
    if(!(is.vector(sd)) || length(sd) == 1 ) sd <- rep(sd,ncol(mat))
    sig <- t(sd * t(sig)) * sd
  }
  rownames(sig) <- rownames(mat)
  colnames(sig) <- colnames(mat)
  return(sig)
}


##' @rdname make_sigma
##' @export
make_sigma_mat_dist_graph <- function(graph, state = NULL, cor = 0.8, sd = 1, absolute = FALSE){
  if(!is.null(get.edge.attribute(graph, "state"))){
    state <- get.edge.attribute(graph, "state")
  } else {
    # add default state if not specified
    if(is.null(state)){
      state <- "activating"
    }
  }
  mat <- make_distance_graph(graph, absolute = absolute)
  if(!(all(diag(mat) == 1))) stop("distance matrix must have diagonal of zero")
  if(!(max(mat[mat != 1]) > 0) || !(max(mat[mat!=1]) <= 1)) stop("distance matrix expected, not adjacency matrix")
  sig <- mat/max(mat[mat != 1]) * cor
  sig <- ifelse(sig > 0, sig, 0)
  diag(sig) <- 1
  #derive states directly from graph if available
  state_mat <- make_state_matrix(graph, state)
  sig <- sig * sign(state_mat)
  if(sd[1] != 1 || length(sd) != 1){
    if(!(is.vector(sd)) || length(sd) == 1 ) sd <- rep(sd,length(V(graph)))
    sig <- t(sd * t(sig)) * sd
  }
  rownames(sig) <- rownames(mat)
  colnames(sig) <- colnames(mat)
  return(sig)
  return(sig)
}
```
