## Supplementary material for "graphsim: An R package for simulating gene expression data from graph structures of biological pathways": graphsim 1.0.0 R package: state.R

```
##' @name make_state_matrix
##' @rdname make_state_matrix
##'
##' @title Make State Matrix
##' 
##' @description Functions to compute the matrix of states (1 for activating and -1 for inhibiting) 
##' for link signed correlations, from a vector of edge states to a signed adjacency matrix for use
##' in \code{\link[graphsim]{generate_expression}}. This resolves edge states to determine the sign
##' of all correlations between nodes in a network. These are computed interally for sigma matrices
##' as required.
##' @param graph An \code{\link[igraph]{igraph}} object. May be directed or weighted as long as
##' a shortest path can be computed.
##' @param state numeric vector. Vector of length E(graph). Sign used to calculate state matrix,
##' may be an integer state or inferred directly from expected correlations for each edge. May be
##' applied a scalar across all edges or as a vector for each edge respectively. May also be
##' entered as text for "activating" or "inhibiting" or as integers for activating (0,1) or
##' inhibiting (-1,2). Compatible with inputs for \code{\link[graphsim]{plot_directed}}. Vector 
##' input is supported either directly calling the function with a value for each edge in 
##'  \code{E(graph)} or as an edge "attribute" in the igraph object (using 
##'  \code{E(g)$state <- states}).
##' @keywords graph network igraph mvtnorm simulation
##' @importFrom igraph graph.edgelist get.edge.attribute get.edgelist
##' @import igraph
##' @examples 
##' 
##' # construct a synthetic graph module
##' library("igraph")
##' graph_test_edges <- rbind(c("A", "B"), c("B", "C"), c("B", "D"))
##' graph_test <- graph.edgelist(graph_test_edges, directed = TRUE)
##' 
##'  # compute state matrix for toy example
##' state_matrix <- make_state_matrix(graph_test)
##' 
##' # construct a synthetic graph network
##' graph_structure_edges <- rbind(c("A", "C"), c("B", "C"), c("C", "D"), c("D", "E"),
##'                                c("D", "F"), c("F", "G"), c("F", "I"), c("H", "I"))
##' graph_structure <- graph.edgelist(graph_structure_edges, directed = TRUE)
##' 
##' # compute state matrix for toy network
##' graph_structure_state_matrix <- make_state_matrix(graph_structure)
##' graph_structure_state_matrix
##' 
##' # compute state matrix for toy network with inhibitions
##' edge_state <- c(1, 1, -1, 1, 1, 1, 1, -1)
##' # edge states are a variable
##' graph_structure_state_matrix <- make_state_matrix(graph_structure, state = edge_state)
##' graph_structure_state_matrix
##' 
##' # compute state matrix for toy network with inhibitions
##' E(graph_structure)$state <- c(1, 1, -1, 1, 1, 1, 1, -1)
##' # edge states are a graph attribute
##' graph_structure_state_matrix <- make_state_matrix(graph_structure)
##' graph_structure_state_matrix
##' 
##' library("igraph")
##' graph_test_edges <- rbind(c("A", "B"), c("B", "C"), c("B", "D"))
##' graph_test <- graph.edgelist(graph_test_edges, directed = TRUE)
##' state_matrix <- make_state_matrix(graph_test)
##' 
##' # import graph from package for reactome pathway
##' # TGF-\eqn{\Beta} receptor signaling activates SMADs (R-HSA-2173789)
##' TGFBeta_Smad_graph <- identity(TGFBeta_Smad_graph)
##' 
##' # compute sigma (\eqn{\Sigma}) matrix from geometric distance directly from TGF-\eqn{\Beta} pathway
##' TFGBeta_Smad_state <- E(TGFBeta_Smad_graph)$state
##' table(TFGBeta_Smad_state)
##' # states are edge attributes
##' state_matrix_TFGBeta_Smad <- make_state_matrix(TGFBeta_Smad_graph)
##' # visualise matrix
##' library("gplots")
##' heatmap.2(state_matrix_TFGBeta_Smad , scale = "none", trace = "none",
##'           dendrogram = "none", Rowv = FALSE, Colv = FALSE,
##'           col = colorpanel(50, "blue", "white", "red"))
##' 
##' # compare the states to the sign of expected correlations in the sigma matrix
##' sigma_matrix_TFGBeta_Smad_inhib <- make_sigma_mat_dist_graph(TGFBeta_Smad_graph,
##'                                                              cor = 0.8,
##'                                                              absolute = FALSE)
##' # visualise matrix
##' heatmap.2(sigma_matrix_TFGBeta_Smad_inhib,
##'           scale = "none", trace = "none",
##'           dendrogram = "none", Rowv = FALSE, Colv = FALSE,
##'           col = colorpanel(50, "blue", "white", "red"))
##' 
##' # compare the states to the sign of final correlations in the simulated matrix
##' TFGBeta_Smad_data <- generate_expression(100, TGFBeta_Smad_graph, cor = 0.8)
##' heatmap.2(cor(t(TFGBeta_Smad_data)), scale = "none", trace = "none",
##'           dendrogram = "none", Rowv = FALSE, Colv = FALSE,
##'           col = colorpanel(50, "blue", "white", "red"))
##' 
##' 
##' @return An integer matrix indicating the resolved state
##' (activating or inhibiting for each edge or path between nodes)
##' @export
make_state_matrix <- function(graph, state = NULL){
  if(!is.null(get.edge.attribute(graph, "state"))){
    state <- get.edge.attribute(graph, "state")
  } else {
    # add default state if not specified
    if(is.null(state)){
      state <- "activating"
    }
  }
  if(is.null(get.edge.attribute(graph, "state"))){
    E(graph)$state <-  state
  }
  # this could also be done with igraph::simplify(graph, remove.multiple = TRUE, remove.loops = TRUE, edge.attr.comb = igraph_opt("edge.attr.comb"))
  ## to do: migrate to this
  graph <- as.undirected(graph, mode = "collapse", edge.attr.comb = function(x) ifelse(any(x %in% list(-1, 2, "inhibiting", "inhibition")), -1, 1))
  # remove duplicate edges (and corresponding states)
  graph <- as.directed(graph, mode = "arbitrary")
  state <- get.edge.attribute(graph, "state")
  if(length(state) == 1) state <- rep(state, length(E(graph)))
  state[state == 2] <- -1
  state[state == 1 | state == 0] <- 1
  if(is.numeric(state)) state[!(state %in% -1:2)] <- sign(state[!(state %in% -1:2)])
  if(is.character(state)){
    state <- as.list(state)
    state[grep("activating", state)] <- 1
    state[grep("activation", state)] <- 1
    state[grep("activate", state)] <- 1
    state[grep("active", state)] <- 1
    state[grep("positive", state)] <- 1
    state[grep("inhibiting", state)] <- -1
    state[grep("inhibition", state)] <- -1
    state[grep("inhibitory", state)] <- -1
    state[grep("inhibit", state)] <- -1
    state[grep("negative", state)] <- -1
    if(is.character(state)){
      warning("Please give state as a scalar or vector of length(E(graph)): input must be 'activating', 'inhibiting' or an integer")
    }
    state <- unlist(as.numeric(state))
  }
  if(!all(state %in% -1:2)){
    state <- sign(state) # coerce to vector or 1 and -1 if not already
    warning("State inferred from non-integer weighted edges: Please give numeric states as integers: 0 or 1 for activating, -1 or 2 for inhibiting")
  }
  #define starting states
  state_mat <- matrix(1, length(V(graph)), length(V(graph)))
  rownames(state_mat) <- colnames(state_mat) <- names(V(graph))
  # check for inhibiting edges
  if(all(state == 1)){
    #return without resolving for activations
    return(state_mat)
  } else if (length(V(graph)) == 1){
    state_mat <- 1
    return(state_mat)
  } else if (length(V(graph)) == 2){
    if(length(state) == 1){
      state_mat[row(state_mat) != col(state_mat)] <- state # one edge
      return(state_mat)
    } else{
      warning("only one edge state expected")
    }
  } else {
    #resolve inhibitions downstream
    # define shortest paths to all possible nodes
    #check if connected or use connected subgraphs
    compute_paths <- function(graph){
      if(is.connected(graph)){
        #define paths from 1st node
        paths <- shortest_paths(as.undirected(graph), V(graph)$name[1])$vpath
      } else {
        subgraphs <- decompose(graph)
        nodes <- sapply(subgraphs, function(subgraph) V(subgraph)$name[1])
        paths <- as.list(rep(NA, length(V(graph))))
        jj <- 0
        for(ii in 1:length(nodes)){
          subpaths <- shortest_paths(as.undirected(subgraphs[[ii]]), V(subgraphs[[ii]])$name[1])$vpath
          paths[jj+1:length(subpaths)] <- subpaths
          jj <- jj + length(subpaths)
        }
      }
      paths
    }
    paths <- compute_paths(graph)
    #check for cycles
    is.cyclic <- function(paths){
      cylic_paths <- sapply(paths, function(path){
        if(length(path) <= 1){
          FALSE
        } else if (path[1] == path[length(path)]){
          TRUE
        } else {
          FALSE
        }
      })
      any(cylic_paths)
    }
    if(is.cyclic(paths)){
      warning("Graph contains a cycle, computing minimal spanning tree. This may result in unresolved inhibitions.")
      tree <- minimum.spanning.tree(graph)
      paths <- compute_paths(tree)
      if(is.cyclic(paths)){
        warning("Graph contains cycles and cannot compute a minimal spanning tree. This will result in unresolved inhibitions.")
        stop()
      }
    }
    #sort paths into longest to shortest
    paths <- paths[order(sapply(paths, length), decreasing = TRUE)]
    #remove subpaths
    for(ii in 2:length(paths)){
      remove <- FALSE
      for(jj in 1:(ii-1)){
        # test if all nodes in path in a larger path above
        if(all(names(paths[[ii]]) %in% names(paths[[jj]]))){
          remove <- TRUE
        }
      }
      if(remove){
        paths[[ii]] <- NA
      }
    }
    na.omit.list <- function(y) { return(y[!sapply(y, function(x) all(is.na(x)))]) }
    paths <- na.omit.list(paths)
    # check inhibition state through graph paths
    edges <- as.matrix(get.edgelist(graph))
    for(ii in 1:length(paths)){
      if(length(paths[[ii]]) > 1){
        state_path <- c(1, rep(NA, length(paths[[ii]])-1))
        for(jj in 2:length(paths[[ii]])){
          #find edges
          kk <- which(
          edges[,1] == names(paths[[ii]])[jj-1] & edges[,2] == names(paths[[ii]])[jj] |
          edges[,2] == names(paths[[ii]])[jj-1] & edges[,1] == names(paths[[ii]])[jj]
          )
          #check for multiple edges
          if(length(kk) > 1){
            #check for same edges
            if(!all(apply(edges[kk,], 2, function(x) all(x == x[1])))){
              #reverse edge if no match
              edges[kk,][edges[kk[1],1] != edges[kk,1],] <- edges[kk,][edges[kk[1],1] != edges[kk,1],2:1]
            }
              #check is still no match
            if(all(apply(edges[kk,], 2, function(x) all(x == x[1])))){
              #check for same state
              if(all(state[kk] == state[kk][1])){
                state[kk] <- state[kk][1]
                kk <- kk[1]
              } else {
                warning(paste("Conflicting state for redundant edges", edges[kk[1],1], edges[kk[1],2], "with state", state[kk]))
                Mode <- function(x) unique(x)[which.max(tabulate(match(x, unique(x))))][1]
                state[kk] <- Mode(state[kk])
                kk <- kk[1]
              }
            }
          }
          #state for edges
          if(length(state_path[jj]) == length(state[kk])){
            state_path[jj] <- state[kk]
          } else {
            print(ii)
            print(jj)
            stop("State not compatible with graph paths")
          }
        }
        #cumulative product changes state along a path (*-1)
        state_path <- cumprod(state_path)
        #skip if activating path
        if(any(state_path != 1)){
          #cross-product produces a matrix form
          state_cross <- state_path %*% t(state_path)
          #add to state_matrix (all adjusted to start of path)
          state_mat[names(paths[[ii]]), names(paths[[ii]])] <- state_cross
        }
      }
    }
    #check for negative and positive links not in paths (against the 1st node)
    neg_nodes <- colnames(state_mat)[which(state_mat[1,] == -1)]
    pos_nodes <- colnames(state_mat)[2:ncol(state_mat)][which(state_mat[1,2:ncol(state_mat)] == 1)] # can assume at least 3 nodes due to checks above
    #check for if not computed in shorted paths already
    ##paths <- compute_paths(tree) # not minimal spanning tree
    for(aa in neg_nodes){
      for(bb in pos_nodes){
        aa_in_path <- sapply(paths, function(path) aa %in% names(path))
        bb_in_path <- sapply(paths, function(path) bb %in% names(path))
        #check if in ANY same path
        if(!any(aa_in_path & bb_in_path)){
          #if not invert
          state_mat[aa, bb] <- state_mat[bb, aa] <- -1
        }
      }
    }
    # check if same sign (to do: allow disconnected)
    for(aa in pos_nodes){
      for(bb in pos_nodes){
        aa_in_path <- sapply(paths, function(path) aa %in% names(path))
        bb_in_path <- sapply(paths, function(path) bb %in% names(path))
        #check if in ANY same path
        if(!any(aa_in_path & bb_in_path)){
          #if not invert
          state_mat[aa, bb] <- state_mat[bb, aa] <- 1
        }
      }
    }
    #ensure symmetric matrix
    state_mat[t(state_mat) == -1] <- -1
    return(state_mat)
  }
}
```
