## Supplementary material for "graphsim: An R package for simulating gene expression data from graph structures of biological pathways": graphsim 1.0.0 R package: tfgb_smad_graph.R

```
#' TGF-\eqn{\beta} receptor signaling activates SMADs
#'
#' Reactome pathway R-HSA-2173789 for the interactions in the TGF-\eqn{\beta} receptor signaling activates SMADs
#' 
#' @format A graph object of 32 vertices and 173 edges:
#' \describe{
#'   \item{V}{gene symbol (human)}
#'   \item{E}{directed relationship for pathway}
#'   \item{state}{type of relationship (activating or inhibiting) as edge attribute}
#' }
#' @source PathwayCommons \url{https://reactome.org/content/detail/R-HSA-2173789}
"TGFBeta_Smad_graph"
```
