## Supplementary material for "graphsim: An R package for simulating gene expression data from graph structures of biological pathways": graphsim 1.0.0 R package: README.md

[![Travis-CI Build Status](https://travis-ci.org/TomKellyGenetics/graphsim.svg?branch=master)](https://travis-ci.org/TomKellyGenetics/graphsim)
[![CircleCI build status](https://circleci.com/gh/TomKellyGenetics/graphsim.svg?style=svg)](https://circleci.com/gh/TomKellyGenetics/graphsim)
[![AppVeyor Build Status](https://ci.appveyor.com/api/projects/status/github/TomKellyGenetics/graphsim?branch=master&svg=true)](https://ci.appveyor.com/project/TomKellyGenetics/graphsim)
[![Project Status: Active – The project has reached a stable, usable state and is being actively developed.](http://www.repostatus.org/badges/latest/active.svg)](http://www.repostatus.org/#active)
[![codecov](https://codecov.io/gh/TomKellyGenetics/graphsim/branch/master/graph/badge.svg)](https://codecov.io/gh/TomKellyGenetics/graphsim)
[![GitHub Views](http://hits.dwyl.com/tomkellygenetics/graphsim.svg)](http://hits.dwyl.com/tomkellygenetics/graphsim)

[![CRAN_Status_Badge](http://www.r-pkg.org/badges/version/graphsim)](https://cran.r-project.org/package=graphsim)
[![Downloads](https://cranlogs.r-pkg.org/badges/graphsim)](https://CRAN.R-project.org/package=graphsim)
[![Total Downloads](https://cranlogs.r-pkg.org/badges/grand-total/graphsim?color=orange)](https://CRAN.R-project.org/package=graphsim)

[![DOI](https://zenodo.org/badge/67395230.svg)](https://zenodo.org/badge/latestdoi/67395230)
[![bioRxiv](https://img.shields.io/badge/biorXiv-10.1101%2F2020.03.02.972471-blue)](https://doi.org/10.1101/2020.03.02.972471)
[![status](https://joss.theoj.org/papers/96016c6a55d7f74bacebd187c6ededd6/status.svg)](https://joss.theoj.org/papers/96016c6a55d7f74bacebd187c6ededd6)
[![](https://img.shields.io/badge/Altmetric-71-blue.svg)](https://www.altmetric.com/details/77053356)

# graphsim

## Version 1.0.0

###  Simulate Expression Data from 'igraph' Networks 

This package provides functions to develop simulated continuous data 
(e.g., gene expression) from a sigma covariance matrix derived from a 
graph structure in 'igraph' objects. Intended to extend 'mvtnorm' to 
take 'igraph' structures rather than sigma matrices as input. This 
allows the use of simulated data that correctly accounts for pathway
relationships and correlations. Here we present a versatile statistical 
framework to simulate correlated gene expression data from biological 
pathways, by sampling from a multivariate normal distribution derived 
from a graph structure. This package allows the simulation of biological
pathways from a graph structure based on a statistical model of 
gene expression, such as simulation of expression profiles that
of log-transformed and normalised data from microarray and RNA-Seq data.
experiments. 

#### Motivation

Network analysis of molecular biological pathways is important
for insights into biology and medical genetics. 
Gene expression profiles capture the regulatory state of a cell
and can be used to analyse complex molecular states with genome-scale data.
Biological pathways are more than simply sets of genes involved in functions,
they are rich in information of relationships defined by pathway structure.

Methods to infer biological pathways and gene regulatory networks from gene
expression data can be tested on  simulated datasets using this framework. This also allows for
pathway structures to be considered as a confounding variable when 
simulating gene expression data to test the performance of genomics analyses.

This package enable the generation of simulated gene expression datasets
containing pathway relationships from a known underlying network.
These simulated datasets can be used to evaluate various bioinformatics
methodologies, including statistical and network inference procedures.

Network analysis techniques have an important role in understanding
of biological pathways and interpretation of genomics studies.
Modelling biological pathways allows the evaluation of gene
regulatory network inference techniques (which so far rely on
experimental validation or resampling). This technique also
enables modelling datasets with correlated pathway-structures
to assess whether other genomics analysis techniques perform
as expected with the background of complex pathways.


## Installation

To install the latest release from CRAN:

```R
install.packages("graphsim")
```

To install the stable release of this package from github:

```R
# install.packages("devtools")
devtools::install_github("TomKellyGenetics/graphsim", ref = "master")
```

To get the development version of this package from github:

```R
# install.packages("devtools")
devtools::install_github("TomKellyGenetics/graphsim", ref = "dev")
```

## Usage

Please see the vignettes for demonstrations of this package on examples of simple simulated networks and the reactome pathway TGF-&beta; receptor signaling activates SMADs (R-HSA-2173789). A manuscript with further details has been submitted for peer-review.

## Citation

To cite the graphsim package in publications use:

  S. Thomas Kelly and Michael A. Black (2020). graphsim: Simulate Expression data from iGraph networks 
  R package version 1.0.0. https://github.com/TomKellyGenetics/graphsim doi:10.5281/zenodo.1313986

A BibTeX entry for LaTeX users is

```
  @Manual{,
    title = {{graphsim}: Simulate Expression data from {iGraph} networks},
    author = {S. Thomas Kelly and Michael A. Black},
    year = {2020},
    note = {R package version 1.0.0},
    url = {https://github.com/TomKellyGenetics/graphsim},
    doi = {10.5281/zenodo.1313986},
  }
```

Please also acknowledge the manuscript describing use of this package once it is published. It is currently avaliable as a preprint.

```
@article {Kelly2020.03.02.972471,
	author = {Kelly, S Thomas and Black, Michael A},
	title = {graphsim: An R package for simulating gene expression data from graph structures of biological pathways},
	elocation-id = {2020.03.02.972471},
	year = {2020},
	doi = {10.1101/2020.03.02.972471},
	publisher = {Cold Spring Harbor Laboratory},
	URL = {https://www.biorxiv.org/content/early/2020/03/04/2020.03.02.972471},
	eprint = {https://www.biorxiv.org/content/early/2020/03/04/2020.03.02.972471.full.pdf},
	journal = {bioRxiv}
}
```

## Contributions and Bug Reports

Please submit [issues](https://github.com/TomKellyGenetics/graphsim/issues) on GitHub to report
problems or suggest features. [Pull requests](https://github.com/TomKellyGenetics/graphsim/pulls)
to the `dev` branch on GitHub are also welcome to add features or correct problems. Please see
the [contributor guide](CONTRIBUTING.md) for more details.
