## Supplementary material for "graphsim: An R package for simulating gene expression data from graph structures of biological pathways": graphsim 1.0.0 R package: test_commonlink.R

```
library("graphsim")
library("igraph")
context("Make Commonlink Matrix")

test_that("Generate common link matrix from adjacency matrix", {
  graph_test1_edges <- rbind(c("A", "B"), c("B", "C"), c("B", "D"))
  graph_test1 <- graph.edgelist(graph_test1_edges, directed = TRUE)
  adjacency_matrix1 <- make_adjmatrix_graph(graph_test1)
  common_link_matrix1 <- make_commonlink_adjmat(adjacency_matrix1)
  expect_equal(isSymmetric(common_link_matrix1), TRUE)
  expect_equal(diag(common_link_matrix1), degree(graph_test1))
  expect_equal(sum(diag(common_link_matrix1)), sum(degree(graph_test1)))
  expect_equal(nrow(common_link_matrix1), length(V(graph_test1)))
  expect_equal(ncol(common_link_matrix1), length(V(graph_test1)))
  expect_equal(sum(common_link_matrix1), length(E(graph_test1))*4)
  expect_true(all(adjacency_matrix1 == cbind(c(0, 1, 0, 0), c(1, 0, 1, 1), c(0, 1, 0, 0), c(0, 1, 0, 0))))
  expect_equal(all(is.matrix(common_link_matrix1)), TRUE)
  expect_true(all(common_link_matrix1 == cbind(c(1, 0, 1, 1), c(0, 3, 0, 0), c(1, 0, 1, 1), c(1, 0, 1, 1))))
})

test_that("Generate common link matrix from graph structure", {
  graph_test1_edges <- rbind(c("A", "B"), c("B", "C"), c("B", "D"))
  graph_test1 <- graph.edgelist(graph_test1_edges, directed = TRUE)
  common_link_matrix1 <- make_commonlink_graph(graph_test1)
  expect_equal(isSymmetric(common_link_matrix1), TRUE)
  expect_equal(diag(common_link_matrix1), degree(graph_test1))
  expect_equal(sum(diag(common_link_matrix1)), sum(degree(graph_test1)))
  expect_equal(nrow(common_link_matrix1), length(V(graph_test1)))
  expect_equal(ncol(common_link_matrix1), length(V(graph_test1)))
  expect_equal(sum(common_link_matrix1), length(E(graph_test1))*4)
  expect_equal(all(is.matrix(common_link_matrix1)), TRUE)
  expect_true(all(common_link_matrix1 == cbind(c(1, 0, 1, 1), c(0, 3, 0, 0), c(1, 0, 1, 1), c(1, 0, 1, 1))))
})
```
