## Supplementary material for "graphsim: An R package for simulating gene expression data from graph structures of biological pathways": graphsim 1.0.0 R package: test_generate.R

```
library("graphsim")
library("igraph")
context("Generate Expression Data")

test_that("Generate Expression matrix from adjacency matrix", {
  graph_test1_edges <- rbind(c("A", "B"), c("B", "C"), c("B", "D"))
  graph_test1 <- graph.edgelist(graph_test1_edges, directed = TRUE)
  adjacency_matrix1 <- make_adjmatrix_graph(graph_test1)
  n <- 100
  suppressWarnings(expression_matrix1 <- generate_expression_mat(n, adjacency_matrix1, cor = 0.8))
  expect_equal(nrow(expression_matrix1), length(V(graph_test1)))
  expect_equal(ncol(expression_matrix1), n)
  expect_equal(round(mean(expression_matrix1), 0), 0)
  expect_equal(round(sd(expression_matrix1), 0), 1)
  expect_true(is.matrix(expression_matrix1))
})


test_that("Generate Expression matrix from graph object", {
  graph_test1_edges <- rbind(c("A", "B"), c("B", "C"), c("B", "D"))
  graph_test1 <- graph.edgelist(graph_test1_edges, directed = TRUE)
  adjacency_matrix1 <- make_adjmatrix_graph(graph_test1)
  n <- 100
  suppressWarnings(expression_matrix1 <- generate_expression(n, graph_test1, cor = 0.8))
  expect_equal(nrow(expression_matrix1), length(V(graph_test1)))
  expect_equal(ncol(expression_matrix1), n)
  expect_equal(round(mean(expression_matrix1), 0), 0)
  expect_equal(round(sd(expression_matrix1), 0), 1)
  expect_true(is.matrix(expression_matrix1))
})

test_that("Generate Expression matrix from adjacency matrix by commonlinks", {
  graph_test1_edges <- rbind(c("A", "B"), c("B", "C"), c("B", "D"))
  graph_test1 <- graph.edgelist(graph_test1_edges, directed = TRUE)
  adjacency_matrix1 <- make_adjmatrix_graph(graph_test1)
  n <- 100
  suppressWarnings(expression_matrix1 <- generate_expression_mat(n, adjacency_matrix1, cor = 0.8, comm = TRUE))
  expect_equal(nrow(expression_matrix1), length(V(graph_test1)))
  expect_equal(ncol(expression_matrix1), n)
  expect_equal(round(mean(expression_matrix1), 0), 0)
  expect_equal(round(sd(expression_matrix1), 0), 1)
  expect_true(is.matrix(expression_matrix1))
})

test_that("Generate Expression matrix from graph object by commonlinks", {
  graph_test1_edges <- rbind(c("A", "B"), c("B", "C"), c("B", "D"))
  graph_test1 <- graph.edgelist(graph_test1_edges, directed = TRUE)
  adjacency_matrix1 <- make_adjmatrix_graph(graph_test1)
  n <- 100
  suppressWarnings(expression_matrix1 <- generate_expression(n, graph_test1, cor = 0.8, comm = TRUE))
  expect_equal(nrow(expression_matrix1), length(V(graph_test1)))
  expect_equal(ncol(expression_matrix1), n)
  expect_equal(round(mean(expression_matrix1), 0), 0)
  expect_equal(round(sd(expression_matrix1), 0), 1)
  expect_true(is.matrix(expression_matrix1))
})

test_that("Generate Expression matrix from adjacency matrix by relative distance", {
  graph_test1_edges <- rbind(c("A", "B"), c("B", "C"), c("B", "D"))
  graph_test1 <- graph.edgelist(graph_test1_edges, directed = TRUE)
  adjacency_matrix1 <- make_adjmatrix_graph(graph_test1)
  n <- 100
  suppressWarnings(expression_matrix1 <- generate_expression_mat(n, adjacency_matrix1, cor = 0.8, dist = TRUE))
  expect_equal(nrow(expression_matrix1), length(V(graph_test1)))
  expect_equal(ncol(expression_matrix1), n)
  expect_equal(round(mean(expression_matrix1), 0), 0)
  expect_equal(round(sd(expression_matrix1), 0), 1)
  expect_true(is.matrix(expression_matrix1))
})

test_that("Generate Expression matrix from graph object by relative distance", {
  graph_test1_edges <- rbind(c("A", "B"), c("B", "C"), c("B", "D"))
  graph_test1 <- graph.edgelist(graph_test1_edges, directed = TRUE)
  adjacency_matrix1 <- make_adjmatrix_graph(graph_test1)
  n <- 100
  suppressWarnings(expression_matrix1 <- generate_expression(n, graph_test1, cor = 0.8, dist = TRUE))
  expect_equal(nrow(expression_matrix1), length(V(graph_test1)))
  expect_equal(ncol(expression_matrix1), n)
  expect_equal(round(mean(expression_matrix1), 0), 0)
  expect_equal(round(sd(expression_matrix1), 0), 1)
  expect_true(is.matrix(expression_matrix1))
})

test_that("Generate Expression matrix from adjacency matrix by absolute distance", {
  graph_test1_edges <- rbind(c("A", "B"), c("B", "C"), c("B", "D"))
  graph_test1 <- graph.edgelist(graph_test1_edges, directed = TRUE)
  adjacency_matrix1 <- make_adjmatrix_graph(graph_test1)
  n <- 100
  suppressWarnings(expression_matrix1 <- generate_expression_mat(n, adjacency_matrix1, cor = 0.8, dist = TRUE, absolute = TRUE))
  expect_equal(nrow(expression_matrix1), length(V(graph_test1)))
  expect_equal(ncol(expression_matrix1), n)
  expect_equal(round(mean(expression_matrix1), 0), 0)
  expect_equal(round(sd(expression_matrix1), 0), 1)
  expect_true(is.matrix(expression_matrix1))
})

test_that("Generate Expression matrix from graph object by absolute distance", {
  graph_test1_edges <- rbind(c("A", "B"), c("B", "C"), c("B", "D"))
  graph_test1 <- graph.edgelist(graph_test1_edges, directed = TRUE)
  adjacency_matrix1 <- make_adjmatrix_graph(graph_test1)
  n <- 100
  suppressWarnings(expression_matrix1 <- generate_expression(n, graph_test1, cor = 0.8, dist = TRUE, absolute = TRUE))
  expect_equal(nrow(expression_matrix1), length(V(graph_test1)))
  expect_equal(ncol(expression_matrix1), n)
  expect_equal(round(mean(expression_matrix1), 0), 0)
  expect_equal(round(sd(expression_matrix1), 0), 1)
  expect_true(is.matrix(expression_matrix1))
})
```
