## Supplementary material for "graphsim: An R package for simulating gene expression data from graph structures of biological pathways": graphsim 1.0.0 R package: test_plot.R

```
library("graphsim")
library("igraph")
library("vdiffr")
context("Plot Directed Graphs")

#generate fixed layout
graph_test1_edges <- rbind(c("A", "B"), c("B", "C"), c("B", "D"))
graph_test1 <- graph.edgelist(graph_test1_edges, directed = TRUE)
## note that it is not possible to pass a set.seed to layout
# coords <- layout_with_fr(graph_test1, niter=50, start.temp=sqrt(10)/10)
coords <- structure(c(0.697871079806861, -0.229514654942849, -1.43395965899018, 
                      0.0584109525510336, 1.04730615951541, 0.189948108158827, 0.556504819830214, 
                      -1.03768519984881), .Dim = c(4L, 2L))


test_that("Plot an undirected graph structure", {
  graph_test1_edges <- rbind(c("A", "B"), c("B", "C"), c("B", "D"))
  graph_test1 <- graph.edgelist(graph_test1_edges, directed = TRUE)
  graph_test1 <- as.undirected(graph_test1)
  set.seed = 9000
  expect_doppelganger("test plotting undirected with igraph", plot(graph_test1, layout = coords))
  set.seed = 9000
  expect_doppelganger("test plotting undirected with plot_directed", plot_directed(graph_test1, layout = coords))
})

test_that("Plot a directed graph structure", {
  graph_test1_edges <- rbind(c("A", "B"), c("B", "C"), c("B", "D"))
  graph_test1 <- graph.edgelist(graph_test1_edges, directed = TRUE)
  set.seed = 9000
  expect_doppelganger("test plotting directed with igraph", plot(graph_test1, layout = coords))
  set.seed = 9000
  expect_doppelganger("test plotting directed with plot_directed", plot_directed(graph_test1, layout = coords))
  set.seed = 9000
  expect_doppelganger("test plotting directed with plot_directedas state vector", plot_directed(graph_test1, layout = coords, state = c(1, 0, 1)))
})

test_that("Plot a graph structure with plotting parameters", {
  graph_test1_edges <- rbind(c("A", "B"), c("B", "C"), c("B", "D"))
  graph_test1 <- graph.edgelist(graph_test1_edges, directed = TRUE)
  set.seed = 9000
  expect_doppelganger("test plotting with plot_directed parameters", plot_directed(graph_test1, layout = coords, labels = 1:3, cex.arrow = 1.2, cex.node = 2, cex.label = 0.8, cex.main = 0.8, fill.node = "lightblue", col.arrow = c("blue", "palevioletred", "grey85"), border.node = "black", main = "test title", xlab = "here is my plot", ylab = "here is my axes", frame.plot = TRUE))
})

test_that("Plot an inhibiting graph structure", {
  graph_test1_edges <- rbind(c("A", "B"), c("B", "C"), c("B", "D"))
  graph_test1 <- graph.edgelist(graph_test1_edges, directed = TRUE)
  state_matrix1 <- make_state_matrix(graph_test1, state = "inhibiting")
  set.seed = 9000
  expect_doppelganger("test plotting inhibitions with plot_directed", plot_directed(graph_test1, layout = coords, state_matrix1))
  set.seed = 9000
  expect_doppelganger("test plotting inhibitions with plot_directed as state vector", plot_directed(graph_test1, layout = coords, state = c(1, -1, 1)))
})
```
