## Supplementary material for "graphsim: An R package for simulating gene expression data from graph structures of biological pathways": graphsim 1.0.0 R package: test_sigma.R

```
library("graphsim")
library("igraph")
context("Make Sigma Matrix")

test_that("Generate Sigma matrix from adjacency matrix", {
    graph_test1_edges <- rbind(c("A", "B"), c("B", "C"), c("B", "D"))
    graph_test1 <- graph.edgelist(graph_test1_edges, directed = TRUE)
    adjacency_matrix1 <- make_adjmatrix_graph(graph_test1)
    sigma_matrix1 <- make_sigma_mat_adjmat(adjacency_matrix1, cor = 0.8)
    expect_equal(isSymmetric(sigma_matrix1), TRUE)
    expect_true(all(diag(sigma_matrix1) == rep(1, nrow(adjacency_matrix1))))
    expect_equal(nrow(sigma_matrix1), length(V(graph_test1)))
    expect_equal(ncol(sigma_matrix1), length(V(graph_test1)))
    expect_equal(sum(sigma_matrix1), 8.8)
    expect_true(all(adjacency_matrix1 == cbind(c(0, 1, 0, 0), c(1, 0, 1, 1), c(0, 1, 0, 0), c(0, 1, 0, 0))))
    expect_true(all(sigma_matrix1 == cbind(c(1, 0.8, 0, 0), c(0.8, 1, 0.8, 0.8), c(0, 0.8, 1, 0), c(0, 0.8, 0, 1))))
    expect_true(all(sigma_matrix1 %in% c(0, 1, 0.8/2^c(0:100))))
  })

test_that("Generate Sigma matrix from distances derived from adjacency matrix", {
  graph_test1_edges <- rbind(c("A", "B"), c("B", "C"), c("C", "D"), c("A", "D"), c("D", "C"), c("A", "C"))
  graph_test1 <- graph.edgelist(graph_test1_edges, directed = TRUE)
  adjacency_matrix1 <- make_adjmatrix_graph(graph_test1)
  expect_error(make_sigma_mat_dist_adjmat(adjacency_matrix1, cor = 0.8, absolute = TRUE), "distance matrix must have diagonal of zero")
  diag(adjacency_matrix1) <- 1
  expect_error(make_sigma_mat_dist_adjmat(adjacency_matrix1, cor = 0.8, absolute = TRUE), "distance matrix expected, not adjacency matrix")
})

test_that("Generate Sigma matrix from commonlink matrix", {
  graph_test1_edges <- rbind(c("A", "B"), c("B", "C"), c("B", "D"))
  graph_test1 <- graph.edgelist(graph_test1_edges, directed = TRUE)
  adjacency_matrix1 <- make_adjmatrix_graph(graph_test1)
  common_link_matrix1 <- make_commonlink_adjmat(adjacency_matrix1)
  sigma_matrix1 <- make_sigma_mat_adjmat(common_link_matrix1, cor = 0.8)
  expect_equal(isSymmetric(sigma_matrix1), TRUE)
  expect_true(all(diag(sigma_matrix1) == rep(1, nrow(adjacency_matrix1))))
  expect_equal(nrow(sigma_matrix1), length(V(graph_test1)))
  expect_equal(ncol(sigma_matrix1), length(V(graph_test1)))
  expect_equal(sum(sigma_matrix1), 8.8)
  expect_true(all(adjacency_matrix1 == cbind(c(0, 1, 0, 0), c(1, 0, 1, 1), c(0, 1, 0, 0), c(0, 1, 0, 0))))
  expect_equal(all(is.matrix(common_link_matrix1)), TRUE)
  expect_true(all(common_link_matrix1 == cbind(c(1, 0, 1, 1), c(0, 3, 0, 0), c(1, 0, 1, 1), c(1, 0, 1, 1))))
  expect_true(all(sigma_matrix1 == cbind(c(1, 0, 0.8, 0.8), c(0, 1, 0, 0), c(0.8, 0, 1, 0.8), c(0.8, 0, 0.8, 1))))
  expect_true(all(sigma_matrix1 %in% c(0, 1, 0.8/2^c(0:100))))
})

test_that("Generate Sigma matrix from distances derived from commonlink matrix", {
  graph_test1_edges <- rbind(c("A", "B"), c("B", "C"), c("C", "D"), c("A", "D"), c("D", "C"), c("A", "C"))
  graph_test1 <- graph.edgelist(graph_test1_edges, directed = TRUE)
  adjacency_matrix1 <- make_adjmatrix_graph(graph_test1)
  common_link_matrix1 <- make_commonlink_adjmat(adjacency_matrix1)
  expect_error(make_sigma_mat_dist_adjmat(common_link_matrix1, cor = 0.8, absolute = TRUE), "distance matrix must have diagonal of zero")
  diag(common_link_matrix1) <- 1
  expect_error(make_sigma_mat_dist_adjmat(common_link_matrix1, cor = 0.8, absolute = TRUE), "distance matrix expected, not adjacency matrix")
})

test_that("Generate Sigma matrix from distance matrix", {
  graph_test1_edges <- rbind(c("A", "B"), c("B", "C"), c("B", "D"))
  graph_test1 <- graph.edgelist(graph_test1_edges, directed = TRUE)
  adjacency_matrix1 <- make_adjmatrix_graph(graph_test1)
  distance_matrix1 <- make_distance_adjmat(adjacency_matrix1)
  sigma_matrix1 <- make_sigma_mat_dist_adjmat(distance_matrix1, cor = 0.8)
  expect_equal(isSymmetric(sigma_matrix1), TRUE)
  expect_true(all(diag(sigma_matrix1) == rep(1, nrow(adjacency_matrix1))))
  expect_equal(nrow(sigma_matrix1), length(V(graph_test1)))
  expect_equal(ncol(sigma_matrix1), length(V(graph_test1)))
  expect_equal(sum(sigma_matrix1), 11.2)
  expect_true(all(adjacency_matrix1 == cbind(c(0, 1, 0, 0), c(1, 0, 1, 1), c(0, 1, 0, 0), c(0, 1, 0, 0))))
  expect_true(all(distance_matrix1 == 1/cbind(c(1, 2, 4, 4), c(2, 1, 2, 2), c(4, 2, 1, 4), c(4, 2, 4, 1))))
  expect_true(all(distance_matrix1 %in% c(0, 1/2^c(0:100))))
  expect_true(all(sigma_matrix1 == cbind(c(1, 0.8, 0.4, 0.4), c(0.8, 1, 0.8, 0.8), c(0.4, 0.8, 1, 0.4), c(0.4, 0.8, 0.4, 1))))
  expect_true(all(sigma_matrix1 %in% c(1, 0.8/2^c(0:100))))
})

## all with graph

test_that("Generate Sigma matrix  from graph structure", {
  graph_test1_edges <- rbind(c("A", "B"), c("B", "C"), c("B", "D"))
  graph_test1 <- graph.edgelist(graph_test1_edges, directed = TRUE)
  adjacency_matrix1 <- make_adjmatrix_graph(graph_test1)
  common_link_matrix1 <- make_commonlink_graph(graph_test1)
  distance_matrix1 <- make_distance_graph(graph_test1)
  sigma_matrix1 <- make_sigma_mat_graph(graph_test1, cor = 0.8)
  expect_equal(isSymmetric(sigma_matrix1), TRUE)
  expect_true(all(diag(sigma_matrix1) == rep(1, nrow(adjacency_matrix1))))
  expect_equal(nrow(sigma_matrix1), length(V(graph_test1)))
  expect_equal(ncol(sigma_matrix1), length(V(graph_test1)))
  expect_equal(sum(sigma_matrix1), 8.8)
  expect_true(all(adjacency_matrix1 == cbind(c(0, 1, 0, 0), c(1, 0, 1, 1), c(0, 1, 0, 0), c(0, 1, 0, 0))))
  expect_true(all(sigma_matrix1 == cbind(c(1, 0.8, 0, 0), c(0.8, 1, 0.8, 0.8), c(0, 0.8, 1, 0), c(0, 0.8, 0, 1))))
  expect_true(all(sigma_matrix1 %in% c(0, 1, 0.8/2^c(0:100))))
})


test_that("Generate Sigma matrix  from graph structure", {
  graph_test1_edges <- rbind(c("A", "B"), c("B", "C"), c("B", "D"))
  graph_test1 <- graph.edgelist(graph_test1_edges, directed = TRUE)
  adjacency_matrix1 <- make_adjmatrix_graph(graph_test1)
  common_link_matrix1 <- make_commonlink_graph(graph_test1)
  distance_matrix1 <- make_distance_graph(graph_test1)
  sigma_matrix1 <- make_sigma_mat_dist_graph(graph_test1, cor = 0.8)
  expect_equal(isSymmetric(sigma_matrix1), TRUE)
  expect_true(all(diag(sigma_matrix1) == rep(1, nrow(adjacency_matrix1))))
  expect_equal(nrow(sigma_matrix1), length(V(graph_test1)))
  expect_equal(ncol(sigma_matrix1), length(V(graph_test1)))
  expect_equal(sum(sigma_matrix1), 11.2)
  expect_true(all(adjacency_matrix1 == cbind(c(0, 1, 0, 0), c(1, 0, 1, 1), c(0, 1, 0, 0), c(0, 1, 0, 0))))
  expect_true(all(sigma_matrix1 == cbind(c(1, 0.8, 0.4, 0.4), c(0.8, 1, 0.8, 0.8), c(0.4, 0.8, 1, 0.4), c(0.4, 0.8, 0.4, 1))))
  expect_true(all(sigma_matrix1 %in% c(0, 1, 0.8/2^c(0:100))))
})

test_that("Generate Sigma matrix  from graph structure", {
  graph_test1_edges <- rbind(c("A", "B"), c("B", "C"), c("B", "D"))
  graph_test1 <- graph.edgelist(graph_test1_edges, directed = TRUE)
  adjacency_matrix1 <- make_adjmatrix_graph(graph_test1)
  common_link_matrix1 <- make_commonlink_graph(graph_test1)
  distance_matrix1 <- make_distance_graph(graph_test1)
  sigma_matrix1 <- make_sigma_mat_graph(graph_test1, cor = 0.8, comm = TRUE)
  expect_equal(isSymmetric(sigma_matrix1), TRUE)
  expect_true(all(diag(sigma_matrix1) == rep(1, nrow(adjacency_matrix1))))
  expect_equal(nrow(sigma_matrix1), length(V(graph_test1)))
  expect_equal(ncol(sigma_matrix1), length(V(graph_test1)))
  expect_equal(sum(sigma_matrix1), 8.8)
  expect_true(all(adjacency_matrix1 == cbind(c(0, 1, 0, 0), c(1, 0, 1, 1), c(0, 1, 0, 0), c(0, 1, 0, 0))))
  expect_true(all(sigma_matrix1 == cbind(c(1, 0, 0.8, 0.8), c(0, 1, 0, 0), c(0.8, 0, 1, 0.8), c(0.8, 0, 0.8, 1))))
  expect_true(all(sigma_matrix1 %in% c(0, 1, 0.8/2^c(0:100))))
})

test_that("Generate Sigma matrix from graph structure with sd passing scalar", {
  graph_test1_edges <- rbind(c("A", "B"), c("B", "C"), c("B", "D"))
  graph_test1 <- graph.edgelist(graph_test1_edges, directed = TRUE)
  adjacency_matrix1 <- make_adjmatrix_graph(graph_test1)
  common_link_matrix1 <- make_commonlink_graph(graph_test1)
  distance_matrix1 <- make_distance_graph(graph_test1)
  sigma_matrix1 <- make_sigma_mat_graph(graph_test1, cor = 0.8, sd = 2)
  expect_equal(isSymmetric(sigma_matrix1), TRUE)
  expect_true(all(diag(sigma_matrix1) == rep(4, nrow(adjacency_matrix1))))
  expect_equal(nrow(sigma_matrix1), length(V(graph_test1)))
  expect_equal(ncol(sigma_matrix1), length(V(graph_test1)))
  expect_equal(sum(sigma_matrix1), 35.2)
  expect_true(all(adjacency_matrix1 == cbind(c(0, 1, 0, 0), c(1, 0, 1, 1), c(0, 1, 0, 0), c(0, 1, 0, 0))))
  expect_true(all(sigma_matrix1 == 4*cbind(c(1, 0.8, 0, 0), c(0.8, 1, 0.8, 0.8), c(0, 0.8, 1, 0), c(0, 0.8, 0, 1))))
  expect_true(all(sigma_matrix1 %in% c(0, 4, 4*0.8/2^c(0:100))))
})

test_that("Generate Sigma matrix from graph structure with sd passing vector", {
  graph_test1_edges <- rbind(c("A", "B"), c("B", "C"), c("B", "D"))
  graph_test1 <- graph.edgelist(graph_test1_edges, directed = TRUE)
  adjacency_matrix1 <- make_adjmatrix_graph(graph_test1)
  common_link_matrix1 <- make_commonlink_graph(graph_test1)
  distance_matrix1 <- make_distance_graph(graph_test1)
  sigma_matrix1 <- make_sigma_mat_graph(graph_test1, cor = 0.8, sd = 1:4)
  expect_equal(isSymmetric(sigma_matrix1), TRUE)
  expect_true(all(diag(sigma_matrix1) == rep((1:4)^2, nrow(adjacency_matrix1))))
  expect_equal(nrow(sigma_matrix1), length(V(graph_test1)))
  expect_equal(ncol(sigma_matrix1), length(V(graph_test1)))
  expect_equal(sum(sigma_matrix1), 55.6)
  expect_true(all(adjacency_matrix1 == cbind(c(0, 1, 0, 0), c(1, 0, 1, 1), c(0, 1, 0, 0), c(0, 1, 0, 0))))
  expect_true(all(abs(sigma_matrix1 - cbind(c(1, 1.6, 0, 0), c(1.6, 4, 4.8, 6.4), c(0.0, 4.8, 9, 0.0), c(0.0, 6.4, 0.0, 16))) < 1e+16))
})
```
