## Supplementary material for "graphsim: An R package for simulating gene expression data from graph structures of biological pathways": graphsim 1.0.0 R package: test_state.R

```
library("graphsim")
library("igraph")
context("Make State Matrix")

test_that("Default to activating matrix", {
  graph_test1_edges <- rbind(c("A", "B"), c("B", "C"), c("B", "D"))
  graph_test1 <- graph.edgelist(graph_test1_edges, directed = TRUE)
  state_matrix1 <- make_state_matrix(graph_test1)
  expect_equal(isSymmetric(state_matrix1), TRUE)
  expect_equal(sum(diag(state_matrix1)), length(V(graph_test1)))
  expect_equal(nrow(state_matrix1), length(V(graph_test1)))
  expect_equal(ncol(state_matrix1), length(V(graph_test1)))
  expect_equal(sum(state_matrix1), length(V(graph_test1))**2)
  expect_equal(all(state_matrix1 > 0), TRUE)
  expect_equal(all(is.matrix(state_matrix1)), TRUE)
  expect_true(all(state_matrix1 == 1))
})

test_that("text input to activating matrix", {
  graph_test1_edges <- rbind(c("A", "B"), c("B", "C"), c("B", "D"))
  graph_test1 <- graph.edgelist(graph_test1_edges, directed = TRUE)
  state_matrix1 <- make_state_matrix(graph_test1, state = "activating")
  expect_equal(isSymmetric(state_matrix1), TRUE)
  expect_equal(sum(diag(state_matrix1)), length(V(graph_test1)))
  expect_equal(nrow(state_matrix1), length(V(graph_test1)))
  expect_equal(ncol(state_matrix1), length(V(graph_test1)))
  expect_equal(sum(state_matrix1), length(V(graph_test1))**2)
  expect_equal(all(state_matrix1 > 0), TRUE)
  expect_true(all(state_matrix1 == 1))
})

test_that("text input to activating matrix", {
  graph_test1_edges <- rbind(c("A", "B"), c("B", "C"), c("B", "D"))
  graph_test1 <- graph.edgelist(graph_test1_edges, directed = TRUE)
  state_matrix1 <- make_state_matrix(graph_test1, state = 0)
  expect_equal(isSymmetric(state_matrix1), TRUE)
  expect_equal(sum(diag(state_matrix1)), length(V(graph_test1)))
  expect_equal(nrow(state_matrix1), length(V(graph_test1)))
  expect_equal(ncol(state_matrix1), length(V(graph_test1)))
  expect_equal(sum(state_matrix1), length(V(graph_test1))**2)
  expect_equal(all(state_matrix1 > 0), TRUE)
  expect_true(all(state_matrix1 == 1))
  state_matrix1 <- make_state_matrix(graph_test1, state = 1)
  expect_equal(isSymmetric(state_matrix1), TRUE)
  expect_equal(sum(diag(state_matrix1)), length(V(graph_test1)))
  expect_equal(nrow(state_matrix1), length(V(graph_test1)))
  expect_equal(ncol(state_matrix1), length(V(graph_test1)))
  expect_equal(sum(state_matrix1), length(V(graph_test1))**2)
  expect_equal(all(state_matrix1 > 0), TRUE)
  expect_true(all(state_matrix1 == 1))
})

test_that("text input to inhibiting matrix", {
  graph_test1_edges <- rbind(c("A", "B"), c("B", "C"), c("B", "D"))
  graph_test1 <- graph.edgelist(graph_test1_edges, directed = TRUE)
  state_matrix1 <- make_state_matrix(graph_test1, state = "inhibiting")
  expect_equal(isSymmetric(state_matrix1), TRUE)
  expect_equal(sum(diag(state_matrix1)), length(V(graph_test1)))
  expect_equal(nrow(state_matrix1), length(V(graph_test1)))
  expect_equal(ncol(state_matrix1), length(V(graph_test1)))
  expect_equal(sum(state_matrix1), length(V(graph_test1)))
  expect_equal(length(V(graph_test1))+sum(state_matrix1 < 0)-sum(state_matrix1 > 0) == 0, TRUE)
  expect_equal(all(state_matrix1 > 0), FALSE)
  expect_true(all(state_matrix1 == 1 || state_matrix1 == -1))
  expect_true(all(diag(state_matrix1) == 1))
  # expect only adjacent nodes to be inhibiting
  expect_true(all(state_matrix1 == (as_adjacency_matrix(as.undirected(graph_test1), sparse = FALSE) - 0.5) / - 0.5))
})

test_that("text input to inhibiting matrix", {
  graph_test1_edges <- rbind(c("A", "B"), c("B", "C"), c("B", "D"))
  graph_test1 <- graph.edgelist(graph_test1_edges, directed = TRUE)
  state_matrix1 <- make_state_matrix(graph_test1, state = 2)
  expect_equal(isSymmetric(state_matrix1), TRUE)
  expect_equal(sum(diag(state_matrix1)), length(V(graph_test1)))
  expect_equal(nrow(state_matrix1), length(V(graph_test1)))
  expect_equal(ncol(state_matrix1), length(V(graph_test1)))
  expect_equal(sum(state_matrix1), length(V(graph_test1)))
  expect_equal(length(V(graph_test1))+sum(state_matrix1 < 0)-sum(state_matrix1 > 0) == 0, TRUE)
  expect_equal(all(state_matrix1 > 0), FALSE)
  expect_true(all(state_matrix1 == 1 || state_matrix1 == -1))
  expect_true(all(diag(state_matrix1) == 1))
  state_matrix1 <- make_state_matrix(graph_test1, state = -1)
  expect_equal(isSymmetric(state_matrix1), TRUE)
  expect_equal(sum(diag(state_matrix1)), length(V(graph_test1)))
  expect_equal(nrow(state_matrix1), length(V(graph_test1)))
  expect_equal(ncol(state_matrix1), length(V(graph_test1)))
  expect_equal(sum(state_matrix1), length(V(graph_test1)))
  expect_equal(length(V(graph_test1))+sum(state_matrix1 < 0)-sum(state_matrix1 > 0) == 0, TRUE)
  expect_equal(all(state_matrix1 > 0), FALSE)
  expect_true(all(state_matrix1 == 1 || state_matrix1 == -1))
  expect_true(all(diag(state_matrix1) == 1))
  # expect only adjacent nodes to be inhibiting
  expect_true(all(state_matrix1 == (as_adjacency_matrix(as.undirected(graph_test1), sparse = FALSE) - 0.5) / - 0.5))
})

test_that("text input to inhibiting matrix as vector edge attributes", {
  graph_test1_edges <- rbind(c("A", "B"), c("B", "C"), c("B", "D"))
  graph_test1 <- graph.edgelist(graph_test1_edges, directed = TRUE)
  # test passing state attribute
  E(graph_test1)$state <-  c(1, -1, 1)
  state_matrix1 <- make_state_matrix(graph_test1)
  expect_equal(isSymmetric(state_matrix1), TRUE)
  expect_equal(sum(diag(state_matrix1)), length(V(graph_test1)))
  expect_equal(nrow(state_matrix1), length(V(graph_test1)))
  expect_equal(ncol(state_matrix1), length(V(graph_test1)))
  expect_equal(sum(state_matrix1), length(V(graph_test1)))
  expect_equal(length(V(graph_test1))+sum(state_matrix1 < 0)-sum(state_matrix1 > 0) == 0, TRUE)
  expect_equal(all(state_matrix1 > 0), FALSE)
  expect_true(all(state_matrix1 == 1 || state_matrix1 == -1))
  expect_true(all(diag(state_matrix1) == 1))
})

test_that("text input to inhibiting matrix as vector args", {
  graph_test1_edges <- rbind(c("A", "B"), c("B", "C"), c("B", "D"))
  graph_test1 <- graph.edgelist(graph_test1_edges, directed = TRUE)
  # test passing args
  state_matrix1 <- make_state_matrix(graph_test1, state = c(1, -1, 1))
  expect_equal(isSymmetric(state_matrix1), TRUE)
  expect_equal(sum(diag(state_matrix1)), length(V(graph_test1)))
  expect_equal(nrow(state_matrix1), length(V(graph_test1)))
  expect_equal(ncol(state_matrix1), length(V(graph_test1)))
  expect_equal(sum(state_matrix1), length(V(graph_test1)))
  expect_equal(length(V(graph_test1))+sum(state_matrix1 < 0)-sum(state_matrix1 > 0) == 0, TRUE)
  expect_equal(all(state_matrix1 > 0), FALSE)
  expect_true(all(state_matrix1 == 1 || state_matrix1 == -1))
  expect_true(all(diag(state_matrix1) == 1))
  state_matrix1 <- make_state_matrix(graph_test1, state = -1)
  expect_equal(isSymmetric(state_matrix1), TRUE)
  expect_equal(sum(diag(state_matrix1)), length(V(graph_test1)))
  expect_equal(nrow(state_matrix1), length(V(graph_test1)))
  expect_equal(ncol(state_matrix1), length(V(graph_test1)))
  expect_equal(sum(state_matrix1), length(V(graph_test1)))
  expect_equal(length(V(graph_test1))+sum(state_matrix1 < 0)-sum(state_matrix1 > 0) == 0, TRUE)
  expect_equal(all(state_matrix1 > 0), FALSE)
  expect_true(all(state_matrix1 == 1 || state_matrix1 == -1))
  expect_true(all(diag(state_matrix1) == 1))
})

test_that("text input to inhibiting matrix as vector with loop", {
  graph_test1_edges <- rbind(c("A", "B"), c("B", "C"), c("B", "D"), c("A", "D"))
  graph_test1 <- graph.edgelist(graph_test1_edges, directed = TRUE)
  # test passing state attribute
  E(graph_test1)$state <-  c(1, -1, 1, 1)
  state_matrix1 <- make_state_matrix(graph_test1)
  expect_equal(isSymmetric(state_matrix1), TRUE)
  expect_equal(sum(diag(state_matrix1)), length(V(graph_test1)))
  expect_equal(nrow(state_matrix1), length(V(graph_test1)))
  expect_equal(ncol(state_matrix1), length(V(graph_test1)))
  expect_equal(sum(state_matrix1), length(V(graph_test1)))
  expect_equal(length(V(graph_test1))+sum(state_matrix1 < 0)-sum(state_matrix1 > 0) == 0, TRUE)
  expect_equal(all(state_matrix1 > 0), FALSE)
  expect_true(all(state_matrix1 == 1 || state_matrix1 == -1))
  expect_true(all(diag(state_matrix1) == 1))
})

test_that("text input to inhibiting matrix as vector with conflicting loop", {
  graph_test1_edges <- rbind(c("A", "B"), c("B", "C"), c("B", "D"), c("A", "D"))
  graph_test1 <- graph.edgelist(graph_test1_edges, directed = TRUE)
  # test passing state attribute
  E(graph_test1)$state <-  c(1, -1, 1, -1)
  state_matrix1 <- make_state_matrix(graph_test1)
  expect_equal(isSymmetric(state_matrix1), TRUE)
  expect_equal(sum(diag(state_matrix1)), length(V(graph_test1)))
  expect_equal(nrow(state_matrix1), length(V(graph_test1)))
  expect_equal(ncol(state_matrix1), length(V(graph_test1)))
  #not expected for conflict
  #expect_equal(sum(state_matrix1), length(V(graph_test1)))
  #not expected for conflict
  #expect_equal(length(V(graph_test1))+sum(state_matrix1 < 0)-sum(state_matrix1 > 0) == 0, TRUE)
  expect_equal(all(state_matrix1 > 0), FALSE)
  expect_true(all(state_matrix1 == 1 || state_matrix1 == -1))
  expect_true(all(diag(state_matrix1) == 1))
})

test_that("text input to inhibiting matrix as vector with disconnected graph", {
  graph_test1_edges <- rbind(c("A", "B"), c("B", "C"), c("B", "D"), c("A", "D"), c("E", "F"))
  graph_test1 <- graph.edgelist(graph_test1_edges, directed = TRUE)
  # test passing state attribute
  E(graph_test1)$state <-  c(1, -1, 1, 1, -1)
  state_matrix1 <- make_state_matrix(graph_test1)
  # test passing args
  state_matrix1 <- make_state_matrix(graph_test1, state = c(1, -1, 1))
  expect_equal(isSymmetric(state_matrix1), TRUE)
  expect_equal(sum(diag(state_matrix1)), length(V(graph_test1)))
  expect_equal(nrow(state_matrix1), length(V(graph_test1)))
  expect_equal(ncol(state_matrix1), length(V(graph_test1)))
  #not expected for disconnected graph
  #expect_equal(sum(state_matrix1), length(V(graph_test1)))
  #not expected for disconnected graph
  #expect_equal(length(V(graph_test1))+sum(state_matrix1 < 0)-sum(state_matrix1 > 0) == 0, TRUE)
  expect_equal(all(state_matrix1 > 0), FALSE)
  expect_true(all(state_matrix1 == 1 || state_matrix1 == -1))
  expect_true(all(diag(state_matrix1) == 1))
})
```
