## Supplementary figures and images for "graphsim: An R package for simulating gene expression data from graph structures of biological pathways"

### demo2.gif

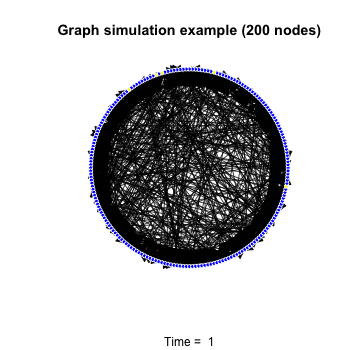

### demo.gif

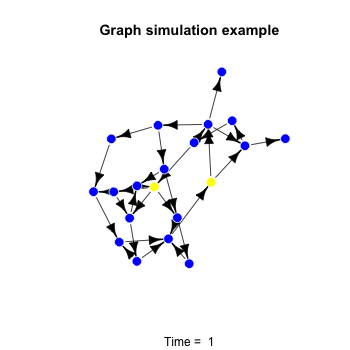
